# AnchorWave: sensitive alignment of genomes with high diversity, structural polymorphism and whole-genome duplication variation

**DOI:** 10.1101/2021.07.29.454331

**Authors:** Baoxing Song, Santiago Marco-Sola, Miquel Moreto, Lynn Johnson, Edward S. Buckler, Michelle C. Stitzer

## Abstract

Millions of species are currently being sequenced and their genomes are being compared. Many of them have more complex genomes than model systems and raised novel challenges for genome alignment. Widely used local alignment strategies often produce limited or incongruous results when applied to genomes with dispersed repeats, long indels, and highly diverse sequences. Moreover, alignment using many-to-many or reciprocal best hit approaches conflicts with well-studied patterns between species with different rounds of whole-genome duplication or polyploidy levels. Here we introduce AnchorWave, which performs whole-genome duplication informed collinear anchor identification between genomes and performs base-pair resolution global alignments for collinear blocks using the wavefront algorithm and a 2-piece affine gap cost strategy. This strategy enables AnchorWave to precisely identify multi-kilobase indels generated by transposable element (TE) presence/absence variants (PAVs). When aligning two maize genomes, AnchorWave successfully recalled 87% of previously reported TE PAVs between two maize lines. By contrast, other genome alignment tools showed almost zero power for TE PAV recall. AnchorWave precisely aligns up to three times more of the genome than the closest competitive approach, when comparing diverse genomes. Moreover, AnchorWave recalls transcription factor binding sites (TFBSs) at a rate of 1.05-74.85 fold higher than other tools, while with significantly lower false positive alignments. AnchorWave shows obvious improvement when applied to genomes with dispersed repeats, active transposable elements, high sequence diversity and whole-genome duplication variation.

**Significance statement:** One fundamental analysis needed to interpret genome assemblies is genome alignment. Yet, accurately aligning regulatory and transposon regions outside of genes remains challenging. We introduce AnchorWave, which implements a genome duplication informed longest path algorithm to identify collinear regions and performs base-pair resolved, end-to-end alignment for collinear blocks using an efficient 2-piece affine gap cost strategy. AnchorWave improves alignment of partially synthetic and real genomes under a number of scenarios: genomes with high similarity, large genomes with high TE activity, genomes with many inversions, and alignments between species with deeper evolutionary divergence and different whole-genome duplication histories. Potential use cases for the method include genome comparison for evolutionary analysis of non-genic sequences and population genetics of taxa with complex genomes.

## Introduction

Genome alignment tools are fundamental for comparative evolutionary analysis. Unlike initial genome sequencing efforts, which concentrated on cost-effective sequencing of model species, fulfilling the goal of sequencing a million eukaryotic reference genomes(1) adds many species with complex genomes(2). Aligning those genomes provides a revolutionary opportunity to understand biodiversity on Earth. Although the alignment of genic regions is bolstered by their modest length and conservation of amino acid residues, genes comprise only a minority of the nucleotides in a genome. Distal regulatory regions can be recalcitrant to alignment, often pushed far away from the genes they regulate by transposable element insertions(3). In addition, recursive whole-genome duplications result in fractionation of duplicated copies by deletion or pseudogenization(4) and chromosomal rearrangement further complicating genome alignments.

Seed-and-extend local alignment strategies(5, 6) have been widely successful for comparing model genomes. Such strategies generally trigger alignments with pairs of highly similar *k*-mers (seeds) from two genomes. When aligning genomes with many dispersed repeats, this strategy can trigger false alignments or even fail to generate an alignment when repetitive genome sequences are masked. The seed-and-extend strategy often fails when aligning regulatory elements with essential functions while diverse sequences. For example, the core motifs of transcription factor binding sites (TFBSs)(7–9) are 6.8 bp on average that is much shorter than the genome alignment seed size. Furthermore, the presence of highly diverse fragments can limit alignment extension(5) and confound local alignment. In addition, alignment using the affine gap cost strategy does not model mechanisms that generate indels of different length distributions(10), and alignment extension terminates in front of long indels (i.e., transposable element (TE) presence/absence variants (PAVs)). Finally, most genome alignment approaches generate many-to-many alignments or are limited to one-to-one alignments(11), which may not reflect the true evolutionary history when comparing taxa with ploidy level variation or unshared whole-genome duplications.

Here, we developed AnchorWave (Anchored Wavefront alignment), a whole-genome alignment method that could utilize well studied whole-genome duplication variation, perform sensitive sequence alignment with high accuracy and recall long indels. Those features are a significant improvement compared to current methods when aligning genomes with enriched dispersed repeats, high sequence diversity, high transposon activity or whole-genome duplication variation. Those complex genomic variations could be found in vertebrates(12) and are commonly in plant species(13).

## Results

AnchorWave leverages collinear regions to improve genome alignment (Fig. 1). Syntenic or collinear gene arrangements among taxa have been investigated by aligning protein-coding gene sequences(14). These collinear blocks are parsimoniously interpreted as being derived from a shared ancestor(14, 15). AnchorWave takes the reference genome sequence and gene annotation as input and extracts the reference full-length coding sequence (CDS) to use as anchors. We use minimap2(16) to lift over the position of the reference full-length CDS to the query genome (Fig. 1, step 1). AnchorWave then identifies collinear anchors using one of three user-specified algorithm options (Fig. 1, step 2) and uses the wavefront alignment (WFA) algorithm(17) to perform alignment for each anchor and inter-anchor interval (Fig. 1, step 4). Some anchor/inter-anchor regions cannot be aligned using our standard approach due to high computational costs. For these situations, AnchorWave either identifies novel anchors within long inter-anchor regions (Fig. 1, step 3) or, for those that cannot be split by novel anchors, aligns them using the ksw_extd2_sse function implemented in minimap2(16) or a reimplemented sliding window approach(18) (Fig. 1, step 4). AnchorWave concatenates base pair sequence alignment for each anchor and inter-anchor region and outputs the alignment in MAF format (Fig. 1, step 5).

**Fig. 1.**
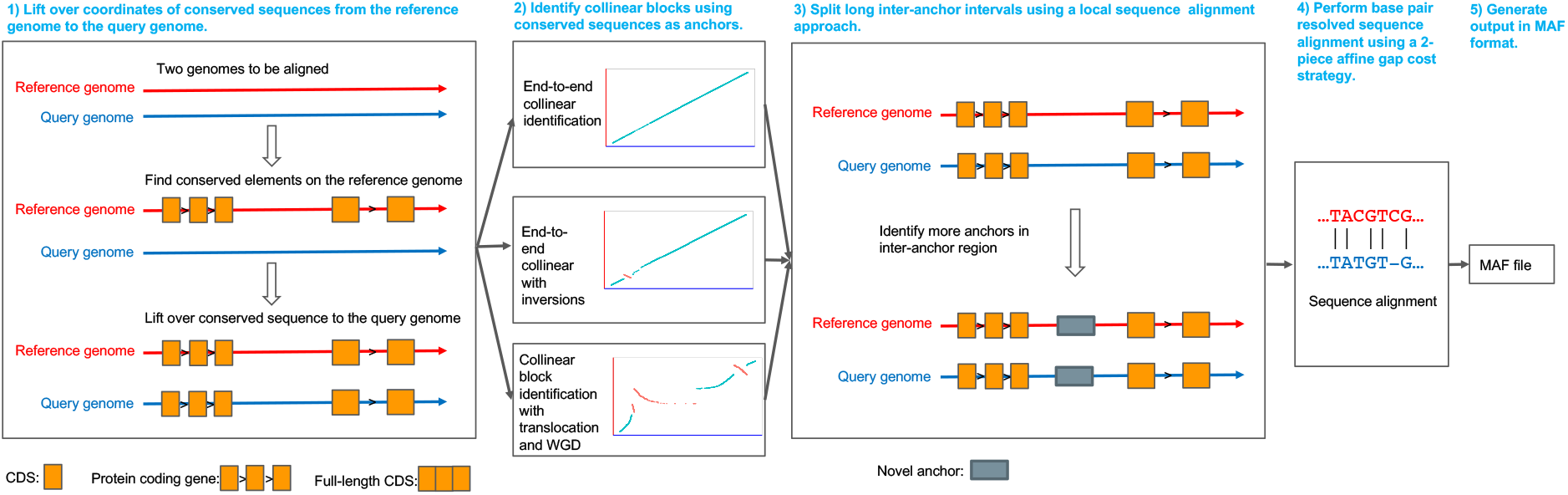
Principle of the AnchorWave process. AnchorWave identifies collinear regions via conserved anchors (here, full-length CDS) and breaks collinear regions into shorter fragments, i.e. anchor and inter-anchor intervals. By merging shorter intervals together after performing sensitive sequence alignment via a 2-piece affine gap cost strategy, AnchorWave generates a whole-genome alignment. AnchorWave implements commands to guide collinear block identification with or without chromosomal rearrangements and provides options to use known polyploidy levels or whole-genome duplications to inform the alignment.

To benchmark AnchorWave, we used partially synthetic and real genomes under a number of scenarios: small genomes with high similarity, large genomes with high TE activity, genomes with many inversions, and alignments between species with deeper evolutionary divergence and whole-genome duplication histories. We focused on the alignment performance in terms of TFBS sensitivity, TE alignment, and computational resources. To test AnchorWave for the alignment of highly similar genomes, we synthesized benchmark alignments by introducing variant calls of 18 Arabidopsis (*Arabidopsis thaliana*) accessions(19) to the TAIR10 reference genome. In these benchmark alignments, variants account for 2.13-3.60% of genome sites for each accession, including 1.22-2.38% of genome sites caused by variants longer than 50 bp (Fig. S1). We synthesized genomes with these variant calls and aligned them against the TAIR10 reference genome using AnchorWave, minimap2(16), LAST(11), MUMmer4(20), and GSAlign(21) and compared the newly generated alignments with benchmark alignments. AnchorWave was the only algorithm that aligns chromosomes end-to-end and aligned highly diverse fragments that were not aligned in the benchmark (Fig. S2), leading to a slight decrease in precision, with only minimap2 ranking higher. While AnchorWave had the highest F-score and recall (Fig. 2A, Fig. S3, and Supplementary Note 1).

**Fig. 2.**
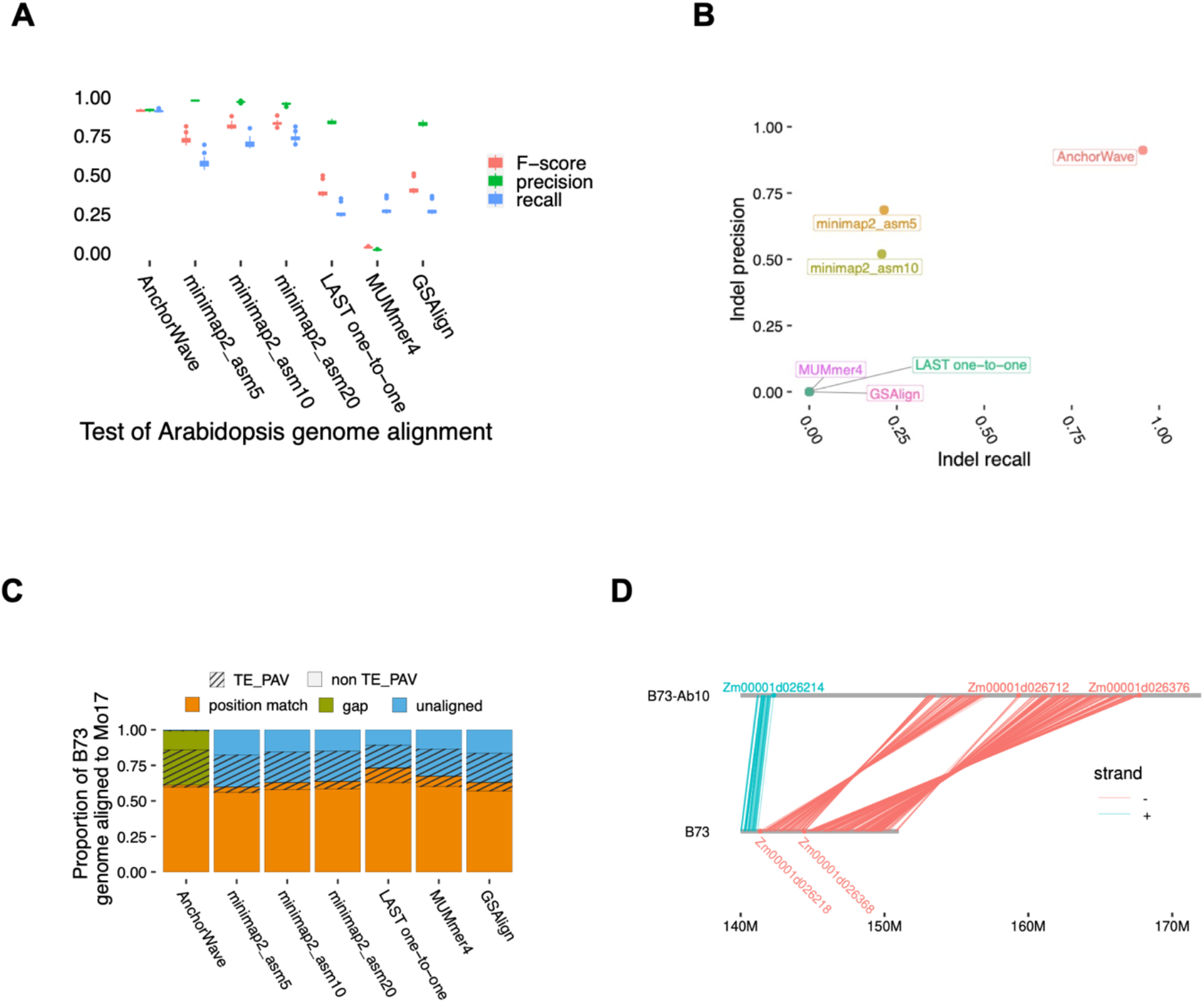
Comparison of genome alignment tools using genomes of different individuals in the same species. (**A**) Comparison of the performance of genome alignment tools at variant sites of 18 Arabidopsis accession alignment benchmarks. Genome alignments were performed using minimap2 with presettings asm5, asm10, and asm2 which terminate extension in regions with 5%, 10%, and 20% sequence divergence respectively. GSAlign and MUMmer4 alignments were performed with default parameters. LAST genome alignments with default parameters were labeled as LAST many-to-many. LAST many-to-many alignments were processed with a chain and net procedure to generate LAST many-to-one alignments (each query genome nucleotide may be aligned multiple times, while each reference nucleotide can be aligned up to one time). LAST many-to-one alignments were filtered to generate LAST one-to-one alignments (Materials and Methods, and Supplementary Note 1). (**B**) Recall and precision of TE deletions by aligning the TE-removed maize B73 genome against the reference genome. MUMmer4, GSAlign and LAST one-to-one had zero recall ratio. (**C**) Overview of the maize B73 genome sites aligned to the maize Mo17 genome. In those TE variation regions, which were previously reported as present in B73 and absent in Mo17 (TE on the legend), no position match alignments are expected. A higher number of position matches in these regions (striped orange) indicates a higher false-positive ratio. (**D**) Two inversions located using AnchorWave between the maize B73-Ab10 assembly and the B73 v4 reference genome.

To evaluate the performance for detecting long indels in repeat-rich genomes, we developed a benchmark by removing ~60% of LTR retrotransposons from the maize (*Zea mays* L.) B73 v4 assembly(22). This synthetic TE-removed genome had 84,271 deletions with lengths ranging from 1,144 to 33,730 bp (Fig. S4). We counted the number of TE deletions that could be correctly recalled by aligning the TE-removed genome against the reference genome (Supplementary Note 2). The genome alignment results from GSAlign, LAST, and MUMmer4 did not generate any variant longer than 1 Kbp. Minimap2(16) recalled ~21% of these long deletions correctly, likely benefiting from the usage of global alignment between adjacent anchors in a chain. AnchorWave recalled ~95% of these deletions correctly (Fig. 2B).

To evaluate the performance of the alignment approaches using a realistic polymorphism landscape, where long indels are mixed together with SNPs and short indels, we aligned the genomes of two maize individuals (B73 v4 and Mo17(23), Supplementary Note 3). AnchorWave aligned 61% of the B73 genome as position match (defined as an ungapped alignment, either matched or mismatched nucleotides, Fig. S5), which is comparable with other tools (Fig. 2C). However, AnchorWave produced the lowest number of position matches in previously reported B73 TE absent regions(24), suggesting that AnchorWave generated the fewest false positives among compared tools (Fig. 2C). Moreover, AnchorWave aligned much (37%) of the genome as deletions (gaps). Such gaps are expected for alignments between maize individuals, where indel variation arises from TE PAVs. Previous approaches have identified precise boundaries arising from transposition for 15,182 TEs that are present in one genotype and absent in another(24). AnchorWave increased this number to 28,321, recovering 87% (13,181) of these site-defined TE PAVs from Anderson et al. 2019 (24). Other tools had almost zero recall ratios of TE PAVs (Table 1), as they generated few gapped alignments (Fig. 2C).

**Table 1:**
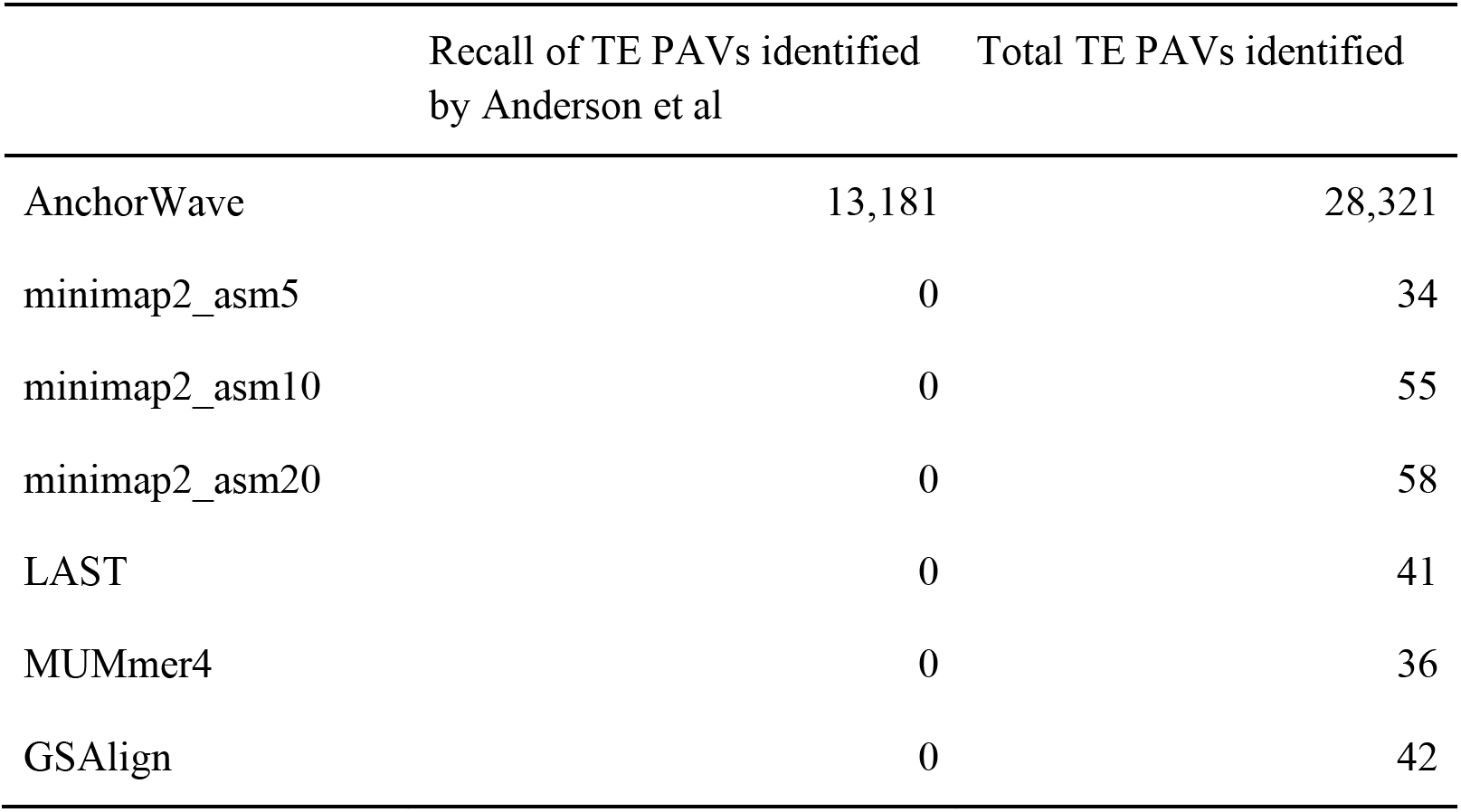
TE PAVs identified using different genome alignment tools. There are 341,426 TEs with complete target site duplication (TSD) sequences in the B73 genome and 305,022 TEs in the Mo17 genome. 15,182 TEs were identified as wholly absent in either B73 or Mo17 by Anderson et al. 2019 (24) (there, termed “site-defined” TEs).

The frequent presence of inversions in eukaryotic genomes poses another obstacle to the end-to-end alignment of chromosomes. By incorporating anchor strand information into our collinear identification approach, AnchorWave efficiently identifies inversions. As an example, we show gene-level resolution of two neighboring inversions of Abnormal chromosome 10 of maize, a cytologically known inversion that carries a meiotic drive locus(25) (Fig. 2D, Fig. S6 and Dataset S1). Other recall cases of previously reported inversions using AnchorWave are included in Supplementary Note 4 and Dataset S2.

Unlike other alignment approaches, AnchorWave can use knowledge about whole-genome duplications to guide whole-genome alignment. We leveraged this feature to align the genomes of goldfish (*Carassius auratus*)(26) and zebrafish (*Danio rerio*)(27), which represents a young (~14 Mya) vertebrate whole-genome duplication (Supplementary Note 5). Goldfish have twice as many chromosomes as zebrafish, but the size of their genome assemblies are similar (~1.3 Gbp), suggesting that extensive DNA PAVs. Using parameters that allowed each zebrafish anchor to define up to two collinear blocks in goldfish, AnchorWave aligned ~80% of the zebrafish genome sequence, facilitating inference of the evolutionary consequences of sequence PAVs.

The alignment quality of extant sequences is notoriously difficult to assess. Here we used biologically informed expectations about sequence conservation between maize and sorghum (*Sorghum bicolor*) to compare genome alignment approaches. The maize lineage has been under a whole-genome duplication since its divergence with sorghum (*Sorghum bicolor*)(28), while the subsequent chromosomal fusion caused these species to have the same chromosome number (n = 10). Due to independent TE movement and diversification along the maize and sorghum lineages, we expect very little sharing of individual TE copies; 70%-85%(29, 30) of the maize genome is composed of structurally recognizable TEs with estimated insertion times more recent than the divergence from sorghum (Fig. 3A). Allowing up to two collinear paths for each sorghum CDS (Fig. 3B, Fig. S7, Supplementary Note 6), AnchorWave aligned a significantly larger proportion of the maize genome compared to other tools (3.4-times that of the second-highest, generated via LAST many-to-many) (Fig. 3C). TFBSs are functional elements that are under purifying selection and are expected to be less affected by absence variants than other sequences(31). In TFBS(7) regions of the maize genome, AnchorWave genome alignment showed the highest recall, as well as a higher match ratio compared to the rest of the genome (Fig. 3D), suggesting it aligns these functional sequences more comprehensively.

**Fig. 3.**
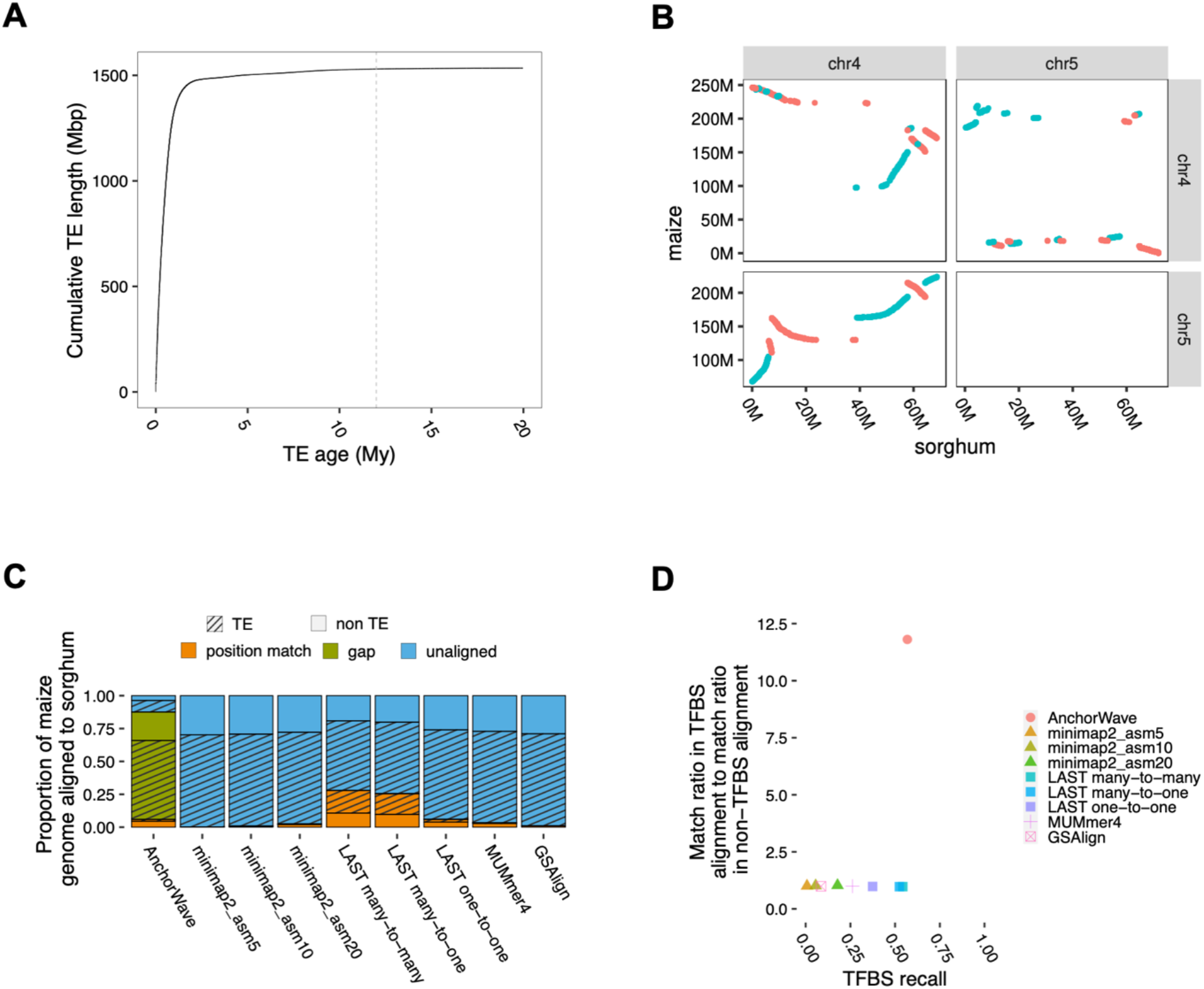
A comparison of different genome alignment tools for aligning maize B73 v4 assembly and sorghum genome. (**A**) Cumulative distribution of TE length v.s. TE age in the B73v4 maize genome, with age measured in millions of years (My). The dashed line indicates 12 My, the estimated divergence time of maize and sorghum(28). TE age data are from Stitzer et al., 2019(29); 371 TEs older than 20 My were not plotted, the total length of these 371 TEs is 531 Kbp. (**B**) The identified collinear anchors between the maize B73 v4 assembly and sorghum genome on chr4 and chr5. Each dot was plotted based on the start coordinate on the reference genome and query genome of an anchor. Collinear anchors on the same strand between the reference genome and query genome are shown in blue, otherwise red. (**C**) Sequence alignment between the maize B73 v4 genome and the sorghum genome. Minimap2, MUMmer4, GSAlign generated many-to-many alignments. Since most maize TEs are not shared with sorghum, a higher number of position matches in maize TE regions (striped orange) indicates a higher false-positive ratio. AnchorWave aligns 87.6% of the maize genome to the sorghum genome, while the second highest is 28.0% generated by LAST many-to-many. (**D**) Comparison of the proportion of maize TFBS that were aligned as a position match (recall) and the match ratios (number of match sites to number of aligned sites) in TFBS v.s. non-TFBS region.

AnchorWave exploits the latest advances in sequence alignment using the wavefront algorithm(17), which reduces the overall memory and computational requirements for global pairwise alignment. To limit memory usage for long sequence comparisons, AnchorWave implements two strategies: identifying novel anchors within long inter-anchor regions, and for inter-anchor regions that lack sufficient homology, approximating alignment using either a banded approach or sliding windows. In this study, we tuned sliding windows and novel anchor parameters to make full use of the available computer memory (Materials and Methods), using consistent parameters for different genome comparisons. Although the execution time of AnchorWave is high for some experiments, it remains comparable when considering that most other methods fail to align many bases in the genome (Table S1–S3).

## Discussion

Genome evolution across the tree of life has resulted in species with vastly differentiated ploidy, chromosome number, and genome organization. Beyond genome structure, species differ in the complexity and magnitude of non-genic repetitive sequences. Despite this widespread variation, genomes are always punctuated by evolutionarily conserved regulatory sequences and genes. AnchorWave makes use of this conservation, utilizing gene collinearity to guide genome alignment. This approach does not need highly similar seeds to trigger alignment in diverse regions and increases the sequence alignment sensitivity. AnchorWave uses collinear anchors to guide the alignment of repeat elements, and the employment of the 2-piece affine gap cost strategy enables the alignment of long indels. Chromosomal fusions after whole genome duplication can complicate the separation of subgenomes, but the whole genome duplication informed collinear blocks identification function in AnchorWave improved the alignment for genomes with whole genome duplication variation.

We highlight AnchorWave’s ability to align maize and sorghum (Fig. 3), as these species differ by many difficult issues for genome alignment. Maize is an allopolyploid, merging two diploid species with n=10 (28, 32), extensive fractionationation of subgenomes and chromosome fusions have occurred to generate the extant maize genome (33). Thousands of maize genes have remained duplicated relative to sorghum (33), but without an explicit model of whole genome duplication most genome alignment strategies fail to align these regions. By generating alignment paths in each subgenome, AnchorWave allows interpretation of the conservation and evolution of these genes and their local regulatory regions. As much of this regulatory sequence is embedded between repetitive TEs, AnchorWave reduces false positive alignments in this repetitive space.

While AnchorWave provides improved alignments for many complicated but real issues in genomics, it has some limitations. AnchorWave relies on the existence of collinear regions to guide alignment, and the extent of collinearity decays with phylogenetic distance. In addition, fractionation after whole-genome duplication can break collinear blocks (Supplementary Note 7). In comparisons between the human and mouse genomes, gene collinearity is limited. AnchorWave still provided alignment over regulatory DNA (Supplementary Note 8), showing value even for comparisons without extensive collinearity. At an extreme, although the autosomes of human and chimpanzee share largely collinear genes, the gene-depauperate Y chromosome could not be aligned (Supplementary Note 9). Finally, technical limitations such as fragmented genome assemblies can prevent the identification of collinearity, although this will likely pose less of a problem with advances in long-read sequencing.

When comparing genomes with different rounds of whole-genome duplication, AnchorWave significantly increased the proportion of the genome that was aligned compared to one-to-one alignments and reduced false-positive alignments compared to many-to-many alignments (Fig. 3C). Compared to alignment approaches using a seed-and-extension strategy, AnchorWave increased sensitivity for regulatory sequences and could recall long indel variants. AnchorWave’s collinear approach further reduces false-positive alignments from dispersed repeats. We showed that AnchorWave can generate whole-genome alignments, facilitating studies of the evolution of regulatory elements, TE polymorphisms, and chromosomal rearrangements such as inversions. AnchorWave even allows duplicated collinear blocks to be aligned, making it particularly relevant to plants, where an estimated 35% of species are recent polyploids(13).

## Materials and Methods

### Collinear anchor identification

AnchorWave takes the reference genome in FASTA format, the reference genome annotation in GFF(3) format, and the query genome in FASTA format as input. AnchorWave extracts the full-length CDS from the reference genome using the reference genome annotation. The positions of the reference full-length CDS to the query genome are lifted over using minimap2. AnchorWave then implements a longest-path dynamic programming algorithm to identify collinear anchors. Base pair resolution sequence alignments within each anchor and inter-anchor region are conducted using the 2-piece affine gap cost strategy, and these alignments are concatenated together to generate the alignment for the collinear block.

A longest path dynamic programming approach is applied to a pair of chromosomes. On the reference chromosome is a list of ***n*** anchors:

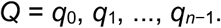

On the query chromosome is a list of ***m*** lift over hits:

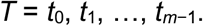

For each anchor and its hit, ***q*** and ***t***, the start and end positions, respectively, are identified from the SAM file output from minimap2. We set up a list of ***o*** anchor matches:

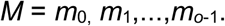

Individual matches are defined as:

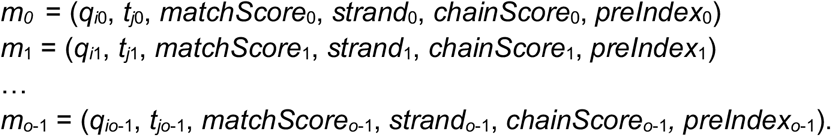

Where ***t_jk_*** is the lift over hit of ***q_ik_*** identified via minimap2, 0 ≤ *k* < o, 0 ≤ *i* < *n*, 0 ≤ *j* < *m*. ***matchScore_k_*** is the sequence similarity (ratio of the number of identical nucleotides to the length of the reference full-length CDS). If ***q_ik_*** and ***t_jk_*** are on the same strand, ***strand_k_*** is set as positive; otherwise, it is set as negative. ***chainScore_k_*** is initialized with ***matchScore_k_***. ***preIndex_k_*** is the index of the previous anchor on the chain and is initialized as −1, meaning that the current anchor is the first one on a chain.

Various longest path approaches have been developed for genomes with different types of chromosomal rearrangements. The goal is to guide the alignments using prior knowledge. When a user has limited background knowledge on the species being aligned, the third approach can be used as the default choice.

1. **Longest path approach for genome sequences without inversions or rearrangements** The target is to select a subset of positive-strand, non-overlapping anchor matches from ***M*** that give a maximum value of ***chainScore***, where the positions of anchors on the reference and query genomes increase. To do this, the matches in ***M*** are first sorted in ascending order by reference anchor start positions. Then, with 0 ≤ *e* < *f* ≤ *o*, for a previous element, *m*_e_ = (*q_ie_, t_je_, matchScore*_e_, *strand*_e_, *chainScore*_e_, *preIndex*_e_) and for a current element *m*_f_ = (*q_if_, t_je_, matchScore*_f_, *strand*_f_, *chainScore*_f_, *preIndex*_f_). The list of matches is iterated, incrementing ***e*** and ***f***, while the following conditions are true:

1. The strands of ***m*_f_** and ***m*_e_** are positive.
2. The end position of reference anchor ***q_ie_*** is smaller than the start position of reference anchor ***q_if_***.
3. The end position of query anchor *t_ie_* is smaller than the start position of query anchor ***t_if_***.
4. For each ***m_e_*** and ***m_f_*** pair, update the ***m_f_***’s ***chainScore*** based on the following:

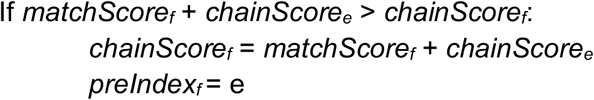 Starting from the match that has the maximum ***chainScore***, AnchorWave will use the ***preIndex*** values to track back and produce a list of matches that give the highest score.
2. **Longest path considering inversions** After sorting the matches in ascending order based on the reference anchor start positions, to consider inversions, we create ***currentScore*** which is a cumulative score, and ***maxScore*** which is the maximum value that ***currentScore*** has ever reached for each round of iteration. When a match is encountered on the negative strand, we assign its ***matchScore*** to ***currentScore***. If the next match is on the negative strand and the query start position is smaller than the query start position of the current one, we add this ***matchScore*** to the ***currentScore***. Otherwise, we subtract this ***matchScore*** from the ***currentScore***. When the ***currentScore*** drops below 0, the iteration is terminated and the next iteration starts. If the maximum cumulative score (***maxScore***) is larger than a preset threshold, for all matches in a kept group, we reverse their order in the list of matches. Those reversed matches have increasing query start positions and decreasing reference positions and thus remain on the diagonal. A similar longest path dynamic programming algorithm is applied to find an end-to-end chain as described above, except that the end position of anchor ***q_if_*** is smaller than the start position of anchor ***q_ie_*** on the reference chromosome when ***strand_e_*** and ***strand_f_*** are negative. With 0≤ *e* < *f* ≤ *o*, for a previous element, *m*_e_ = (*q_ie_, t_je_, matchScore*_e_, *strand*_e_, *chainScore*_e_, *preIndex*_e_) and a current element *m*_f_ = (*q_if_, t_jf_, matchScore*_f_, *strand*_f_, *chainScore_f_, preIndex_f_*), iterate the list of matches, incrementing ***e*** and ***f***, while the following conditions are true:

1. If the current match and previous match are on the negative strand, the end position of reference anchor ***q_if_*** is smaller than the start position of reference anchor ***q_ie_***. Otherwise, the end position of reference anchor ***q_ie_*** is smaller than the start position of reference anchor ***q_if_***.
2. The end position of query anchor ***t_ie_*** is smaller than the start position of query anchor ***t_if_***.
3. For each ***m_e_*** and ***m_f_*** pair, update the ***m_f_***’s chainScore based on the following:

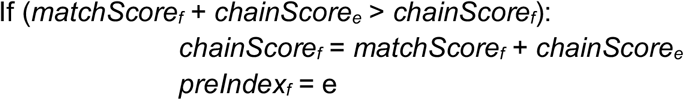 Starting from the match that has the maximum ***chainScore***, AnchorWave will use the ***preindex*** values to track back and output the list of matches that give the highest score.
3. **Longest path considering inversions, rearrangements, and whole-genome duplications** AnchorWave implements a function to constrain the alignment depths for both the reference genome and query genome, which is useful when there may be multiple collinear paths, i.e., genomes with chromosomal translocations, chromosome fusion, and varying numbers of rounds of whole-genome duplications. To do this, the matches are sorted in ascending order based on the reference anchor start positions. Then, with 0 ≤ *e* < *f* ≤ *o*, for a previous element, *m*_e_=(*q_ie_, t_je_, matchScore*_e_, *strand*_e_, *chainScore*_e_) and a current element, *m*_f_=(*q_if_, t_jf_, matchScore*_f_, *strand*_f_, *chainScore*_f_) iterate the list of matches, incrementing ***e*** and ***f***, while the following conditions are true:

1. The current match and previous match are on the same strand.
2. The end position of anchor ***q_ie_*** is smaller than the start position of anchor ***q_if_*** on the reference chromosome.
3. If ***strand_e_*** is positive, the end position of anchor ***t_ie_*** is smaller than the start position of anchor ***t_if_*** on the query chromosome. If ***strand_e_*** is negative, the end position of anchor ***t_ie_*** is smaller than the start position of anchor ***t_if_*** on the query chromosome.
4. For each ***m_e_*** and ***m_f_*** pair, update the ***m_f_***’s ***chainScore*** based on the following:

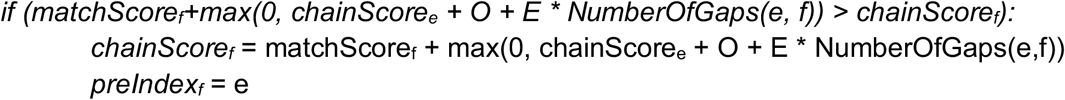 ***O*** is a gap opening penalty and ***E*** is a gap extension penalty, Let ***a*** be the number of anchors between anchor ***q_ie_*** and ***q_if_***, and let ***b*** be the number of anchors between ***t_je_*** and ***t_jf_***.

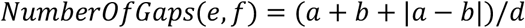

***d*** is a settable parameter(by default: 3). AnchorWave selects the chain with the maximum ***chainScore***. If the maximum ***chainScore*** is larger than a settable threshold, then output the chain. All the reference and query anchors that fall into the chain range will be counted. Matches with anchors that fall into chain ranges that are counted as larger than a settable threshold are marked and will not be used for the next round of iteration. Then, the next iteration starts using all the matches not being marked, until the maximum ***chainScore*** is smaller than a settable threshold.

### Filtering anchors to improve alignment quality

Correcting lift over full-length CDSs from the reference genome to the query genome is central in the AnchorWave pipeline. We use minimap2 to lift over full-length CDSs. Minimap2 misaligns small exons (https://github.com/lh3/minimap2#limitations). To reduce this side effect, when extracting full-length CDSs, AnchorWave ignores CDS exon records <20 bp (although this limit is a user-settable parameter). If the full-length CDSs of two or more genes are identical, AnchorWave ignores all of them in subsequent analysis.

We first identify full-length CDSs that might produce incorrect lift overs. We use minimap2 to map extracted full-length CDSs to the reference genome and rank the hits of each full-length CDS based on similarity (ratio of the number of identical DNA base pairs to the length of the reference full-length CDS). To remove tandem duplications, AnchorWave uses two thresholds: a number of mapping hits threshold (***e***, 1 by default) and a similarity ratio threshold (0.6 by default). Any full-length CDS with the number of mapping hits > ***e*** on a single reference chromosome sequence is further investigated using the similarity ratio threshold. For the sorted hits of this full-length CDS, if the ratio of ***e + 1*** hit similarity to the highest hit similarity is above the similarity ratio threshold, we drop this full-length CDS and its hits on any chromosome. We then use the longest path approach for genome sequences without inversions or rearrangements as described above to further filter the hits. We compare the coordinates of left hits with the original GFF(3) file; any full-length CDS with different coordinates between the original GFF(3) file and lift over hits would be placed on an unwanted list.

We then use minimap2 to lift over extracted full-length CDSs to the query genome. The fulllength CDSs in the unwanted list are not used. Anchors are further filtered to reduce the impact of tandem duplications using the same approach and parameters as described above. After filtering, the remaining anchors are fed into a user-specified longest path algorithm to identify collinear blocks.

### Identifying additional anchors to reduce the size of inter-anchor intervals

To improve sequence alignment quality and computational efficiency, additional anchors are needed in long inter-anchor intervals. AnchorWave used the minimap2 library to perform local alignment in collinear inter-anchor regions. In each inter-anchor region, AnchorWave selects the minimap2 primary alignment and specifies this as a novel anchor. This step is iterated until all inter-anchor intervals are shorter than a settable threshold, the sequence similarity of the new primary alignment is lower than a threshold (by default, we do not filter novel anchors using similarity and this is set as 0), or no new minimap2 matches can be found.

### Base pair resolution sequence alignment using the 2-piece affine gap cost strategy

Based on the assumption that pairs of sequences in each anchor or inter-anchor region are passed down from a common ancestor, AnchorWave performs base-pair resolution global sequence alignment for each anchor and inter-anchor interval using the 2-piece affine gap cost strategy(16). We implemented the 2-piece affine gap cost strategy in the WFA(17) library, and sequence alignments are conducted using WFA by default. If the WFA library requires more memory than a preset threshold, the “ksw_extd2_sse” functions implemented in minimap2 are called using a calculated bandwidth. The memory cost threshold of the WFA library is calculated using the “-w” parameter of AnchorWave. The bandwidth of minimap2 is calculated using the “-w” parameter of AnchorWave and anchor/inter-anchor sequence length. For longer sequences with calculated minimap2 bandwidth smaller than “-w”, we implement the 2-piece affine gap cost strategy with a sliding window approach(18), which generates approximate sequence alignments. The sliding window size (-w) was set as 38,000 in this study, which was also used as the minimum bandwidth for minimap2.

Long indels are generally derived from TE movements, while different mechanisms introduce shorter indel mutations(10). Since most available genome alignment tools align long indels and short indels using the same gap penalty profile, long indels generally fail to be aligned. Here, we implemented the 2-piece gap cost strategy to align long indels following equation 4 described by Li (16). Let the reference ***r = r_0_, r_1_… r_n−1_*** and the query ***q = q_0_, q_1_… q_m−1_*** be a pair of sequence fragments in an anchor or inter-anchor interval from the reference genome and the query genome, respectively, with length |***r***| = ***n*** and |***q***| = ***m***. ***O_1_*** is the first piece affine gap open penalty, ***E_1_*** is the first piece affine extend open penalty, ***O_2_*** is the second piece affine gap open penalty, and ***E_2_*** is the second piece affine extend open penalty. Let ***l*** be the gap length, min{***O_1_*** + |***l***| · ***E_1_, O_2_*** + |***l***| · ***E_2_***} was used as the indel penalty for the dynamic programming sequence alignment approach. We always assume ***O_1_*** + ***E_1_*** < ***O_2_*** + ***E_2_***. On the condition that ***E_1_*** > ***E_2_***, it applies cost ***O_1_*** + |***l***| · ***E_1_*** to gaps shorter than ⌊(***O_2_*** – ***O_1_***) / (***E_1_*** – ***E_2_***)⌋ and applies ***O_2_*** + |***l***| · ***E_2_*** to longer gaps.

The values of the first piece affine gap cost and second piece affine gap cost were selected based on the finding that TE copies are longer than 50 bp(29) (Fig. S8).

### Data sources and methods for validation

The released variant calling results for 18 Arabidopsis individuals were downloaded in sdi format from http://mtweb.cs.ucl.ac.uk/mus/www/19genomes/variants.SDI/. Taking the TAIR10 genome as a reference, we generated synthetic genome sequences using the “pseudogeno” command of GEAN(18) and benchmark genome alignments by replacing the reference alleles with alternative alleles for each accession separately.

To compare the newly generated genome alignments with benchmark genome alignments, we first counted the number of positions in the alignment that differed between the reference and alternative genome (referred to as “variant sites”, Fig. S5). True positive variant sites in the alignment are reference sites that are shared at the same query genome position in both the benchmark and newly generated alignment. False-positive variant sites are those present in the newly generated alignment but not in the benchmark alignment. False-negative variant sites are those present in the benchmark alignment but not in the newly generated alignment.

We calculate precision as:

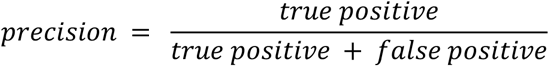

We calculate recall as:

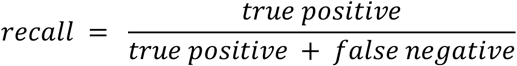

The harmonic mean of precision and recall is used to calculate the balanced F-score:

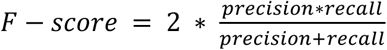

Most sites in these alignments are invariant, so we also calculate genome-wide precision and genome-wide recall in addition to those described above for variant sites. We calculate precision for genome-wide sites as the ratio of the number of aligned sites shared between the benchmark alignment and the newly generated alignment to the number of sites in the newly generated alignment. Conversely, the ratio of the number of shared aligned sites to the number of sites in the benchmark is defined as the genome-wide recall. We calculate the F-score for genome-wide sites using genome-wide precision and recall with the formula shown above.

The movement of TEs to new positions in the genome generates repeated interspersed sequences, which can cause false-positive alignments. TE presence-absence variants terminate the extension of sequence alignment and reduce the proportion of the genome that is aligned. We removed ~60% of long terminal repeat (LTR) retrotransposons from the maize B73 v4 genome, as these TEs make up the majority of genomic DNA. We used results from LTRharvest (34) with the parameters “-motif tgca -minlenltr 100 -maxlenltr 7000 -mindistltr 1000 -maxdistltr 20000 -similar 85 -motifmis 1 -mintsd 5 -overlaps best”, to generate a GFF3 file with entries for the LTR retrotransposon, as well as the two flanking target site duplications generated when the TE is inserted. We deleted the LTR retrotransposon as well as one of the two target site duplications in order to recapitulate the empty allele that would have existed before the TE was inserted. We merged adjacent TE deletion records together and formed a list of non-overlapping benchmark deletion records. We then aligned this TE-deleted genome to the B73 v4 genome to assess alignment quality in these highly repetitive regions. To evaluate the deletion recall ratios of the genome alignment approaches, we defined two deletion records as identical if by removing the sequence from the reference genome they could generate the identical synthetic genome sequences. This was done in order to avoid the impact of genomic variants that are represented in many different ways(18), e.g., when microhomology at the boundaries of a deletion means that both alignments are equally optimized.

We adjusted the parameters “-w 38000 -fa3 200000” of AnchorWave for genome alignment. The “genoali” command of AnchorWave was used to perform alignments for 18 Arabidopsis synthetic genomes against the TAIR10 reference genome and the alignment of the B73 v4 TE removed sequence against the B73 v4 reference genome. The parameter “-IV” of the AnchorWave “genoAli” command was appended to identify inversions between maize *de novo* assemblies against the maize B73 v4 reference genome. To align the sorghum genome against the maize genome, we used the “proali” command of AnchorWave with parameters “-w 38000 - fa3 200000 -R 1 -Q 2” to utilize the knowledge that the maize lineage has been through a whole-genome duplication since its divergence with sorghum(28).

The values of asm5, asm10, and asm20 of the -x setting of minimap2 v2.16-r922(16) were used separately. When aligning the TE-removed maize B73 genome sequence using a single thread, the setting asm20 gave an “insufficient memory” error on a computer with 2 terabyte memory available, and we did not obtain the corresponding result.

The “lastal” function from the LAST toolkit v932(11) was used to perform genome alignment with default parameters; the results were termed LAST many-to-many. Following previously described method(35), the LAST many-to-many results were transformed into psl format using the “maf-convert” comment of LAST, and the psl files were fed into the chain-net pipeline, “axtChain -linearGap=loose”, “chainMergeSort”, “chainPreNet”, “chainNet”, “netToAxt”, “axtSort” and “axtToMaf” in sequential order to generate the LAST many_to_one results. The LAST many_to_one results were further processed via the “last-split | maf-swap | last-split | maf-swap” command to generate LAST one_to_one results. “last-split” and “maf-swap” are components of the LAST genome alignment toolkits.

The parameters --sam-short of MUMmer4(20) were used to produce genome alignments in SAM format. We used the parameter “-fmt 1 “ of GSAlign(21) to perform genome alignments with default settings and output in MAF format.

To calculate the proportion of the reference genome that was aligned and matched, all the alignments in MAF format were reformatted into bam files using the “maf-convert sam” command of LAST and SAMtools v1.10(36). We used the “depth” command of SAMtools to calculate how many base pairs of the maize genome were aligned and calculated the proportion of reference genome that is aligned as the ratio of the number of aligned base pairs of the maize genome to maize genome size. We used the “samtools depth | awk ‘$3>0{print $0}’ | wc - l” command to calculate how many base pairs of the maize genome have their positions matched.

Due to the limit of available computational resources (maximum of 2 terabytes of memory), we split the maize B73 genome and maize Mo17 genome into individual chromosomes and performed alignments using LAST and minimap2 for each pair of homologous chromosomes independently. When aligning chromosome 1 using minimap2 asm20, we set “-w 19” to reduce memory usage to less than 2 terabytes. The outputs of minimaps2, MUMmer4, and GSAlign were filtered as one-to-one alignment for subsequent analysis using the “last-split | maf-swap | last-split | maf-swap” command as described above.

## Author contributions

B.S., M.C.S., and E.S.B. designed the analysis and wrote the manuscript. S.M.S. and M.M. implemented the 2-piece affine gap cost model in the WFA library. B.S. implemented the AnchorWave software. B.S. and L.J. optimized the longest path algorithm. All authors revised and reviewed the manuscript.

## Data availability

The source code of AnchorWave is hosted by GitHub at: https://github.com/baoxingsong/anchorwave.

All the commands, plotting scripts, and plot source data for this manuscript could be found at: https://github.com/baoxingsong/genomeAlignment.

## Acknowledgments

This project is supported by the USDA-ARS, National Science Foundation #1822330 and #1854828 to E.S.B. M.C.S was supported by NSF PRFB #1907343. Also, this project was supported by the European Union’s Horizon 2020 Framework Programme under the DeepHealth project [825111], and by the European Union Regional Development Fund within the framework of the ERDF Operational Program of Catalonia 2014-2020 with a grant of 50% of total cost eligible under the DRAC project [001-P-001723]. M.M. was partially supported by the Spanish Ministry of Economy, Industry, and Competitiveness under Ramón y Cajal fellowship number RYC-2016-21104. We thank the members of the Buckler lab (Cornell University, US) (Cornell University, US) for helpful discussions. We thank Travis Wrightsman (Cornell University, US) for suggesting the name AnchorWave, and Merritt Khaipho-Burch, Qi Sun, Cheng Zou, Minghui Wang, Zack Miller, Sara Miller (Cornell University, US), Jeffrey Ross-Ibarra (University of California, Davis, US), Shi Huang(University of California San Diego, US) for suggestions on the manuscript writing and proofreading.

## Supplementary Information

**Fig. S1.**
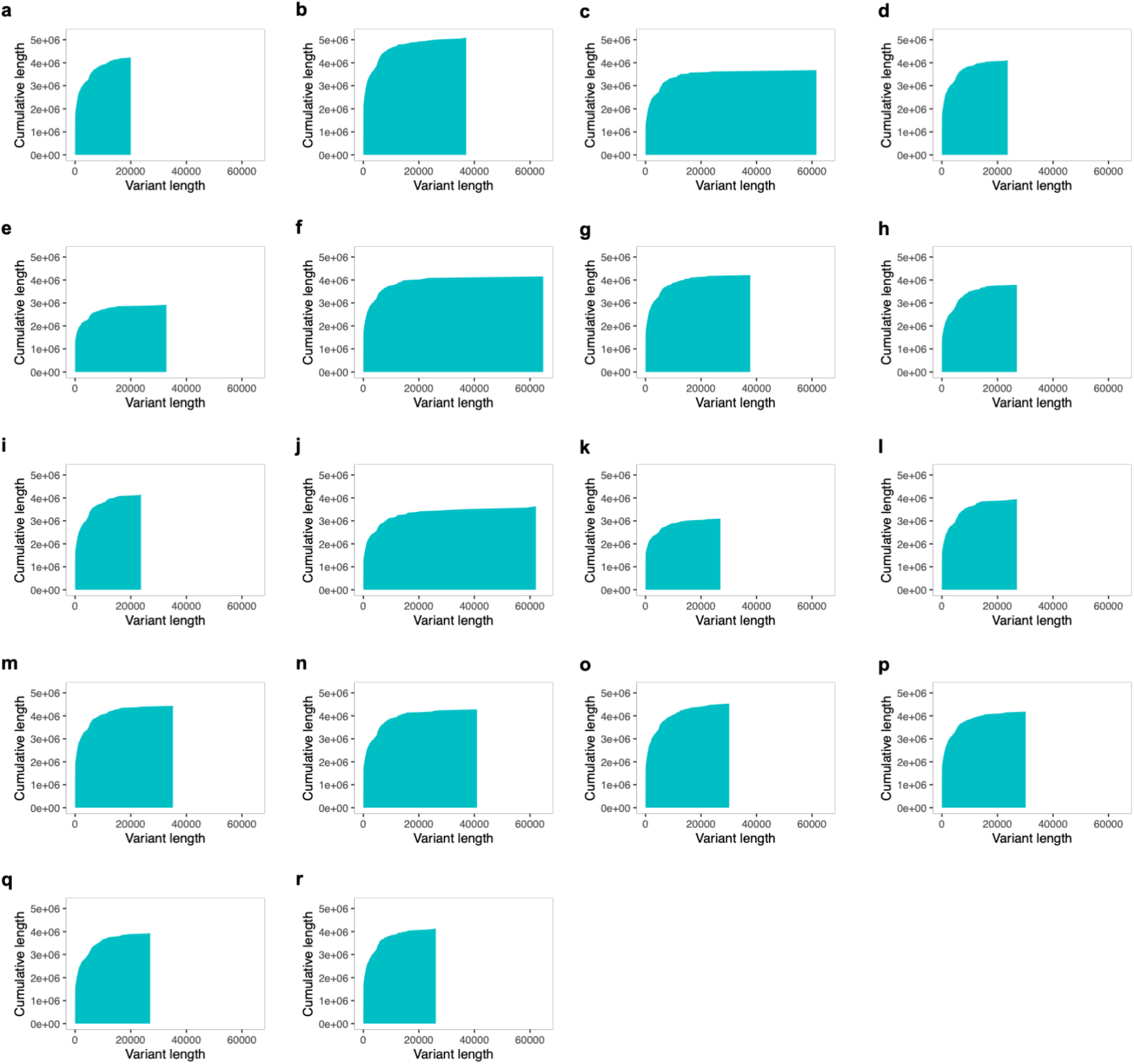
The length distribution of previously published variants for 18 Arabidopsis accessions. (a) Bur-0, (b) Can-0, (c) Ct-1, (d) Edi-0, (e) Hi-0, (f) Kn-0, (g) L*er*-0, (h) Mt-0, (i) No-0, (j) Oy-0, (k) Po-0, (l) Rsch-4, (m) Sf-2, (n) Tsu-0, (o) Wil-0, (p) Ws-0, (q) Wu-0, (r) Zu-0.

**Fig. S2.**
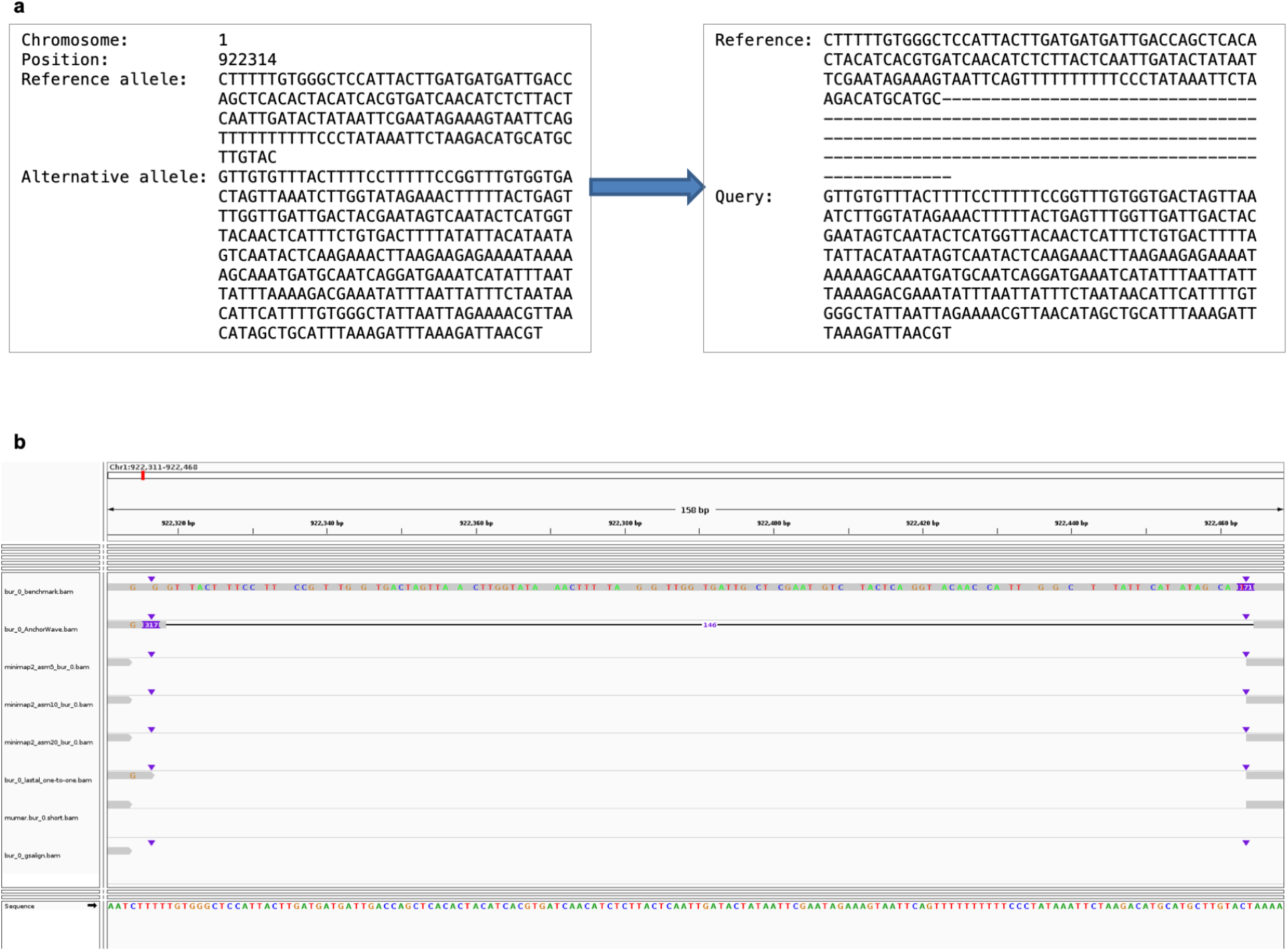
A variant record from Arabidopsis bur_0.v7c.sdi and the inferred benchmark alignment in this region. (a)[left] The reference allele in the TAIR10 genome spans 150 bp, and the variant record is neither an SNP nor an indel. [right] This region is not aligned in the benchmark, and the alignment across the region from AnchorWave is not expected to be consistent with the benchmark alignment, thus decreasing the precision of summarized AnchorWave alignments. (b) A screenshot showing the sequence alignments in this region generated by different tools. Tools except AnchorWave failed to perform sequence alignment in this region.

**Fig. S3.**
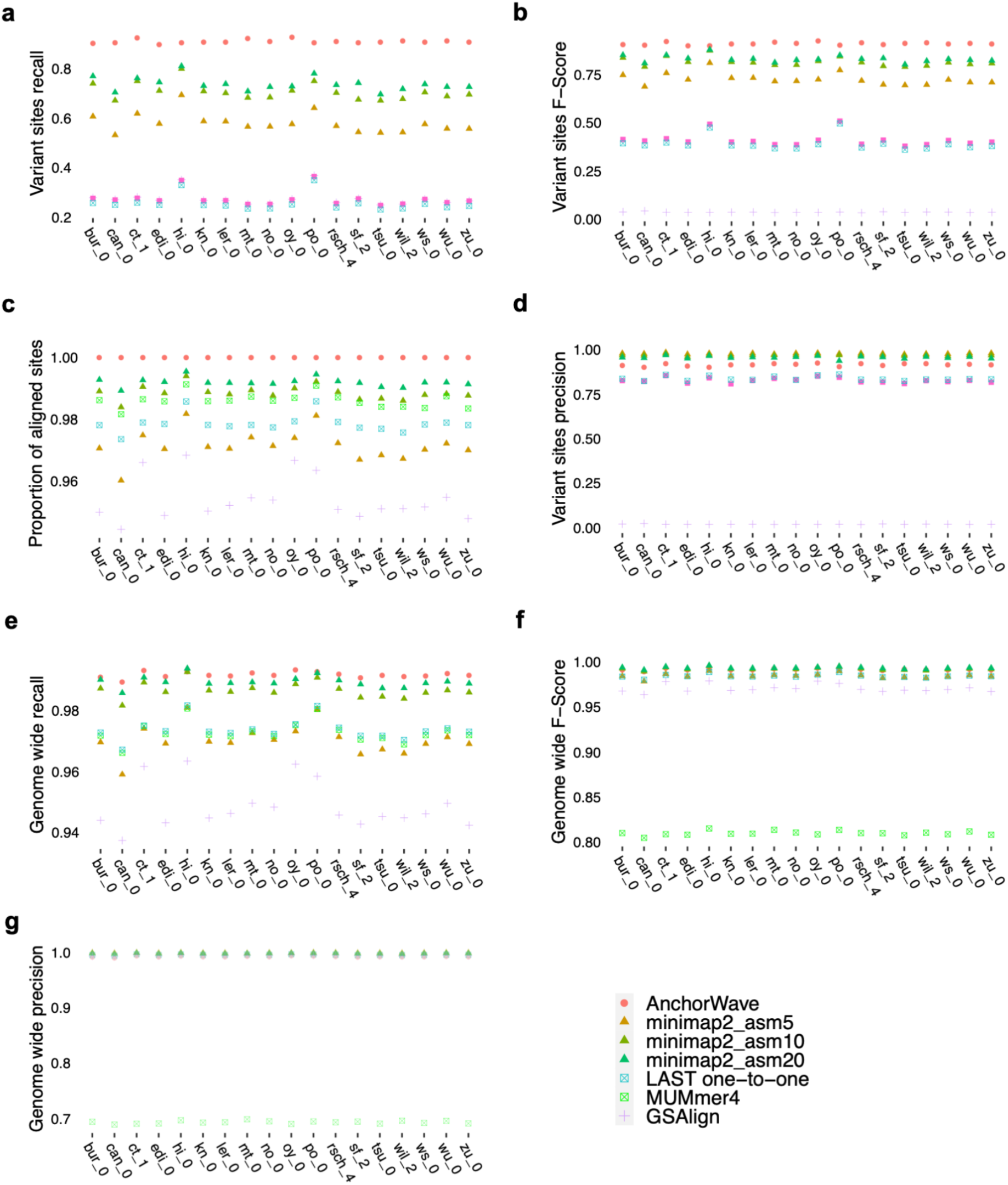
Genome alignment metrics using the synthetic benchmark alignments of 18 Arabidopsis accessions. The results of AnchorWave are ranked as highest or comparable to the highest values.

**Fig. S4.**
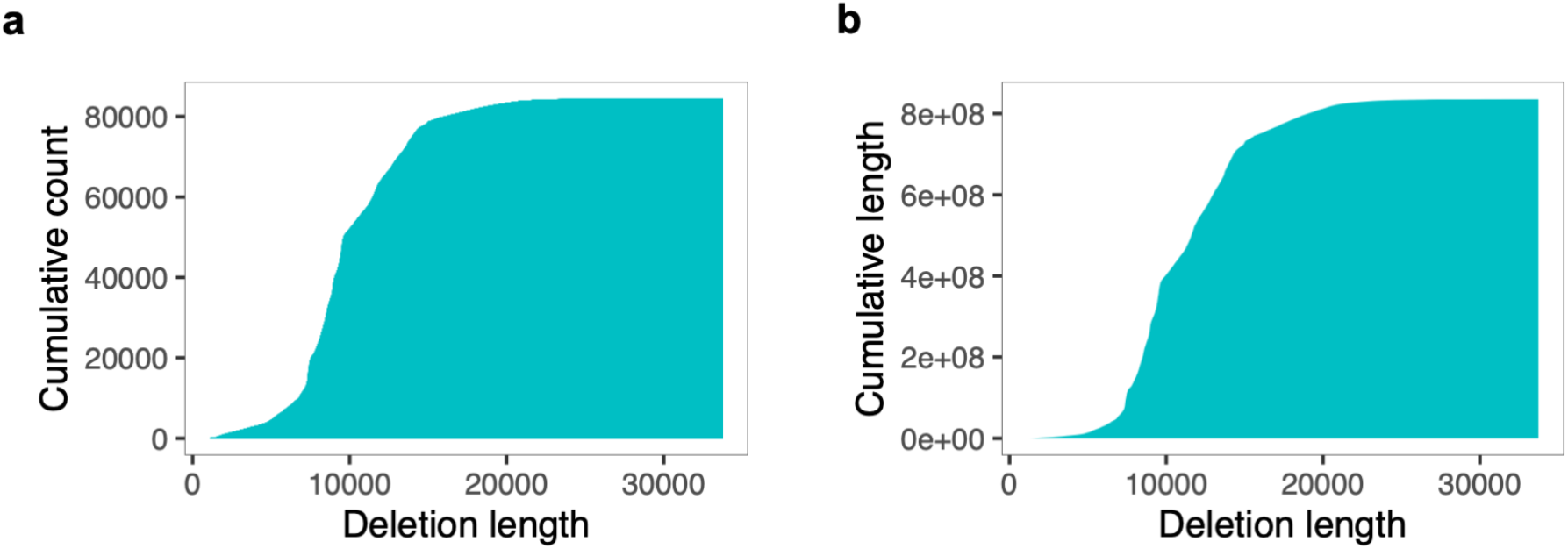
Length distribution of removed LTR retrotransposon deletions from the maize B73 V4 reference genome.

**Fig. S5.**
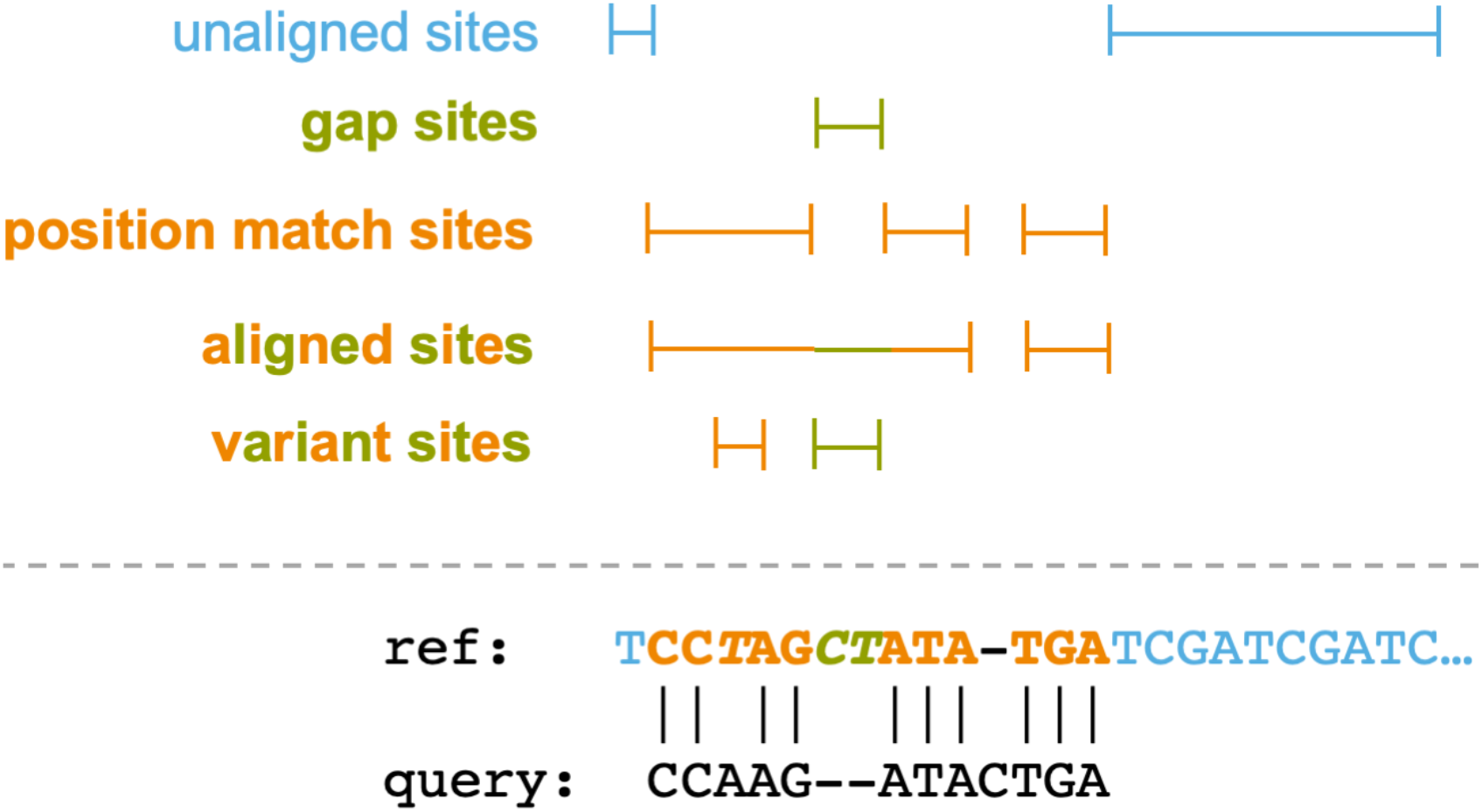
An example illustrating the concepts of (position) match sites, aligned sites, gap sites, and unaligned sites of the reference sequence used in this study. The ***T*** site of the reference sequence(ref) in orange, and italic font of the reference sequence is aligned as a nucleotide mismatch, but a position match.

**Fig. S6.**
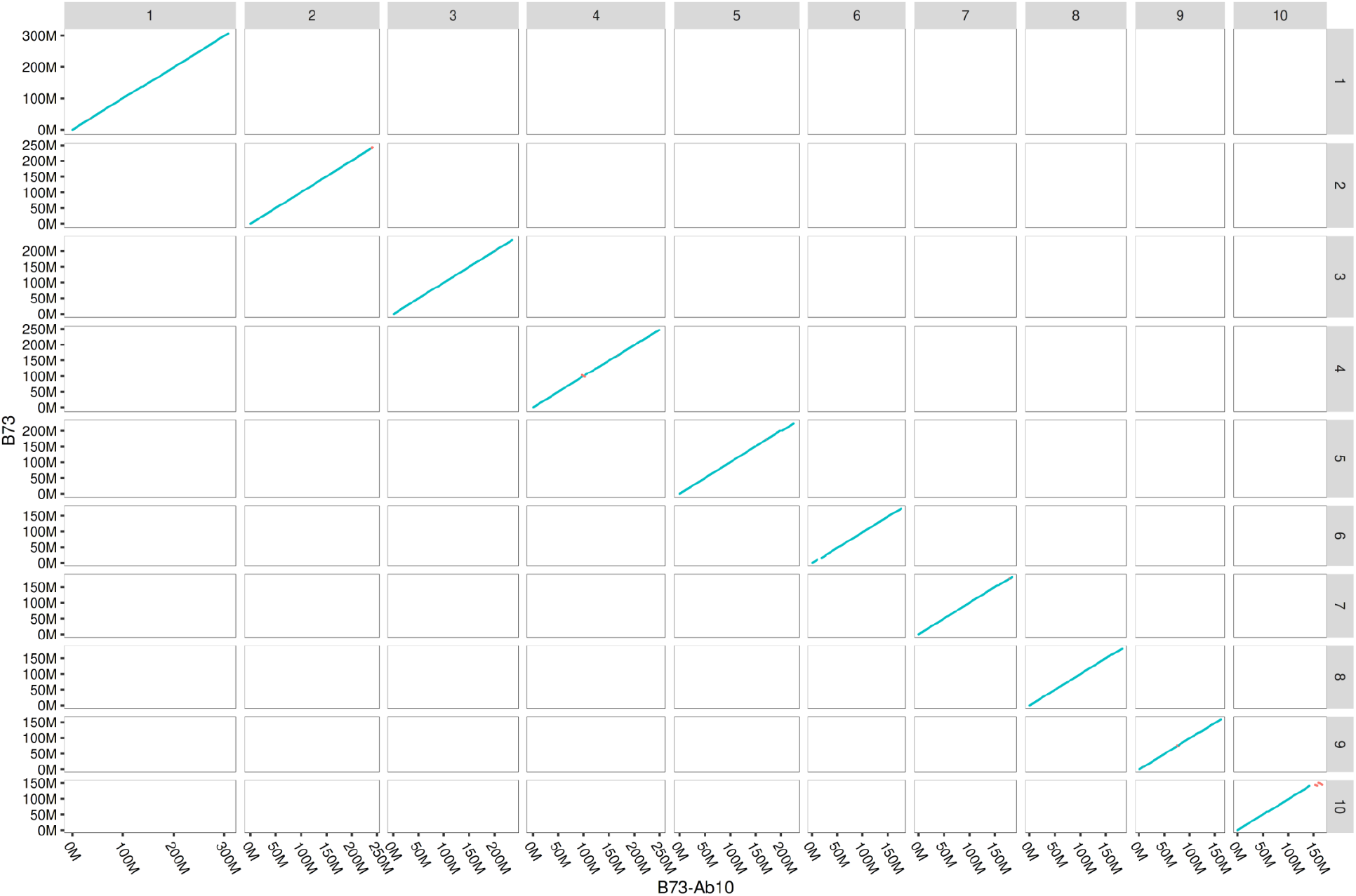
Collinear anchors between the maize B73 v4 and B73-AB10 genome assemblies. Each point was plotted based on the start position of an anchor in the reference genome (B73) and the query genome (B73-Ab10). Blue points represent anchors on the same strand in the reference genome and the query genome. Red points represent anchors on different strands in the reference genome and the query genome.

**Fig. S7.**
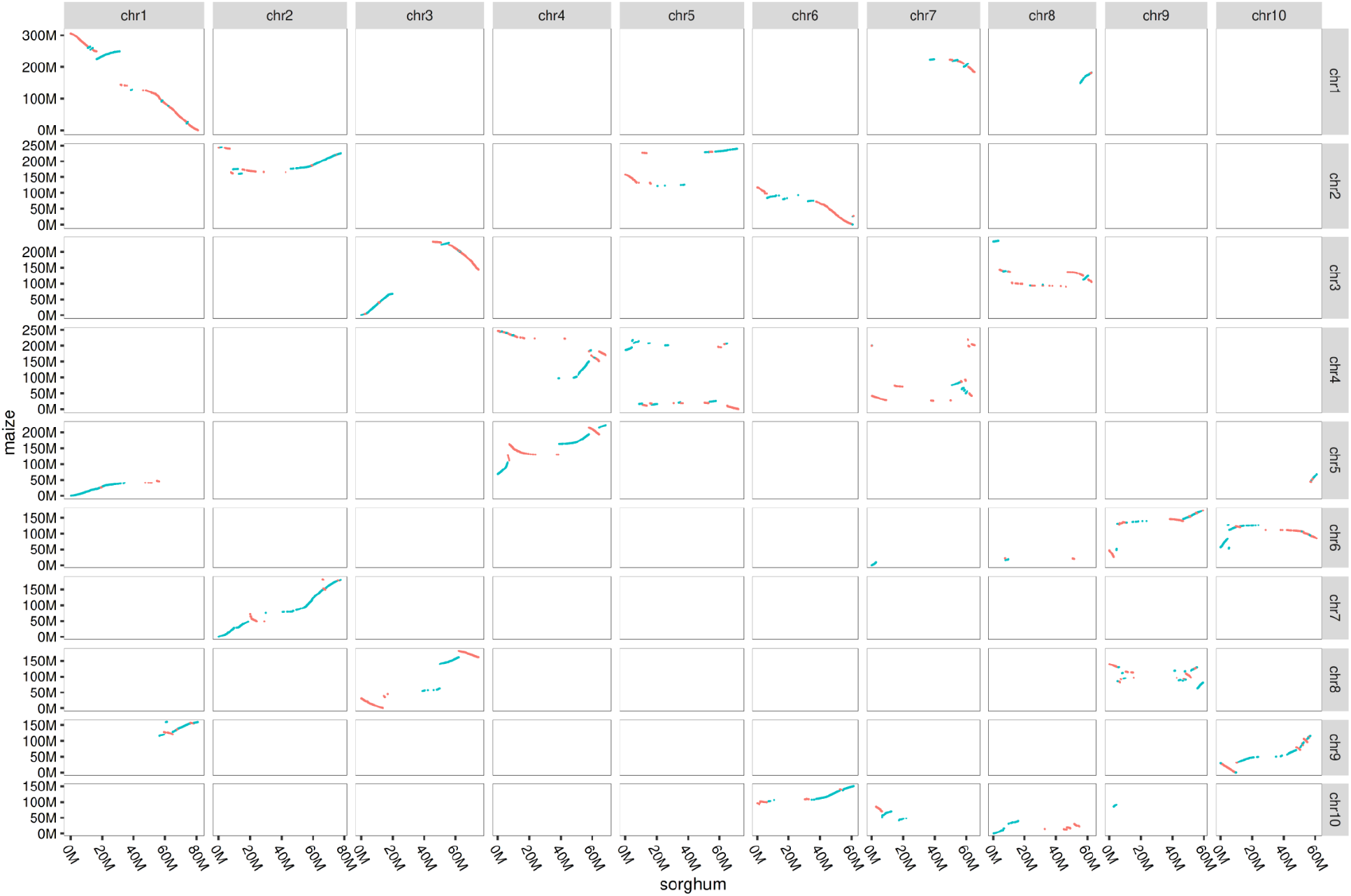
Collinear anchors between the maize B73 v4 and sorghum genome assemblies. Each point was plotted based on the start position of an anchor in the reference genome (maize) and the query genome (sorghum). Blue points represent anchors on the same strand in the reference genome and the query genome. Red points represent anchors on different strands in the reference genome and the query genome.

**Fig. S8.**
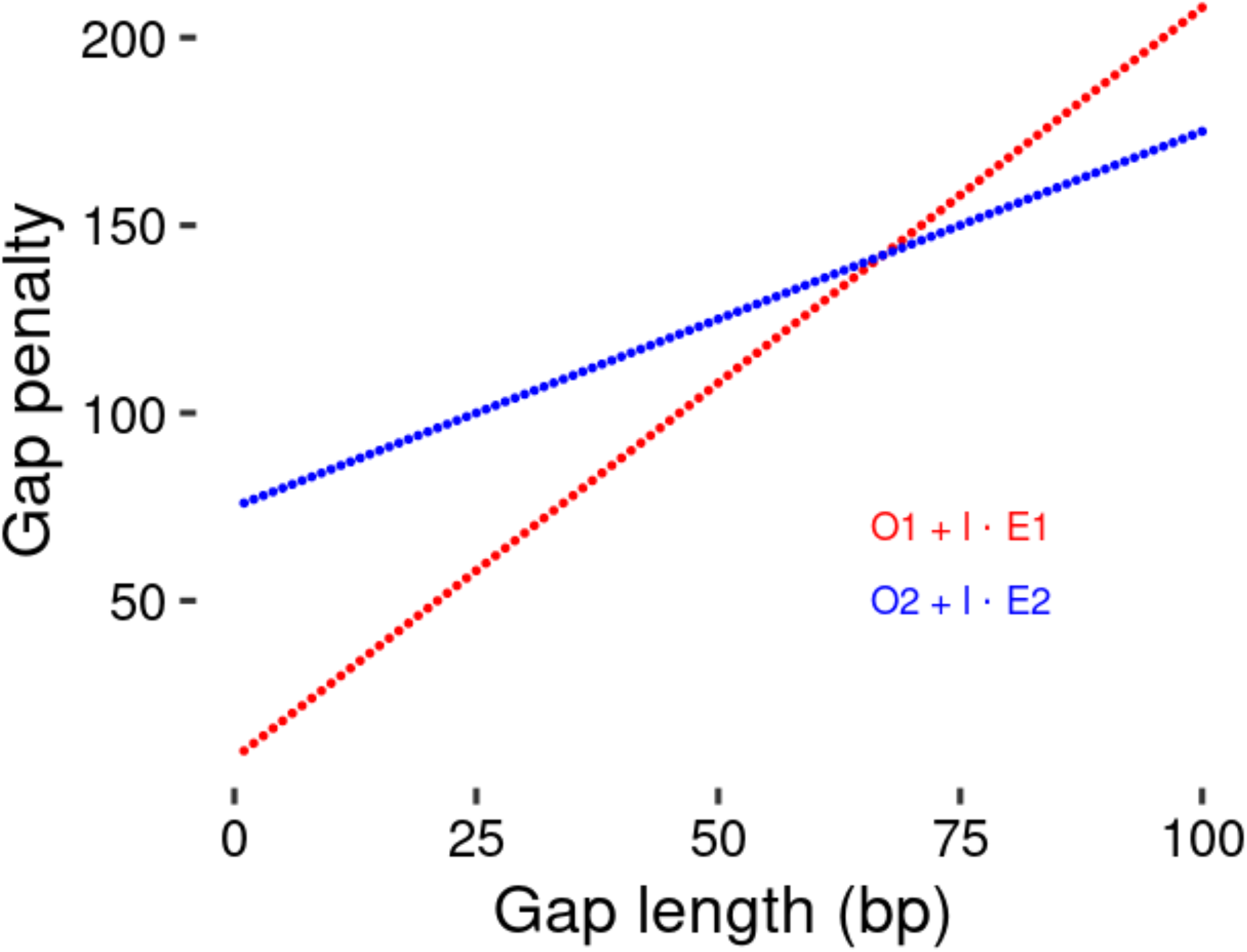
The 2-piece affine gap cost strategy as a dynamic programming approach for sequence alignment. The value of min{***O_1_***+|***l***| · ***E_1_, O_2_***+|***l***| · ***E_2_***} is used as the gap penalty. The aim is to model different types of mutational mechanisms that introduce indels of different length distributions.

**Dataset S1:**

Collinear anchors identified using AnchorWave between the maize B73 v4 assembly and the maize B73-Ab10 assembly.

**Dataset S2:**

Collinear anchors identified using AnchorWave between the maize B73 v4 assembly and the maize SK assembly.

**Table S1:** The CPU time and memory cost of testing software for aligning Arabidopsis synthetic genomes against the reference genome. The computational costs were tested on a computer with 512Gb RAM and Intel(R) Xeon(R) Gold 6230 CPU. Here, we used the single thread model for comparison purposes.

|  | memory | time(HH:MM:SS) | memory | time(HH:MM:SS) | memory | time(HH:MM:SS) | memory | time(HH:MM:SS) | memory | time(HH:MM:SS) | memory | time(HH:MM:SS) | memory | time(HH:MM:SS) |
| --- | --- | --- | --- | --- | --- | --- | --- | --- | --- | --- | --- | --- | --- | --- |
|  | AnchorWave |  | minimap2_asm5 |  | minimap2_asm10 |  | minimap2_asm20 |  | LAST |  | MUMmer4 |  | GSAalign |  |
| bur_0 | 9.9Gb | 00:06:13 | 1.0Gb | 00:02:15 | 1.0Gb | 00:01:52 | 1.7Gb | 00:02:30 | 11Gb | 00:32:48 | 1.5Gb | 00:08:13 | 0.8Gb | 00:01:41 |
| can_0 | 9.4Gb | 00:08:15 | 1.3Gb | 00:02:17 | 1.3Gb | 00:01:54 | 1.8Gb | 00:02:28 | 10Gb | 00:36:13 | 1.5Gb | 00:08:16 | 0.8Gb | 00:01:43 |
| ct_1 | 9.5Gb | 00:04:51 | 0.9Gb | 00:01:55 | 0.9Gb | 00:01:45 | 1.8Gb | 00:02:22 | 13Gb | 00:36:00 | 1.5Gb | 00:08:07 | 0.8Gb | 00:01:32 |
| edi_0 | 9.Gb | 00:06:05 | 1.2Gb | 00:02:12 | 1.2Gb | 00:01:53 | 1.9Gb | 00:02:29 | 10Gb | 00:34:08 | 1.5Gb | 00:08:07 | 0.8Gb | 00:01:51 |
| hi_0 | 7.8Gb | 00:03:59 | 0.9Gb | 00:01:58 | 0.9Gb | 00:01:41 | 1.7Gb | 00:02:28 | 13Gb | 00:35:07 | 1.5Gb | 00:07:38 | 0.7Gb | 00:01:46 |
| kn_0 | 9.6Gb | 00:05:57 | 1.1Gb | 00:02:07 | 1.1Gb | 00:01:51 | 1.7Gb | 00:02:30 | 13Gb | 00:33:21 | 1.5Gb | 00:07:47 | 0.8Gb | 00:01:39 |
| ler_0 | 9.6Gb | 00:06:17 | 1.2Gb | 00:02:10 | 1.2Gb | 00:01:52 | 1.7Gb | 00:02:31 | 12Gb | 00:35:25 | 1.5Gb | 00:07:46 | 0.8Gb | 00:01:52 |
| mt_0 | 8.8Gb | 00:05:27 | 1.0Gb | 00:02:07 | 1.0Gb | 00:01:46 | 1.7Gb | 00:02:27 | 15Gb | 00:36:07 | 1.5Gb | 00:07:44 | 0.8Gb | 00:01:48 |
| no_0 | 9.2Gb | 00:06:10 | 1.1Gb | 00:02:08 | 1.1Gb | 00:01:48 | 1.7Gb | 00:02:29 | 10Gb | 00:33:50 | 1.5Gb | 00:07:44 | 0.8Gb | 00:01:40 |
| oy_0 | 9.6Gb | 00:05:31 | 1.2Gb | 00:02:00 | 1.2Gb | 00:01:39 | 1.7Gb | 00:02:23 | 14Gb | 00:34:03 | 1.5Gb | 00:07:53 | 0.8Gb | 00:01:37 |
| po_0 | 27Gb | 00:03:21 | 1.1Gb | 00:02:05 | 1.1Gb | 00:01:48 | 1.7Gb | 00:02:27 | 6G | 00:33:49 | 1.5Gb | 00:07:55 | 0.7Gb | 00:01:51 |
| rsch_4 | 9.9Gb | 00:04:45 | 1.3Gb | 00:02:18 | 1.3Gb | 00:01:57 | 1.7Gb | 00:02:35 | 12Gb | 00:34:59 | 1.5Gb | 00:08:15 | 0.8Gb | 00:01:41 |
| sf_2 | 9.2Gb | 00:06:59 | 1.4Gb | 00:02:17 | 1.3Gb | 00:01:55 | 1.7Gb | 00:02:31 | 9Gb | 00:34:18 | 1.5Gb | 00:08:22 | 0.8Gb | 00:01:49 |
| tsu_0 | 8.8Gb | 00:06:38 | 1.4Gb | 00:02:12 | 1.2Gb | 00:01:51 | 1.7Gb | 00:02:28 | 13Gb | 00:33:52 | 1.5Gb | 00:09:09 | 0.8Gb | 00:01:49 |
| wil_2 | 9.4Gb | 00:07:22 | 1.1Gb | 00:02:09 | 1.1Gb | 00:01:53 | 1.7Gb | 00:02:27 | 11Gb | 00:35:32 | 1.5Gb | 00:08:28 | 0.8Gb | 00:01:38 |
| ws_0 | 9.4Gb | 00:06:17 | 1.3Gb | 00:02:12 | 1.2Gb | 00:01:52 | 1.7Gb | 00:02:27 | 11Gb | 00:34:17 | 1.5Gb | 00:08:40 | 0.8Gb | 00:01:46 |
| wu_0 | 9.1Gb | 00:06:00 | 1.0Gb | 00:02:13 | 1.0Gb | 00:01:51 | 1.7Gb | 00:02:27 | 10Gb | 00:33:24 | 1.5Gb | 00:08:31 | 0.8Gb | 00:01:39 |
| zu_0 | 9.3Gb | 00:06:21 | 1.2Gb | 00:02:11 | 1.2Gb | 00:01:50 | 1.7Gb | 00:02:28 | 12Gb | 00:35:42 | 1.5Gb | 00:08:45 | 0.8Gb | 00:01:52 |

**Table S2:**
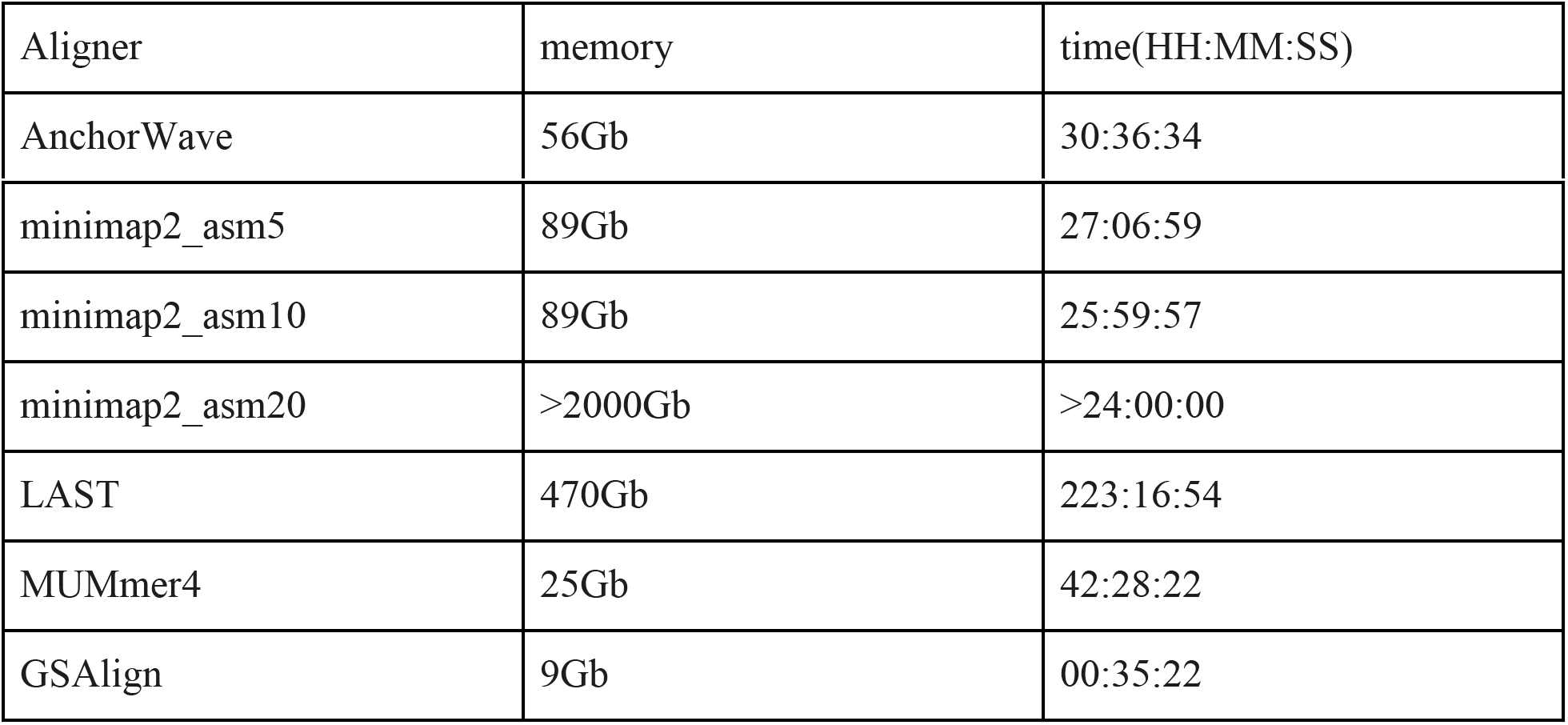
The CPU time and memory cost of testing software for aligning the TE removed maize B73 genome against the maize B73 reference genome. The computational costs of AnchorWave, minimap2 asm5, minimap2 asm10, LAST, Mummer4 and GSAlign, were tested on a computer with 512 gigabyte RAM and Intel(R) Xeon(R) Gold 6230 CPU. Minimap2 asm20 was tested on a computer with 2 terabytes of memory and AMD EPYC 7702 Processor CPU, but gave an insufficient memory error. We could not access a machine with more memory installed.

**Table S3:** The CPU time and memory cost of testing software for aligning the sorghum genome against the maize B73 v4 genome assembly. The computational costs were tested on a computer with 512Gb RAM and Intel(R) Xeon(R) Gold 6230 CPU.

| Aligner | memory | time(HH:MM:SS) |
| --- | --- | --- |
| AnchorWave | 81Gb | 132:07:30 |
| minimap2_asm5 | 7Gb | 00:08:57 |
| minimap2_asm10 | 7Gb | 00:08:00 |
| minimap2_asm20 | 10Gb | 00:12:46 |
| LAST | 21Gb | 07:48:59 |
| MUMmer4 | 25Gb | 00:39:38 |
| GSAlign | 12Gb | 00:34:51 |

**Supplementary Note 1:**

Assessing the quality of an alignment is difficult, as the evolutionary history of sequence orthology is unknown, and simulations do not recover the intricacies of sequence evolution(37). We tested genome alignments using a published variant call dataset of 18 Arabidopsis accessions, originally conducted using a hybrid approach of read mapping and *de novo* assembly(19).

The TAIR10 reference genome sequence and genome annotation(38) were downloaded by commands:

~~~
wget https://www.arabidopsis.org/download_files/Genes/TAIR10_genome_release/TAIR10_gff3/TAIR10_GFF3_genes.gff
wget ftp://ftp.arabidopsis.org/home/tair/Sequences/whole_chromosomes/TAIR10_chr1.fas
wget ftp://ftp.arabidopsis.org/home/tair/Sequences/whole_chromosomes/TAIR10_chr2.fas
wget ftp://ftp.arabidopsis.org/home/tair/Sequences/whole_chromosomes/TAIR10_chr3.fas
wget ftp://ftp.arabidopsis.org/home/tair/Sequences/whole_chromosomes/TAIR10_chr4.fas
wget ftp://ftp.arabidopsis.org/home/tair/Sequences/whole_chromosomes/TAIR10_chr5.fas
cat TAIR10_chr1.fas TAIR10_chr2.fas TAIR10_chr3.fas TAIR10_chr4.fas TAIR10_chr5.fas > tair10.fa
~~~

The variant calling results were downloaded from “http://mtweb.cs.ucl.ac.uk/mus/www/19genomes/variants.SDI”.

For each accession, the reference alleles from the TAIR10(38) genome were replaced with alternative alleles to generate a synthetic genome using the “pseudogeno” command of GEAN(18) with default parameters. GEAN reported error messages for ler_0.v7c.sdi, no_0.v7c.sdi and oy_0.v7c.sdi, due to multiple variants located at the same position. We removed the following two records from ler_0.v7c.sdi manually:

~~~
Chr4 6020491 12 AAGACATCAATATCATCAGGAAAAATACTCATTCTATTATTAG
TAATGACTTAGAGAATAAACTACGAATACAAAAAAAAAAACTCATTATATATCAT 5
Chr4 6020491 33 - TAATGACTTAGAGAATAAACTACGAATACAAAA 4
~~~

We removed the following two records from no_0.v7c.sdi manually:

~~~
Chr5 8499298 -1
CAACAAGTTAAAGATTTAAGGTTTTAAAATCCATTTTATAAATAAATTTTTATTGCAAAACTATATTGGAAATCAGAGAAATTTGAAA
AATTCACTTTTAAAGTTTTTTAATAACTGATAACGTACTACTTCAAAAACTGTAAACCAAATGCTAAGAATGCATATATGTTTCTGG
CAACAAAGAGATAACAGATA
GAGTGATAAAAAGATAAATAGCCTATCTTTTTGTTGGATATCCAACAAGTTGAAGGTTAAGGTTTTAAAAGATCCCAGTTTATAGC
AAAACTTAAGTTAGAAATCAGAGATATATATTTACAAAAATCACTATTCAATTTTTTTTATAATTGATTACTACTTCAAAAACTAGAA
ACCAAAAGCCAAGAATGTAT 5
Chr5 8499298 42 - GAGTGATAAAAAGATAAATAGCCTATCTTTTTGTTGGATATC 4
~~~

We removed the following two records from oy_0.v7c.sdi manually:

~~~
Chr2 7677637 7 ACAAGATAAATATTTAAAAATATATTTCTTAGA GCTAAAGTAATGAAAAAGTGTTTCTAAAACACTAGTTAAT 5
Chr2 7677637 8 - GCTAAAGT 4
~~~

The synthetic genomes were generated using the “pseudogeno” function of GEAN(18):

~~~
gean pseudogeno -r tair10.fa -v bur_0.v7c.sdi -o bur_0.fa
gean pseudogeno -r tair10.fa -v can_0.v7c.sdi -o can_0.fa
gean pseudogeno -r tair10.fa -v ct_1.v7c.sdi -o ct_1.fa
gean pseudogeno -r tair10.fa -v edi_0.v7c.sdi -o edi_0.fa
gean pseudogeno -r tair10.fa -v hi_0.v7c.sdi -o hi_0.fa
gean pseudogeno -r tair10.fa -v kn_0.v7c.sdi -o kn_0.fa
gean pseudogeno -r tair10.fa -v ler_0.v7c.sdi -o ler_0.fa
gean pseudogeno -r tair10.fa -v mt_0.v7c.sdi -o mt_0.fa
gean pseudogeno -r tair10.fa -v no_0.v7c.sdi -o no_0.fa
gean pseudogeno -r tair10.fa -v oy_0.v7c.sdi -o oy_0.fa
gean pseudogeno -r tair10.fa -v po_0.v7c.sdi -o po_0.fa
gean pseudogeno -r tair10.fa -v rsch_4.v7c.sdi -o rsch_4.fa
gean pseudogeno -r tair10.fa -v sf_2.v7c.sdi -o sf_2.fa
gean pseudogeno -r tair10.fa -v tsu_0.v7c.sdi -o tsu_0.fa
gean pseudogeno -r tair10.fa -v wil_2.v7c.sdi -o wil_2.fa
gean pseudogeno -r tair10.fa -v ws_0.v7c.sdi -o ws_0.fa
gean pseudogeno -r tair10.fa -v wu_0.v7c.sdi -o wu_0.fa
gean pseudogeno -r tair10.fa -v zu_0.v7c.sdi -o zu_0.fa
~~~

The benchmark genome alignments were generated using commands:

~~~
anchorwave sdiToMaf -o col_bur.maf -r tair10.fa -s bur_0.sdi -q bur_0.fa
anchorwave sdiToMaf -o col_can.maf -r tair10.fa -s can_0.sdi -q can_0.fa
anchorwave sdiToMaf -o col_ct.maf -r tair10.fa -s ct_1.sdi -q ct_1.fa
anchorwave sdiToMaf -o col_edi.maf -r tair10.fa -s edi_0.sdi -q edi_0.fa
anchorwave sdiToMaf -o col_hi.maf -r tair10.fa -s hi_0.sdi -q hi_0.fa
anchorwave sdiToMaf -o col_kn.maf -r tair10.fa -s kn_0.sdi -q kn_0.fa
anchorwave sdiToMaf -o col_ler.maf -r tair10.fa -s ler_0.sdi -q ler_0.fa
anchorwave sdiToMaf -o col_mt.maf -r tair10.fa -s mt_0.sdi -q mt_0.fa
anchorwave sdiToMaf -o col_no.maf -r tair10.fa -s no_0.sdi -q no_0.fa
anchorwave sdiToMaf -o col_oy.maf -r tair10.fa -s oy_0.sdi -q oy_0.fa
anchorwave sdiToMaf -o col_po.maf -r tair10.fa -s po_0.sdi -q po_0.fa
anchorwave sdiToMaf -o col_rsch.maf -r tair10.fa -s rsch_4.sdi -q rsch_4.fa
anchorwave sdiToMaf -o col_sf.maf -r tair10.fa -s sf_2.sdi -q sf_2.fa
anchorwave sdiToMaf -o col_tsu.maf -r tair10.fa -s tsu_0.sdi -q tsu_0.fa
anchorwave sdiToMaf -o col_wil.maf -r tair10.fa -s wil_2.sdi -q wil_2.fa
anchorwave sdiToMaf -o col_ws.maf -r tair10.fa -s ws_0.sdi -q ws_0.fa
anchorwave sdiToMaf -o col_wu.maf -r tair10.fa -s wu_0.sdi -q wu_0.fa
anchorwave sdiToMaf -o col_zu.maf -r tair10.fa -s zu_0.sdi -q zu_0.fa
~~~

We extracted full-length CDS of the TAIR10 reference genome using the command:

~~~
anchorwave gff2seq -r tair10.fa -i TAIR10_GFF3_genes.gff -o cds.fa
~~~

We mapped full-length CDS to the reference genome using minimap2(16) with the command:

~~~
minimap2 -x splice -t 4 -k 12 -a -p 0.4 -N 20 tair10.fa cds.fa > ref.sam
~~~

We mapped full-length CDS to each synthetic genome using the command:

~~~
minimap2 -x splice -t 4 -k 12 -a -p 0.4 -N 20 synthesis_genomes.fa cds.fa > synthesis_genome.sam
~~~

AnchorWave genome alignments were conducted via the command:

~~~
anchorwave genoAli -i TAIR10_GFF3_genes.gff -as cds.fa -r tair10.fa -a synthesis_genome.sam -ar ref.sam -s synthesis_genome.fa -v synthesis_genome.vcf -n synthesis_genome.anchors -o synthesis_genome.maf -f synthesis_genome.f.maf -w 38000 -fa3 200000
~~~

We used minimap2 to align each synthetic genome against the reference genome with three parameter sets using the commands:

~~~
minimap2 -x asm5 -t 1 -a tair10.fa synthesis_genome.fa > minimap2_asm5_synthesis_genome.sam
minimap2 -x asm10 -t 1 -a tair10.fa synthesis_genome.fa > minimap2_asm10_synthesis_genome.sam
minimap2 -x asm20 -t 1 -a tair10.fa synthesis_genome.fa > minimap2_asm20_synthesis_genome.sam
~~~

These output files in SAM format were reformatted into MAF using the “sam2maf” function implemented in AnchorWave:

~~~
anchorwave sam2maf -r tair10.fa -q synthesis_genome.fa -s minimap2_asm5_synthesis_genome.sam -o minimap2_asm5_synthesis_genome.maf
~~~

The LAST(11) genome alignments were conducted using commands:

~~~
lastdb col tair10.fa
faToTwoBit tair10.fa col.2bit
faSize -detailed tair10.fa > col.size
lastal col synthesis_genome.fa > synthesis_genome_lastal.maf
faSize -detailed synthesis_genome.fa > synthesis_genome.size
faToTwoBit synthesis_genome.fa synthesis_genome.2bit
maf-convert psl synthesis_genome_lastal.maf > synthesis_genome_lastal.psl
axtChain -linearGap=loose -psl synthesis_genome_lastal.psl -faQ -faT tair10.fa synthesis_genome.fa synthesis_genome_lastal.chain
chainMergeSort synthesis_genome_lastal.chain > synthesis_genome_lastal.all.chain
chainPreNet synthesis_genome_lastal.all.chain col.size synthesis_genome.size synthesis_genome_lastal.preNet
chainNet synthesis_genome_lastal.preNet col.size synthesis_genome.size synthesis_genome_lastal.refTarget.net synthesis_genome_lastal.chainNet
netToAxt synthesis_genome_lastal.refTarget.net synthesis_genome_lastal.preNet col.2bit synthesis_genome.2bit stdout | axtSort stdin synthesis_genome_lastal.axt
axtToMaf synthesis_genome_lastal.axt col.size synthesis_genome.size synthesis_genome_lastal_final.maf -qPrefix=query. - tPrefix=col.
perl lastFinalToSplit.pl synthesis_genome_lastal_final.maf > synthesis_genome_lastal_final_forsplit.maf
cat synthesis_genome_lastal.maf | last-split | maf-swap | last-split | maf-swap > synthesis_genome_lastal_split.maf
cat synthesis_genome_lastal_final_forsplit.maf | last-split | maf-swap | last-split | maf-swap > synthesis_genome_lastal_final_split.maf
~~~

The header of MAF files is not compatible between the LAST pipeline and the chain-net pipeline. We implemented “lastFinalToSplit.pl” to reformat the header of MAF files to make them compatible. Those two scripts have been released under “src/tests/scripts/’ of the AnchorWave source code repository. The output file “synthesis_genome_lastal.maf” is termed LAST many-to-many alignment, the output file “synthesis_genome_lastal_final.maf” is termed LAST many-to-one alignment, and the output of “synthesis_genome_lastal_final_split.maf” is termed LAST one-to-one alignment.

We performed alignments using MUMmer4(20) via the command:

~~~
nucmer -t 1 --sam-short=mumer.synthesis_genome.short.sam tair10.fa synthesis_genome.fa
~~~

We aligned synthesis genomes against the TAIR10 reference genome using GSAlign(21) via the command:

~~~
GSAlign -t 1 -r tair10.fa -q synthesis_genome.fa -t 78 -o synthesis_genome_gsalign -fmt 1
~~~

The scripts to summarize alignment recall, precision and F-score, “CompareMafAndCheckMafCoverage.py” and “comparemafandcheckmafcoverageAll.py”, are available in the “./src/tests/scripts/” folder of the source code repository.

AnchorWave outperforms other approaches with the highest recall for variant sites (0.910 on average, the second-best is 0.737 achieved via minimap2 asm20) and for genome-wide sites(0.992 on average, the second-best is 0.989, achieved via minimap2 asm20). Moreover, AnchorWave is the only implementation that performs end-to-end alignment, and the second-highest proportion of aligned sites (0.992) was achieved by minimap2 asm20. The precision of AnchorWave is lower than all parameter sets of minimap2 we tested (6% lower for variant sites and 0.7% lower genome-wide compared to minimap2 asm5, which is ranked the highest). For the genome-wide sites F-score, AnchorWave (0.9917 on average) is ranked as the third-highest and lower than that of minimap2 asm20 (0.9931 on average) and minimap2 asm10 (0.9922 on average). For variant sites, the average F-score of AnchorWave was 0.912, which is the highest one and the second highest one was 0.832 generated by minimap2 asm20 (Fig. S3).

**Supplementary Note 2:**

We tested the performance of AnchorWave for the detection of long indels in repeat-rich genomes by removing retrotransposons from the maize B73 v4 genome assembly and aligning the synthetic genome against the reference genome.

The B73 v4 reference genome sequence and genome annotation(22) were downloaded using commands:

~~~
wget ftp://ftp.ensemblgenomes.org/pub/plants/release-34/gff3/zea_mays/Zea_mays.AGPv4.34.gff3.gz
gunzip Zea_mays.AGPv4.34.gff3.gz
wget ftp://ftp.ensemblgenomes.org/pub/plants/release-34/fasta/zea_mays/dna/Zea_mays.AGPv4.dna.toplevel.fa.gz
gunzip Zea_mays.AGPv4.dna.toplevel.fa.gz
~~~

To generate a GFF3 format LTR retrotransposon annotation, we use ltrharvest(34). First, we generate an index of the genome:

~~~
genometools-1.5.7/bin/gt suffixerator -db Zea_mays.AGPv4.dna.toplevel.fa.gz -indexname B73V4 -tis -suf -lcp -des -ssp -sds -dna - memlimit 48GB
~~~

Then, we search for LTR retrotransposons using ltrharvest:

~~~
genometools-1.5.7/bin/gt ltrharvest -index B73V4 -gff3 B73V4.ltrharvest.gff3 -motif tgca -minlenltr 100 -maxlenltr 7000 -mindistltr 1000 -maxdistltr 20000 -similar 85 -motifmis 1 -mintsd 5 -xdrop 5 -overlaps best -longoutput -out B73V4.fa > B73v4.ltrharvest.out
~~~

Finally, since genometools names sequences with an internal identifier, we convert the sequence names in GFF3 file back to chromosome names:

~~~
wget https://raw.githubusercontent.com/mcstitzer/maize_v4_TE_annotation/master/ltr/mask_subtract/convert_ltrharvest_seq_gff_to_contignames.py
python2 convert_ltrharvest_seq_gff_to_contignames.py B73V4.ltrharvest.gff3 > B73V4.ltrharvest.contignames.gff3
~~~

We use this GFF3 to generate a FASTA file of the genome with these TEs removed (B73V4.pseudomolecule.subtract1.fa), by removing both the LTR retrotransposon and one of the target site duplications generated when it inserted in the genome. This is implemented in the “gffToMaf, c1” function located within “./src/tests/impl/TEGffToAlignment.cpp” of the AnchorWave source code repository. This function replaces each nucleotide within TE regions of maize B73 v4 genome with After all the nucleotides within TE regions are replaced, this function removes the characters and outputs the remaining nucleotides in FASTA format. We created a function “gff2Vcf” under “./src/tests/impl/TEGffToAlignment.cpp” to create the variant records file “B73V4.pseudomolecule.ltrharvest.contignames.gff3.contigpositions.vcf”. This function replaces each nucleotide within TE regions of maize B73 v4 genome with and generates a pairwise sequence alignment and then performs variant calling. The length distribution of deletion records in “B73V4.pseudomolecule.ltrharvest.contignames.gff3.contigpositions.vcf” is shown in (Fig. S4).

We aligned “B73V4.pseudomolecule.subtract1.fa” against the B73 v4 reference genome using AnchorWave, minimap2, LAST, MUMmer4, and GSAlign using the same settings as described in Supplementary Note 1. The genome alignment files in SAM(36) format were transformed into MAF format using the “sam2maf” function implemented in AnchorWave, and the function “maf2vcf” of AnchorWave was used to generate variants in VCF format(39). We implemented the “evaluateTEAlignment” function to compare generated VCF files with the benchmark file “B73V4.pseudomolecule.ltrharvest.contignames.gff3.contigpositions.vcf”.

As discussed in the main text, AnchorWave shows the highest recall of these TE deletions. Besides AnchorWave, minimap2 is the only method tested that could recall long indels. This may be due to minimap2’s usage of global alignment between adjacent anchors in a chain. There is no obvious length distribution difference between correctly recalled deletions and incorrectly recalled deletions for both minimap2 and AnchorWave (Figure S1).

**Figure S1.**
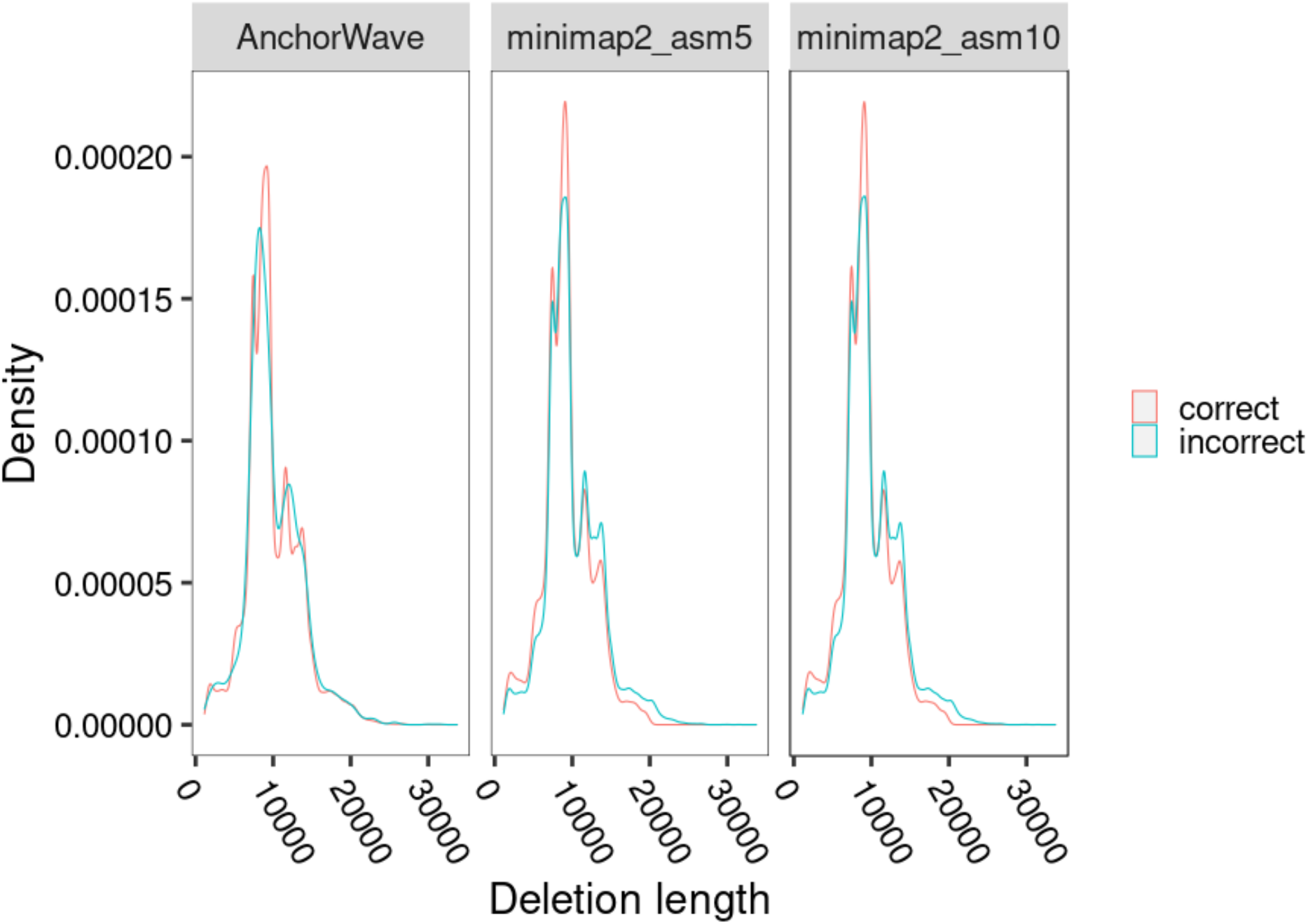
The length distribution of TE deletion records being correctly and incorrectly recalled using AnchorWave and minimap2.

**Supplementary Note 3:**

We downloaded the maize Mo17 genome in FASTA format and renamed chromosome to be consistent with the B73 genome FASTA file using the commands:

~~~
wget https://download.maizegdb.org/Zm-Mo17-REFERENCE-CAU-1.0/Zm-Mo17-REFERENCE-CAU-1.0.fa.gz
gunzip Zm-Mo17-REFERENCE-CAU-1.0.fa.gz
sed -i ‘s/>chr/>/g’ Zm-Mo17-REFERENCE-CAU-1.0.fa
~~~

The B73 V4 reference genome sequence and genome annotation were downloaded as described in Supplementary Note 2. The reference full-length CDSs were extracted and mapped to the reference genome and the query genome as described in Supplementary Note 1.

We performed genome alignment using the following command:

~~~
anchorwave genoAli -i Zea_mays.AGPv4.34.gff3 -as cds.fa -r Zea_mays.AGPv4.dna.toplevel.fa -a cds.sam -ar ref.sam -s Zm-Mo17-REFERENCE-CAU-1.0.fa -n anchors -o anchorwave.maf -f anchorwave.f.maf -w 38000 -fa3 200000 -t 1 -IV
~~~

The minimap2 and MUMmer4 genome alignments were reformatted into MAF format using the “anchorwave sam2maf” function. The minimap2, MUMmer4, and GSalign results were purified into one-to-one alignment using the “last-split | maf-swap | last-split | maf-swap” as described in Supplementary Note 1.

The recall of TE PAVs was evaluated using a script available under the AnchorWave source code repository at “src/tests/scripts/evaluateTePavALignment.pl”. This script evaluates whether deletions from a sam file overlap precisely the positions of features in an input bed file. We used site-defined TE PAVs that differ between B73 and Mo17 from Anderson et al. (2019) (https://github.com/SNAnderson/maizeTE_variation/raw/master/non-redundant_TEs_4genomes_1Feb19.txt.gz), adding the length of a target site duplication (TSD) appropriate for the TE superfamily to generate a bed file.

**Supplementary Note 4:**

The maize B73-Ab10 genome(40) in FASTA format was downloaded and we renamed the entry names to be consistent with the B73 genome file using the commands:

~~~
wget https://ftp.ncbi.nlm.nih.gov/genomes/all/GCA/902/714/155/GCA_902714155.1_Zm-B73_AB10-REFERENCE-NAM-1.0b/GCA_902714155.1_Zm-B73_AB10-REFERENCE-NAM-1.0b_genomic.fna.gz
gunzip GCA_902714155.1_Zm-B73_AB10-REFERENCE-NAM-1.0b_genomic.fna.gz
sed -i ‘s/, whole genome shotgun sequence//’ GCA_902714155.1_Zm-B73_AB10-REFERENCE-NAM-1.0b_genomic.fna
sed -i ‘s/>.*Zea mays genome assembly, chromosome: />/’ GCA_902714155.1_Zm-B73_AB10-REFERENCE-NAM-1.0b_genomic.fna
~~~

The B73 V4 reference genome sequence and genome annotation were downloaded as described in Supplementary Note 2. The reference full-length CDSs were extracted and mapped to the reference genome and query genome as introduced in Supplementary Note 1. We detected collinear anchors using the following command:

~~~
anchorwave genoAli -i Zea_mays.AGPv4.34.gff3 -r Zea_mays.AGPv4.dna.toplevel.fa -as cds.fa -a cds.sam -ar ref.sam -s GCA_902714155.1_Zm-B73_AB10-REFERENCE-NAM-1.0b_genomic.fna -n anchorsiv -IV -ns
~~~

We further test the performance of AnchorWave for inversion detection by comparing with inversions reported between the maize B73 and SK lines(41). We downloaded the maize SK genome in FASTA format and renamed entry names to be consistent with the B73 genome file using commands:

~~~
wget
ftp://download.big.ac.cn/gwh/Plants/Zea_mays_the_genome_of_SK_GWHAACS00000000/GWHAACS00000000.genome.fasta.gz
gunzip GWHAACS00000000.genome.fasta.gz
sed -i ‘s/>.*Chromosome\s/>/g’ GWHAACS00000000.genome.fasta
sed -i ‘s/\s.*//g’ GWHAACS00000000.genome.fasta
~~~

The reference full-length CDSs were extracted and mapped to the reference genome and the query genome as described in Supplementary Note 1.

The CDS mapping on the query genome is visualized as Figure S2.

**Figure S2.**
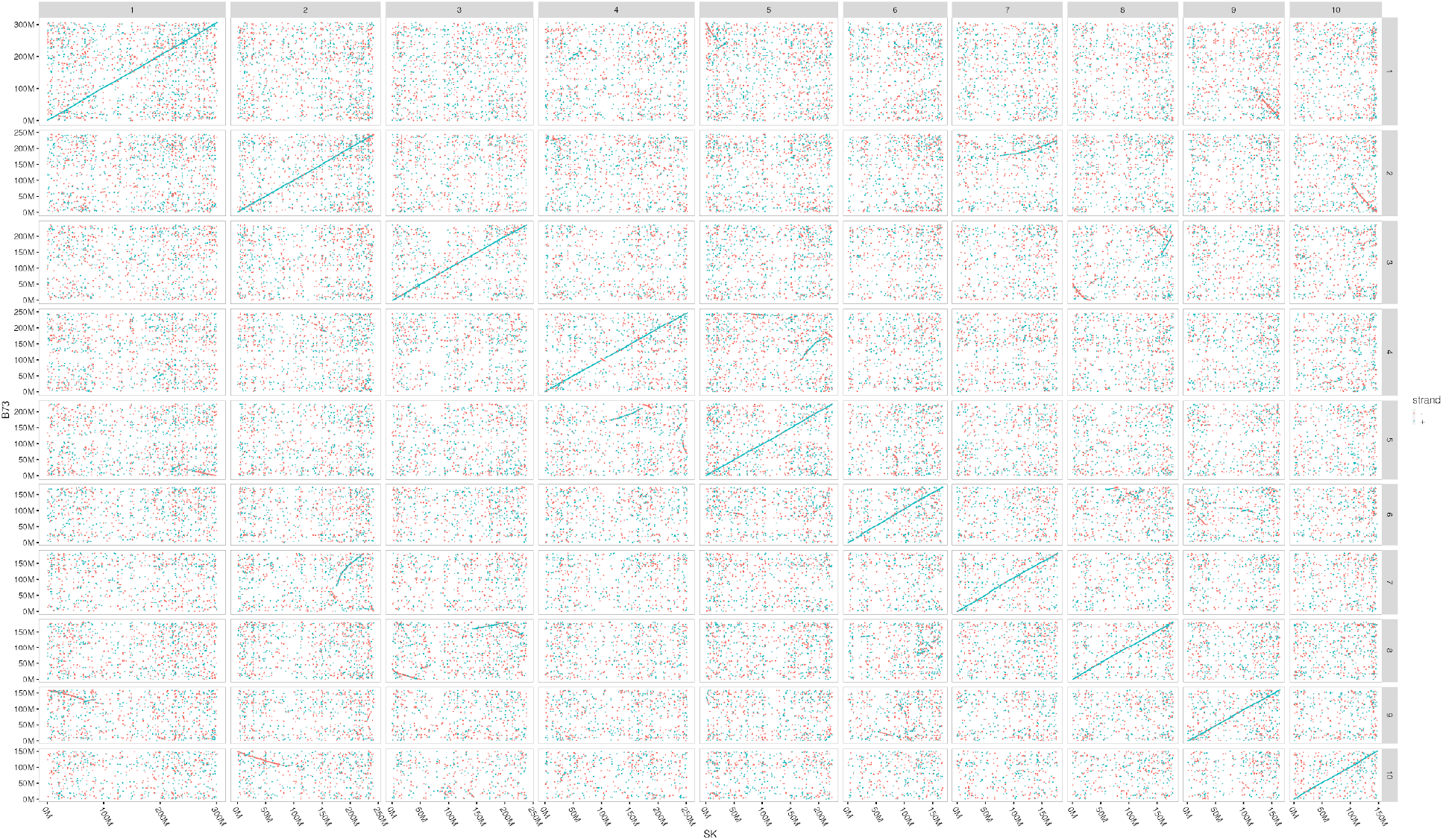
The anchor matches between the maize B73 V4 genome assembly and the SK assembly. Blue dots represent anchors on the same strand between the reference genome and the query genome. Red dots represent anchors on the different strands between the reference genome and the query genome.

We detected collinear anchors using the following command:

~~~
anchorwave genoAli -i Zea_mays.AGPv4.34.gff3 -r Zea_mays.AGPv4.dna.toplevel.fa -as cds.fa -a cds.sam -ar ref.sam -s GWHAACS00000000.genome.fasta -n anchorsiv -IV -ns
~~~

The collinear anchors in the output file (anchorsiv) were plotted for each chromosome separately.

**Figure S3.**
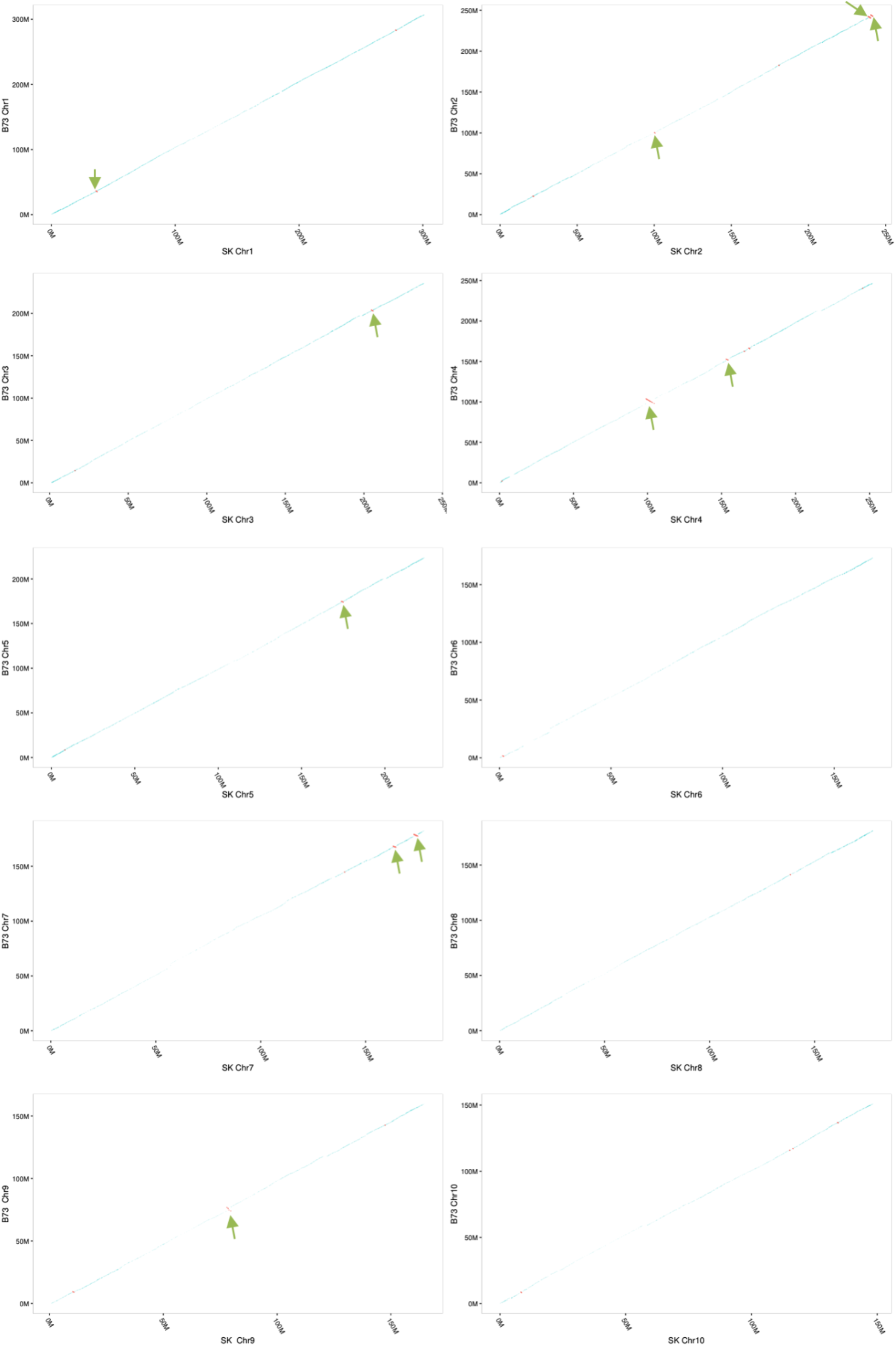
The identified collinear anchors between the B73 v4 assembly and the SK assembly. Those collinear anchors on the positive strand are colored with semi-transparent blue, and the negative anchors are colored with red. Previously reported inversions are labeled with arrows.

AnchorWave identified 11 out of 13 reported inversions(41). For two previously reported inversions, AnchorWave detected colinear gene anchors in the region, suggesting these regions may not be true inversions. The first uncalled inversion was previously reported in the range of Chr3:86.8 Mbp-88.3 Mbp of the B73 v4 assembly and Chr3:85.2 Mbp-87.0 Mbp of the SK assembly. We found 7 full-length CDS collinear anchors located in the range of Chr3:87.0 Mbp-88.1 Mbp of the B73 v4 assembly and Chr3:88.5 Mbp-89.6 Mbp of the SK assembly. These anchors are located on the same strand of the B73 assembly and the SK assembly. The second inversion was previously reported in the range of Chr4:168.3 Mbp-169.1 Mbp of the B73 v4 assembly and Chr4:165.1 Mbp-167.0 Mbp of the SK assembly. We found 7 full-length CDS collinear anchors located in the range of Chr4: 168.6 Mbp-169.0 Mbp of the B73 v4 assembly and Chr4:170.2 Mbp-170.7 Mbp of the SK assembly. Again, these anchors are located on the same strand in the B73 assembly and the SK assembly. We found an additional inversion in an adjacent upstream region of the B73 assembly (Chr4:165.9 Mbp-166.6 Mbp) and from Chr4:168.4 Mbp-169.0 Mbp in the SK assembly. These results indicate that using full-length CDSs as anchors provides enough sensitivity to capture long inversions.

**Supplementary Note 5:**

The common goldfish (*Carassius auratus*) experienced a whole-genome duplication since diverging from the common ancestor with zebrafish, and collinearity between the zebrafish genome assembly and the goldfish genome assembly exists(26). As the goldfish genome assembly size is comparable to that of zebrafish genome assembly(27), fractionation and loss of some homologous zebrafish genome fragments have occurred in goldfish. Each of the two subgenomes of goldfish is found in separate linkage groups; this absence of subgenome chromosome fusion makes the genome comparison more straightforward than the maize-sorghum genome comparison.

We downloaded the zebrafish genome assembly, genome annotation, and the goldfish genome assembly using the following commands:

~~~
wget ftp://ftp.ensembl.org/pub/release-102/fasta/danio_rerio/dna/Danio_rerio.GRCz11.dna.primary_assembly.fa.gz
wget ftp://ftp.ensembl.org/pub/release-102/gff3/danio_rerio/Danio_rerio.GRCz11.102.chr.gff3.gz
wget https://research.nhgri.nih.gov/goldfish/download/carAur01.sm.fa
gunzip *gz
~~~

Following previous work(26), to avoid heterozygous assembled regions, we used only linkage group assemblies when comparing genomes. The linkage group assemblies were extracted from the original FASTA file using the following command:

~~~
head -15923983 carAur01.sm.fa > goldfish.fa
~~~

We extracted the zebrafish full-length CDS and mapped full-length CDS to the zebrafish reference genome and goldfish query genome as introduced in Supplementary Note 1. The mapping matches are plotted in Figure S4.

**Figure S4.**
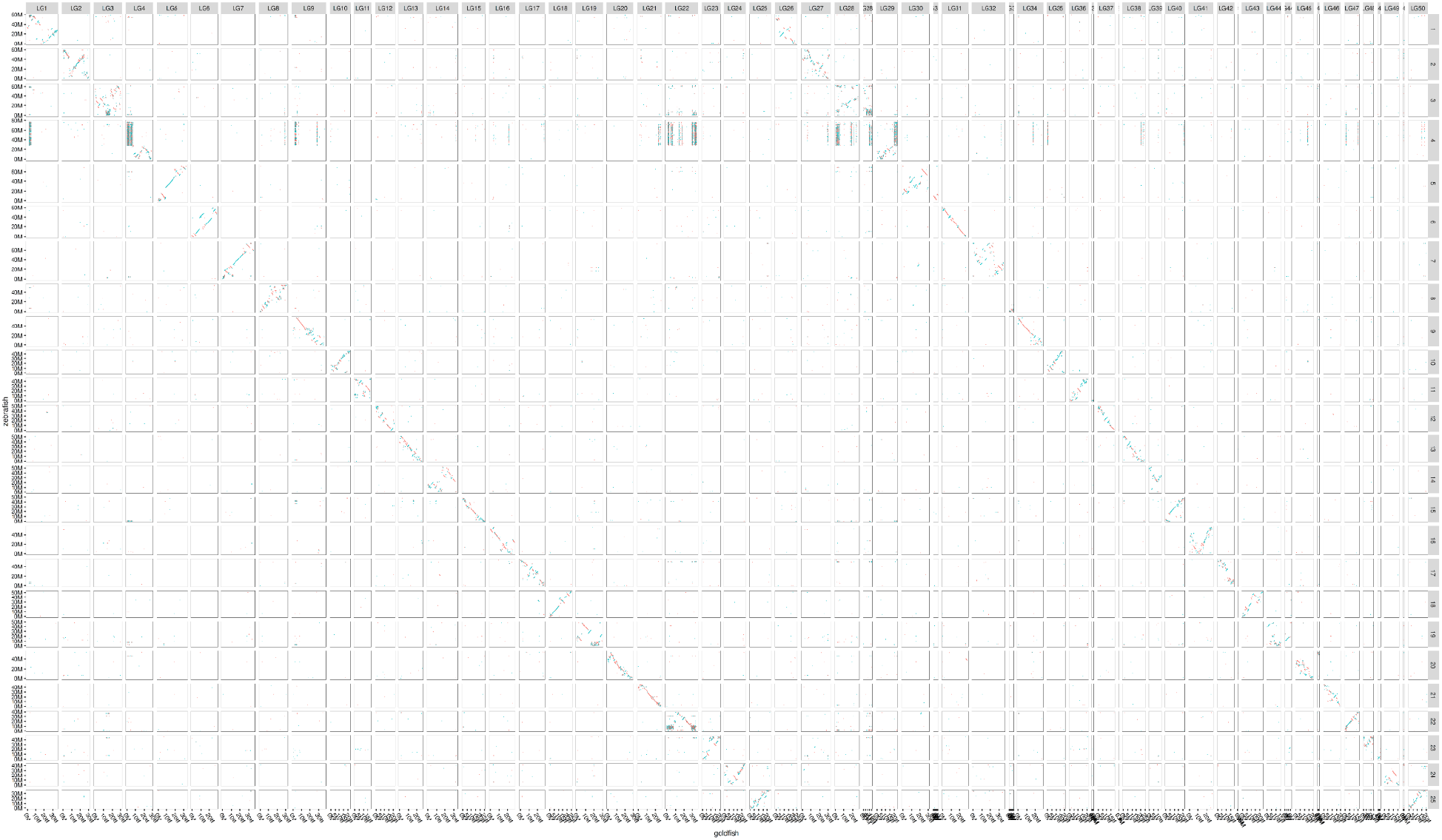
Anchor matches between the zebrafish genome and the goldfish genome. Blue points represent anchors on the same strand between the reference genome and the query genome. Red points represent anchors on different strands between the reference genome and the query genome.

We performed genome alignment using AnchorWave with command:

~~~
anchorwave proali -i Danio_rerio.GRCz11.102.chr.gff3 -r Danio_rerio.GRCz11.dna.primary_assembly.fa -a carAur.sam -as cds.fa - ar ref.sam -s goldfish.fa -n align1.anchors -R 2 -Q 1 -ns
~~~

Collinear anchors between the zebrafish genome and the goldfish genome are plotted in Figure S5:

**Figure S5.**
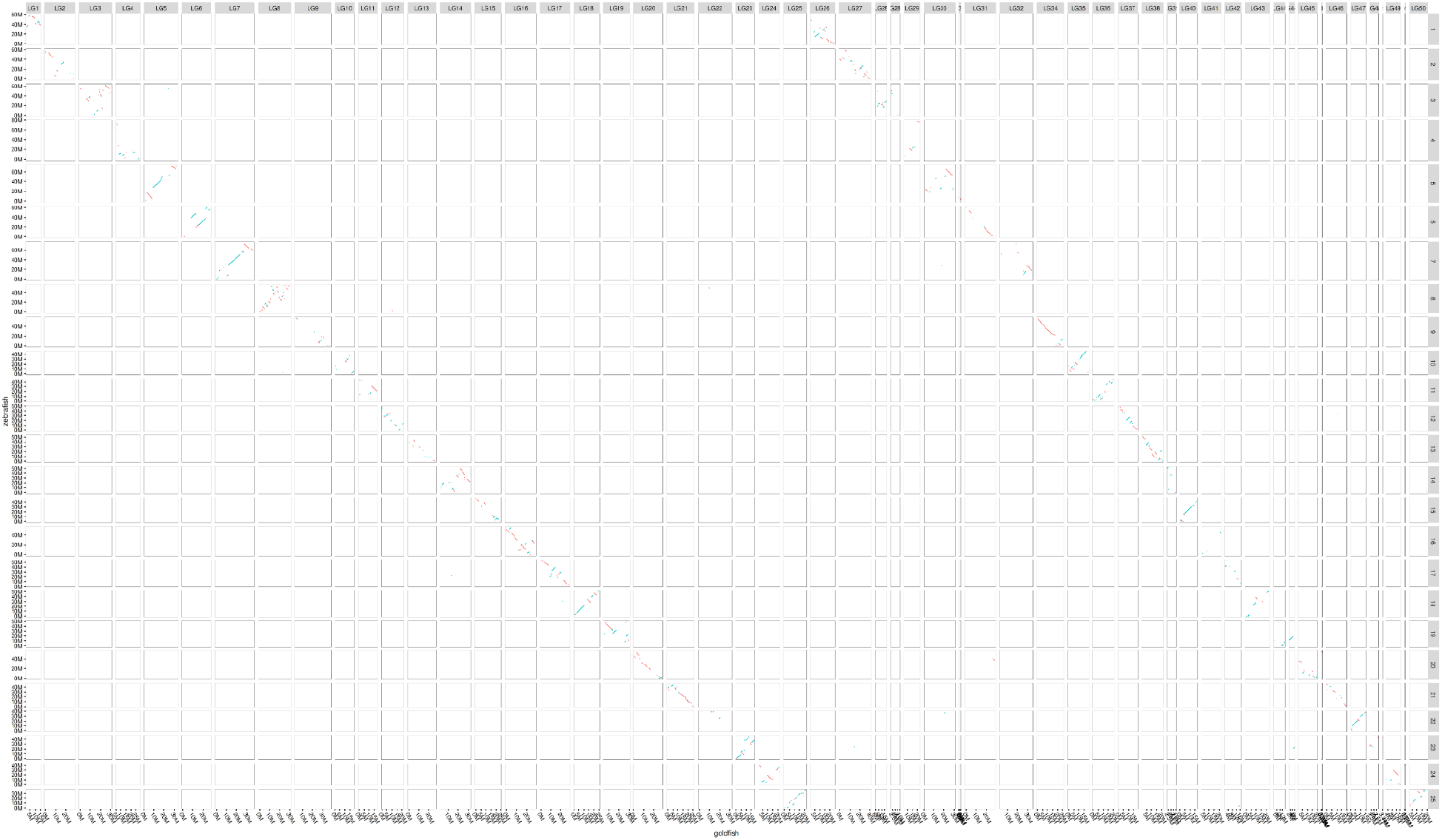
Cllinear anchors between the zebrafish genome assembly and the goldfish genome assembly. Blue points represent anchors on the same strand between the reference genome and the query genome. Red points represent anchors on different strands between the reference genome and the query genome.

We split the goldfish genome into two subgenomes via SeqKit(42) using commands:

~~~
seqkit grep -n -p LG1 goldfish.fa > goldfish1.fa
seqkit grep -n -p LG2 goldfish.fa >> goldfish1.fa
seqkit grep -n -p LG3 goldfish.fa >> goldfish1.fa
seqkit grep -n -p LG4 goldfish.fa >> goldfish1.fa
seqkit grep -n -p LG5 goldfish.fa >> goldfish1.fa
seqkit grep -n -p LG6 goldfish.fa >> goldfish1.fa
seqkit grep -n -p LG7 goldfish.fa >> goldfish1.fa
seqkit grep -n -p LG8 goldfish.fa >> goldfish1.fa
seqkit grep -n -p LG9 goldfish.fa >> goldfish1.fa
seqkit grep -n -p LG10 goldfish.fa >> goldfish1.fa
seqkit grep -n -p LG11 goldfish.fa >> goldfish1.fa
seqkit grep -n -p LG12 goldfish.fa >> goldfish1.fa
seqkit grep -n -p LG13 goldfish.fa >> goldfish1.fa
seqkit grep -n -p LG14 goldfish.fa >> goldfish1.fa
seqkit grep -n -p LG15 goldfish.fa >> goldfish1.fa
seqkit grep -n -p LG16 goldfish.fa >> goldfish1.fa
seqkit grep -n -p LG17 goldfish.fa >> goldfish1.fa
seqkit grep -n -p LG18 goldfish.fa >> goldfish1.fa
seqkit grep -n -p LG19 goldfish.fa >> goldfish1.fa
seqkit grep -n -p LG20 goldfish.fa >> goldfish1.fa
seqkit grep -n -p LG21 goldfish.fa >> goldfish1.fa
seqkit grep -n -p LG22 goldfish.fa >> goldfish1.fa
seqkit grep -n -p LG23 goldfish.fa >> goldfish1.fa
seqkit grep -n -p LG24 goldfish.fa >> goldfish1.fa
seqkit grep -n -p LG25 goldfish.fa >> goldfish1.fa
seqkit grep -n -p LG26 goldfish.fa > goldfish2.fa
seqkit grep -n -p LG27 goldfish.fa >> goldfish2.fa
seqkit grep -n -p LG28 goldfish.fa >> goldfish2.fa
seqkit grep -n -p LG28B goldfish.fa >> goldfish2.fa
seqkit grep -n -p LG29 goldfish.fa >> goldfish2.fa
seqkit grep -n -p LG30 goldfish.fa >> goldfish2.fa
seqkit grep -n -p LG30F goldfish.fa >> goldfish2.fa
seqkit grep -n -p LG31 goldfish.fa >> goldfish2.fa
seqkit grep -n -p LG32 goldfish.fa >> goldfish2.fa
seqkit grep -n -p LG33 goldfish.fa >> goldfish2.fa
seqkit grep -n -p LG34 goldfish.fa >> goldfish2.fa
seqkit grep -n -p LG35 goldfish.fa >> goldfish2.fa
seqkit grep -n -p LG36 goldfish.fa >> goldfish2.fa
seqkit grep -n -p LG36F goldfish.fa >> goldfish2.fa
seqkit grep -n -p LG37 goldfish.fa >> goldfish2.fa
seqkit grep -n -p LG37M goldfish.fa >> goldfish2.fa
seqkit grep -n -p LG38 goldfish.fa >> goldfish2.fa
seqkit grep -n -p LG39 goldfish.fa >> goldfish2.fa
seqkit grep -n -p LG40 goldfish.fa >> goldfish2.fa
seqkit grep -n -p LG41 goldfish.fa >> goldfish2.fa
seqkit grep -n -p LG42 goldfish.fa >> goldfish2.fa
seqkit grep -n -p LG42F goldfish.fa >> goldfish2.fa
seqkit grep -n -p LG43 goldfish.fa >> goldfish2.fa
seqkit grep -n -p LG44 goldfish.fa >> goldfish2.fa
seqkit grep -n -p LG44F goldfish.fa >> goldfish2.fa
seqkit grep -n -p LG45 goldfish.fa >> goldfish2.fa
seqkit grep -n -p LG45M goldfish.fa >> goldfish2.fa
seqkit grep -n -p LG46 goldfish.fa >> goldfish2.fa
seqkit grep -n -p LG47 goldfish.fa >> goldfish2.fa
seqkit grep -n -p LG48 goldfish.fa >> goldfish2.fa
seqkit grep -n -p LG48F goldfish.fa >> goldfish2.fa
seqkit grep -n -p LG49 goldfish.fa >> goldfish2.fa
seqkit grep -n -p LG49B goldfish.fa >> goldfish2.fa
seqkit grep -n -p LG50 goldfish.fa >> goldfish2.fa
~~~

We aligned the two subgenomes of goldfish separately against the zebrafish genome using LAST, minimap2, MUMmer4, and GSAlign with the same parameters as described in Supplementary Note 1. The minimap2 and MUMmer4 genome alignments were reformatted into MAF format using the “anchorwave sam2maf” function. The minimap2, MUMmer4, and GSalign results were filtered to one-to-one alignments using the “last-split | maf-swap | last-split | maf-swap” as described in Supplementary Note 1. We then merged the alignment of the two subgenomes together. The proportion of the zebrafish genome aligned as position match and gap are plotted in Figure S6.

**Figure S6.**
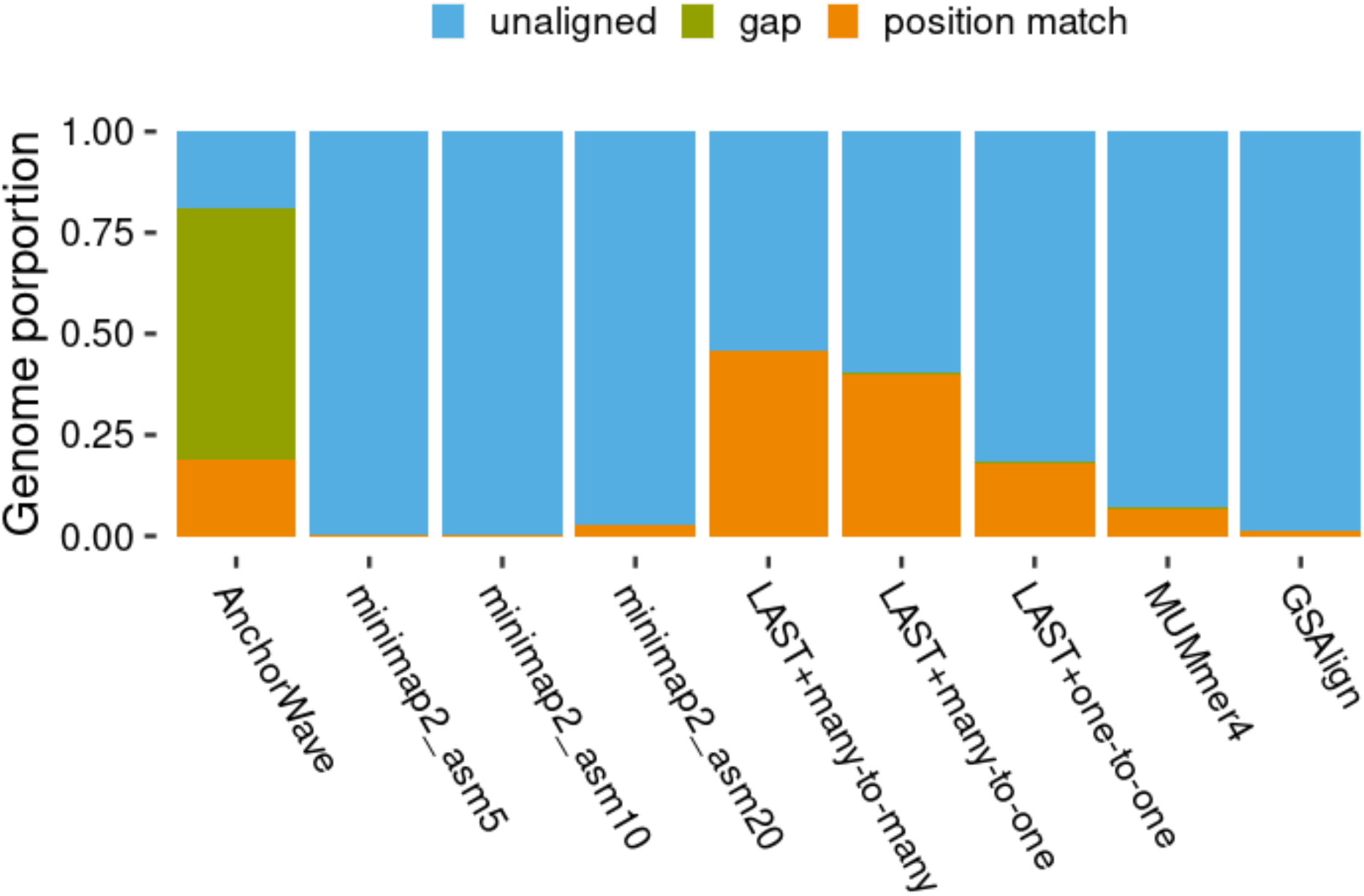
Comparison of the proportion of sites in the zebrafish genome aligned to the goldfish genome using AnchorWave and other genome alignment tools.

**Supplementary Note 6:**

The maize B73 v4 genome sequence and genome annotation were obtained as introduced in Supplementary Note 2, and the sorghum genome(43) was downloaded via the commands:

~~~
wget ftp://ftp.ensemblgenomes.org/pub/plants/release-49/fasta/sorghum_bicolor/dna/Sorghum_bicolor.Sorghum_bicolor_NCBIv3.dna.toplevel.fa.gz
gunzip Sorghum_bicolor.Sorghum_bicolor_NCBIv3.dna.toplevel.fa.gz
~~~

The reference full-length CDSs from maize were extracted and mapped to the reference genome and query genome as introduced in Supplementary Note 1.

The genome alignment using AnchorWave was conducted via:

~~~
anchorwave proali -i Zea_mays.AGPv4.34.gff3 -as cds.fa -r Zea_mays.AGPv4.dna.toplevel.fa -a cds.sam -ar ref.sam -s Sorghum_bicolor.Sorghum_bicolor_NCBIv3.dna.toplevel.fa -n anchors -mi 0 -R 1 -Q 2 -o alignment.maf -f alignment.f.maf -w 38000 -fa3 200000
~~~

The alignments was reformatted into bam format:

~~~
maf-convert sam anchorwave.maf | sed ‘s/Sorghum_bicolor.Sorghum_bicolor_NCBIv3.dna.toplevel.fa.//g’ | sed ‘s/Zea_mays.AGPv4.dna.toplevel.fa.//g’ | samtools view -O BAM --reference Zea_mays.AGPv4.dna.toplevel.fa - | samtools sort - > anchorwave.bam
~~~

The sequence alignments using LAST, MUMmer4, minimap2, and GSAlign were conducted with the same settings as introduced in Supplementary Note 1.

Fig. S5 illustrates how we define position match sites, aligned sites, gap sites, and unaligned sites on the reference genome in this study.

We counted the number of position match sites using the command:

~~~
samtools depth alignment.bam | awk ‘$3>0 {print $0}’ | wc -l
~~~

We counted the number of aligned sites using the the command:

~~~
samtools depth alignment.bam | wc -l
~~~

The number of gap sites is calculated as the number of aligned sites minus the number of position match sites, and the number of unaligned sites is the maize genome size minus the number of aligned sites.

The maize TFBS annotation (all_reproducible_peaks_summits_merged.bed) is available as “Supplementary Data 5” from a previous publication(7). We used the following commands to count the number of reference sites being aligned, and those aligned as position match in maize TFBS regions:

~~~
samtools depth alignment.bam -b all_reproducible_peaks_summits_merged.bed | wc -l
samtools depth alignment.bam -b all_reproducible_peaks_summits_merged.bed | awk ‘$3>0{print $0}’ | wc -l
~~~

The maize TE annotation file has been released at: https://github.com/mcstitzer/maize_TEs/blob/master/B73.structuralTEv2.fulllength.2018-09-19.gff3.gz. We use the following command to transform it into bed format:

~~~
grep -v “#” B73.structuralTEv2.disjoined.2018-09-19.gff3 | awk ‘{print $1”\t”$4-1”\t”$5}’ > B73.structuralTEv2.disjoined.2018-09-19.bed
~~~

We used the following commands to count, in maize TE regions, the number of reference sites being aligned and aligned as position matches:

~~~
samtools depth alignment.bam -b B73.structuralTEv2.disjoined.2018-09-19.bed | wc -l
samtools depth alignment.bam -b B73.structuralTEv2.disjoined.2018-09-19.bed | awk ‘$3>0{print $0}’ | wc -l
~~~

**Supplementary Note 7:**

Alignment with AnchorWave relies on the presence of collinear blocks between species. We investigated the power of AnchorWave for aligning species with different levels of divergence. We selected several species to span different phylogenetic distances and rounds of wholegenome duplication as illustrated at: https://genomevolution.org/wiki/index.php/Whole_genome_duplication.

Maize and rice are among the most divergent grass species with available genome assemblies.. We downloaded the rice genome using commands:

~~~
wget ftp://ftp.ensemblgenomes.org/pub/plants/release-49/fasta/oryza_sativa/dna/Oryza_sativa.IRGSP-1.0.dna.toplevel.fa.gzgunzipOryza_sativa.IRGSP-1.0.dna.toplevel.fa.gz
~~~

We mapped maize full-length CDS sequence using minimap2 and identified collinear blocks using AnchorWave:

~~~
minimap2 -x splice -t 11 -k 12 -a -p 0.4 -N 20 Oryza_sativa.IRGSP-1.0.dna.toplevel.fa maize_cds.fa > maize.rice.sam
anchorwave proali -i Zea_mays.AGPv4.34.gff3 -r Zea_mays.AGPv4.dna.toplevel.fa -as maize_cds.fa -a maize.rice.sam -ar maize_cds_maize.sam -s Oryza_sativa.IRGSP-1.0.dna.toplevel.fa -n B73_rice.anchorspro -ns -R 1 -Q 2
~~~

1.82 Gbp of the maize genome was identified in collinear blocks with the rice genome.

**Figure S7.**
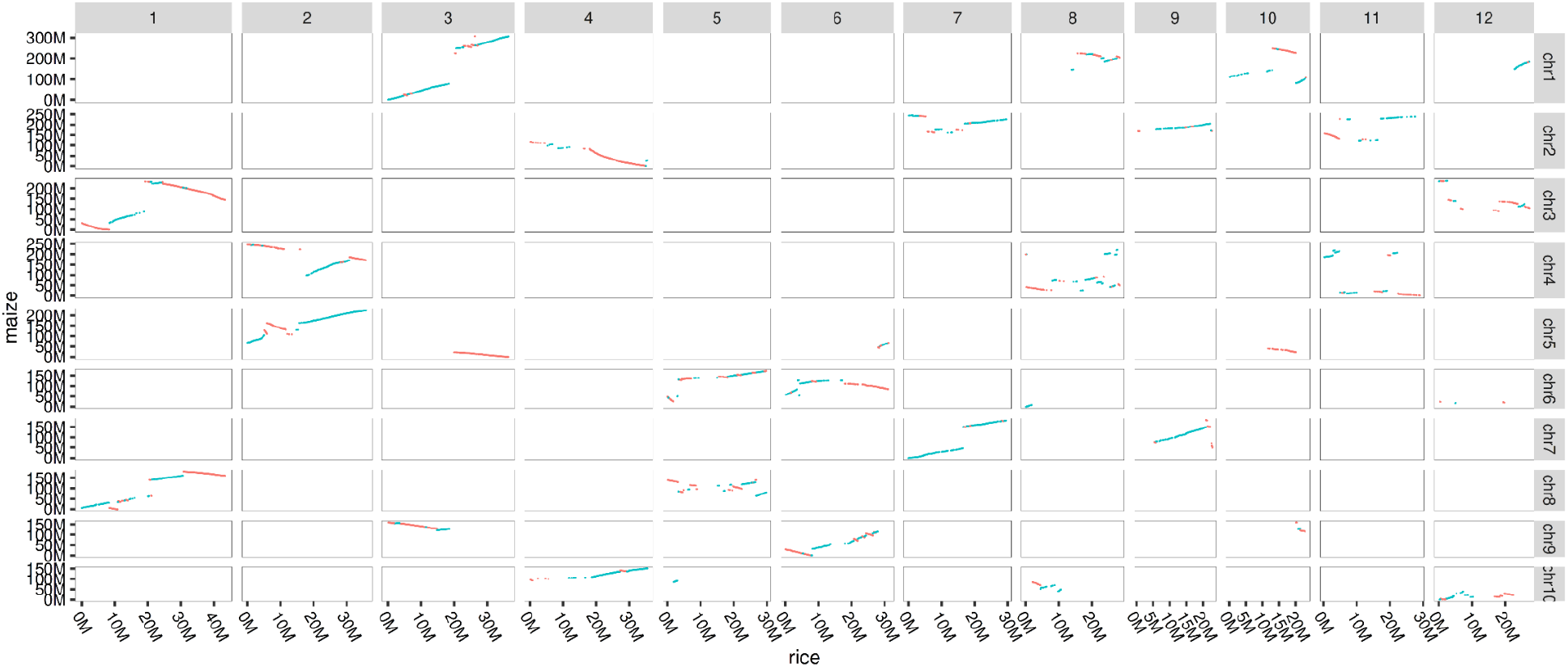
Collinear anchors between maize B73 v4 genome assembly and the rice genome assembly. Blue points represent anchors on the same strand between the reference genome and the query genome. Red points represent anchors on different strands between the reference genome and the query genome.

There have been six rounds of whole-genome duplication since maize and banana diverged. We downloaded the banana genome and aligned the banana genome against the maize genome.

~~~
wget https://www.genoscope.cns.fr/externe/plants/data/Mschizocarpa_chromosomes.fasta # banana minimap2 -x splice -t 11 -k 12 -a -p 0.4 -N 20 Mschizocarpa_chromosomes.fasta maize_cds.fa > maize.banana.sam
anchorwave proali -i Zea_mays.AGPv4.34.gff3 -r Zea_mays.AGPv4.dna.toplevel.fa -as maize_cds.fa -a maize.banana.sam -ar maize_cds_maize.sam -s Mschizocarpa_chromosomes.fasta maize.cds.fa -n B73_banana.anchorspro -ns -R 8 -Q 8
~~~

**Figure S8.**
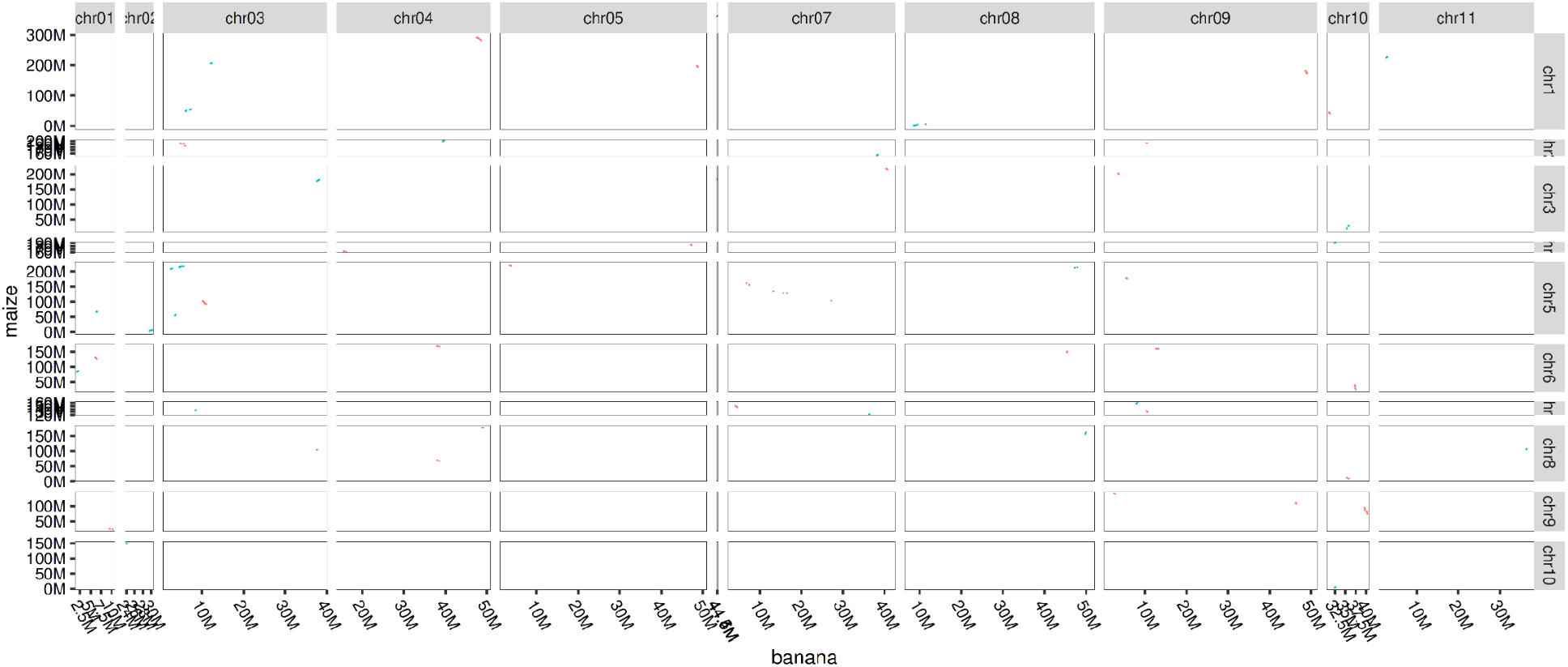
Collinear anchors between the maize B73 v4 genome assembly and the banana genome assembly. Blue points represent anchors on the same strand between the reference genome and the query genome. Red points represent anchors on different strands between the reference genome and the query genome.

The only collinear blocks between maize and banana were short, thus AnchorWave genome alignment between maize and banana did not recall homologous blocks. This represents an upper bound for divergence via whole-genome duplication between species for applying AnchorWave.

The divergence time between *Arabidopsis thaliana* and chocolate is comparable to that between maize and banana, but the species are only separated by two rounds of whole-genome duplication. We downloaded the chocolate genome and aligned the chocolate genome against the *A. thaliana* genome.

~~~
wget ftp://ftp.ensemblgenomes.org/pub/plants/release-49/fasta/theobroma_cacao_matina/dna/Theobroma_cacao_matina.Theobroma_cacao_20110822.dna.toplevel.fa.gz
gunzip Theobroma_cacao_matina.Theobroma_cacao_20110822.dna.toplevel.fa.gz
minimap2 -x splice -t 11 -k 12 -a -p 0.4 -N 20 Theobroma_cacao_matina.Theobroma_cacao_20110822.dna.toplevel.fa tair10_cds.fa > ara.chocolate.sam
~~~

**Figure S9.**
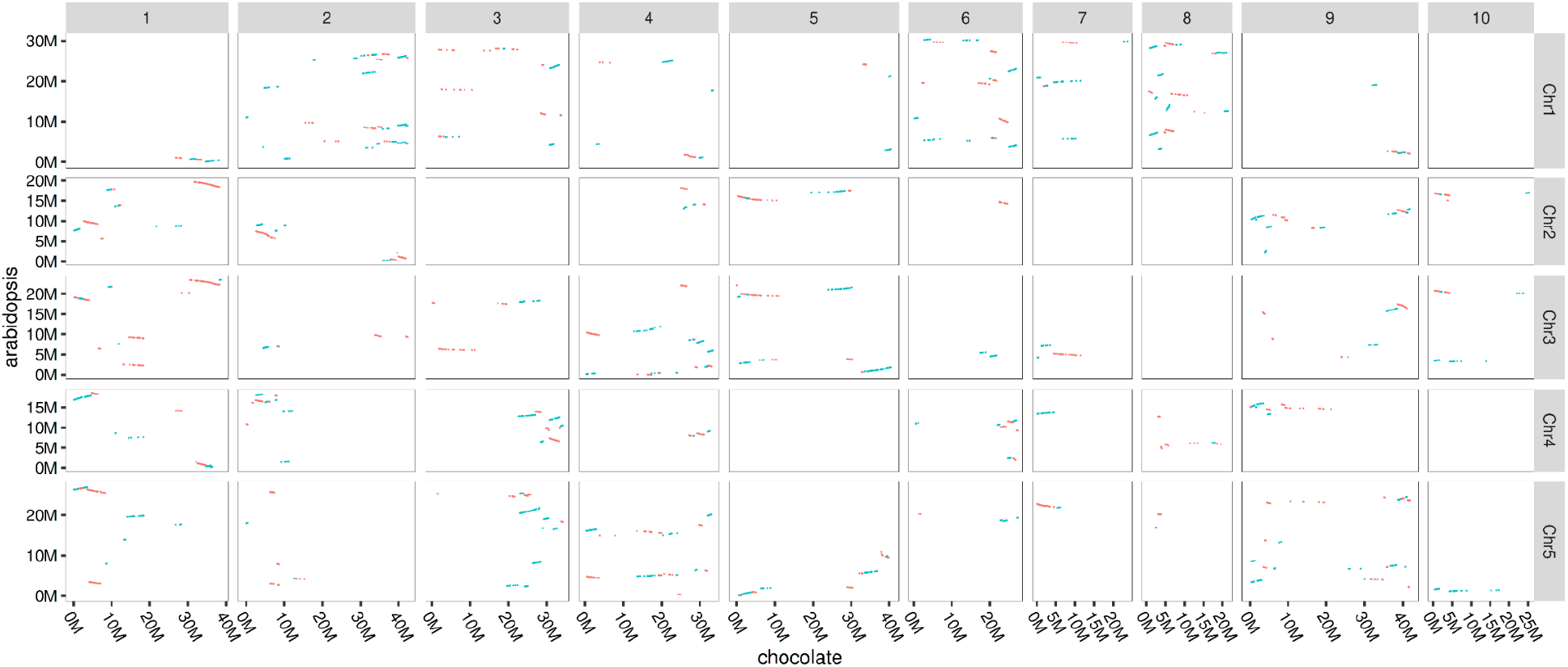
Collinear anchors between the Arabidopsis TAIR10 genome assembly and the chocolate genome assembly. Blue points represent anchors on the same strand between the reference genome and the query genome. Red points represent anchors on different strands between the reference genome and the query genome.

There are more collinear blocks found between the *A. thaliana* genome assembly and the chocolate genome assembly than between the maize genome assembly and the banana assembly. This shows that whole-genome duplications and subsequent decay and fractionation may be one of the key mechanisms that break collinearity.

There are the same number of whole-genome duplications between *A. thaliana* and grape as there are between *A. thaliana* and chocolate. However, the divergence time between *A. thaliana* and grape is longer than that between *A. thaliana* and chocolate. We downloaded the grape genome and aligned the grape genome against the *A. thaliana* genome.

~~~
wget ftp://ftp.ensemblgenomes.org/pub/plants/release-49/fasta/vitis_vinifera/dna/Vitis_vinifera.12X.dna.toplevel.fa.gz minimap2 -x splice -t 11 -k 12 -a -p 0.4 -N 20 Vitis_vinifera.12X.dna.toplevel.fa tair10_cds.fa > ara.grape.sam anchorwave proali -i TAIR10_GFF3_genes.gff -r tair10.fa -as tair10_cds.fa -a ara.grape.sam -ar tair10.sam -s Vitis_vinifera.12X.dna.toplevel.fa -n ara_grape.anchorspro -ns -R 1 -Q 4
~~~

**Figure S10.**
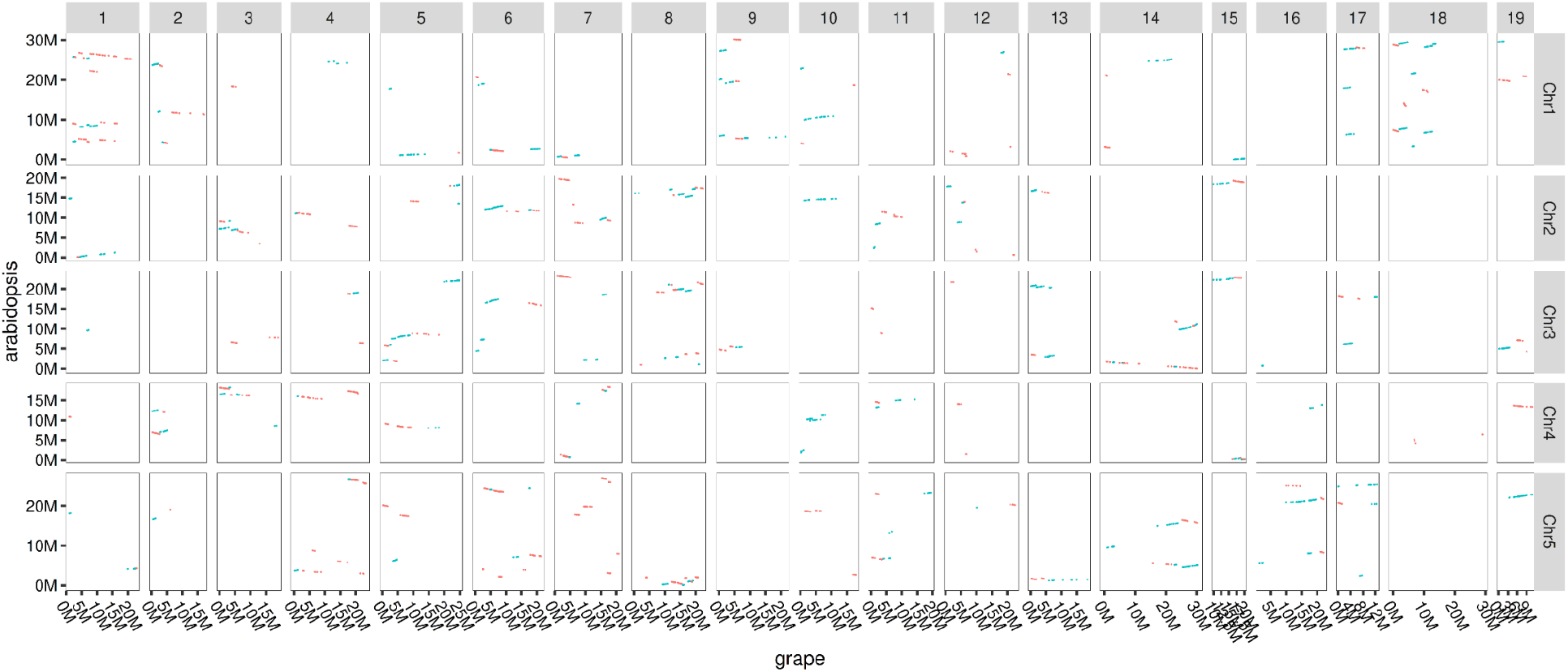
Collinear anchors between the Arabidopsis TAIR10 genome assembly and the grape genome assembly. Blue points represent anchors on the same strand between the reference genome and the query genome. Those red dots represent anchors on the different strands between the reference genome and the query genome.

Fewer *A. thaliana* base pairs are found in collinear blocks with the grape genome than with the chocolate genome. Collinearity may decay along with genomic divergence and reduce the applicability of AnchorWave for genome alignment.

A guideline document to check collinearity and select appropriate parameters of “-Q” and “-R” can be found under the AnchorWave source code repository.

**Supplementary Note 8:**

We test AnchorWave for collinear block identification between the human genome (hg38) and the house mouse (*Mus musculus*) genome (mm39).

We downloaded the mouse genome using the following commands:

~~~
wget https://hgdownload.soe.ucsc.edu/goldenPath/mm39/chromosomes/chr1.fa.gz
wget https://hgdownload.soe.ucsc.edu/goldenPath/mm39/chromosomes/chr2.fa.gz
wget https://hgdownload.soe.ucsc.edu/goldenPath/mm39/chromosomes/chr3.fa.gz
wget https://hgdownload.soe.ucsc.edu/goldenPath/mm39/chromosomes/chr4.fa.gz
wget https://hgdownload.soe.ucsc.edu/goldenPath/mm39/chromosomes/chr5.fa.gz
wget https://hgdownload.soe.ucsc.edu/goldenPath/mm39/chromosomes/chr6.fa.gz
wget https://hgdownload.soe.ucsc.edu/goldenPath/mm39/chromosomes/chr7.fa.gz
wget https://hgdownload.soe.ucsc.edu/goldenPath/mm39/chromosomes/chr8.fa.gz
wget https://hgdownload.soe.ucsc.edu/goldenPath/mm39/chromosomes/chr9.fa.gz
wget https://hgdownload.soe.ucsc.edu/goldenPath/mm39/chromosomes/chr10.fa.gz
wget https://hgdownload.soe.ucsc.edu/goldenPath/mm39/chromosomes/chr11.fa.gz
wget https://hgdownload.soe.ucsc.edu/goldenPath/mm39/chromosomes/chr12.fa.gz
wget https://hgdownload.soe.ucsc.edu/goldenPath/mm39/chromosomes/chr13.fa.gz
wget https://hgdownload.soe.ucsc.edu/goldenPath/mm39/chromosomes/chr14.fa.gz
wget https://hgdownload.soe.ucsc.edu/goldenPath/mm39/chromosomes/chr15.fa.gz
wget https://hgdownload.soe.ucsc.edu/goldenPath/mm39/chromosomes/chr16.fa.gz
wget https://hgdownload.soe.ucsc.edu/goldenPath/mm39/chromosomes/chr17.fa.gz
wget https://hgdownload.soe.ucsc.edu/goldenPath/mm39/chromosomes/chr18.fa.gz
wget https://hgdownload.soe.ucsc.edu/goldenPath/mm39/chromosomes/chr19.fa.gz
wget https://hgdownload.soe.ucsc.edu/goldenPath/mm39/chromosomes/chrM.fa.gz
wget https://hgdownload.soe.ucsc.edu/goldenPath/mm39/chromosomes/chrX.fa.gz
gunzip *gz
cat chr*fa > mm39.fa
~~~

We downloaded the human genome annotation using the following commands:

~~~
wget ftp://ftp.ensembl.org/pub/release-102/gff3/homo_sapiens/Homo_sapiens.GRCh38.102.gff3.gz
~~~

We downloaded the human genome using the following commands:

~~~
wget ftp://ftp.ensembl.org/pub/release-102/fasta/homo_sapiens/dna/Homo_sapiens.GRCh38.dna.chromosome.1.fa.gz
wget ftp://ftp.ensembl.org/pub/release-102/fasta/homo_sapiens/dna/Homo_sapiens.GRCh38.dna.chromosome.2.fa.gz
wget ftp://ftp.ensembl.org/pub/release-102/fasta/homo_sapiens/dna/Homo_sapiens.GRCh38.dna.chromosome.3.fa.gz
wget ftp://ftp.ensembl.org/pub/release-102/fasta/homo_sapiens/dna/Homo_sapiens.GRCh38.dna.chromosome.4.fa.gz
wget ftp://ftp.ensembl.org/pub/release-102/fasta/homo_sapiens/dna/Homo_sapiens.GRCh38.dna.chromosome.5.fa.gz
wget ftp://ftp.ensembl.org/pub/release-102/fasta/homo_sapiens/dna/Homo_sapiens.GRCh38.dna.chromosome.6.fa.gz
wget ftp://ftp.ensembl.org/pub/release-102/fasta/homo_sapiens/dna/Homo_sapiens.GRCh38.dna.chromosome.7.fa.gz
wget ftp://ftp.ensembl.org/pub/release-102/fasta/homo_sapiens/dna/Homo_sapiens.GRCh38.dna.chromosome.8.fa.gz
wget ftp://ftp.ensembl.org/pub/release-102/fasta/homo_sapiens/dna/Homo_sapiens.GRCh38.dna.chromosome.9.fa.gz
wget ftp://ftp.ensembl.org/pub/release-102/fasta/homo_sapiens/dna/Homo_sapiens.GRCh38.dna.chromosome.10.fa.gz
wget ftp://ftp.ensembl.org/pub/release-102/fasta/homo_sapiens/dna/Homo_sapiens.GRCh38.dna.chromosome.11.fa.gz
wget ftp://ftp.ensembl.org/pub/release-102/fasta/homo_sapiens/dna/Homo_sapiens.GRCh38.dna.chromosome.12.fa.gz
wget ftp://ftp.ensembl.org/pub/release-102/fasta/homo_sapiens/dna/Homo_sapiens.GRCh38.dna.chromosome.13.fa.gz
wget ftp://ftp.ensembl.org/pub/release-102/fasta/homo_sapiens/dna/Homo_sapiens.GRCh38.dna.chromosome.14.fa.gz
wget ftp://ftp.ensembl.org/pub/release-102/fasta/homo_sapiens/dna/Homo_sapiens.GRCh38.dna.chromosome.15.fa.gz
wget ftp://ftp.ensembl.org/pub/release-102/fasta/homo_sapiens/dna/Homo_sapiens.GRCh38.dna.chromosome.16.fa.gz
wget ftp://ftp.ensembl.org/pub/release-102/fasta/homo_sapiens/dna/Homo_sapiens.GRCh38.dna.chromosome.17.fa.gz
wget ftp://ftp.ensembl.org/pub/release-102/fasta/homo_sapiens/dna/Homo_sapiens.GRCh38.dna.chromosome.18.fa.gz
wget ftp://ftp.ensembl.org/pub/release-102/fasta/homo_sapiens/dna/Homo_sapiens.GRCh38.dna.chromosome.19.fa.gz
wget ftp://ftp.ensembl.org/pub/release-102/fasta/homo_sapiens/dna/Homo_sapiens.GRCh38.dna.chromosome.20.fa.gz
wget ftp://ftp.ensembl.org/pub/release-102/fasta/homo_sapiens/dna/Homo_sapiens.GRCh38.dna.chromosome.21.fa.gz
wget ftp://ftp.ensembl.org/pub/release-102/fasta/homo_sapiens/dna/Homo_sapiens.GRCh38.dna.chromosome.22.fa.gz
wget ftp://ftp.ensembl.org/pub/release-102/fasta/homo_sapiens/dna/Homo_sapiens.GRCh38.dna.chromosome.X.fa.gz
gunzip *gz
cat Homo_sapiens.GRCh38.dna.chromosome.*.fa > hg38.fa
~~~

We extracted the human full-length CDS and mapped to the human genome and mouse genome separately using the following commands:

~~~
anchorwave gff2seq -r hg38.fa -i Homo_sapiens.GRCh38.102.gff3 -o cds.fa
minimap2 -x splice -t 10 -k 12 -a -p 0.4 -N 20 hg38.fa cds.fa > ref.sam
minimap2 -x splice -t 10 -k 12 -a -p 0.4 -N 20 mm39.fa cds.fa > cds.sam
~~~

The genome sequence alignment was conducted using the following command:

~~~
anchorwave proali -i Homo_sapiens.GRCh38.102.gff3 -r hg38.fa -a cds.sam -as cds.fa -ar ref.sam -s mm39.fa -n mm39.anchors -R 1 -Q 1 -o mm39.maf -f mm39.f.maf -w 38000 -fa3 200000 -B -4 -O1 -4 -E1 -2 -O2 -80 -E2 -1 -t 19
~~~

The collinear anchors are plotted in Figure S11.

**Figure S11.**
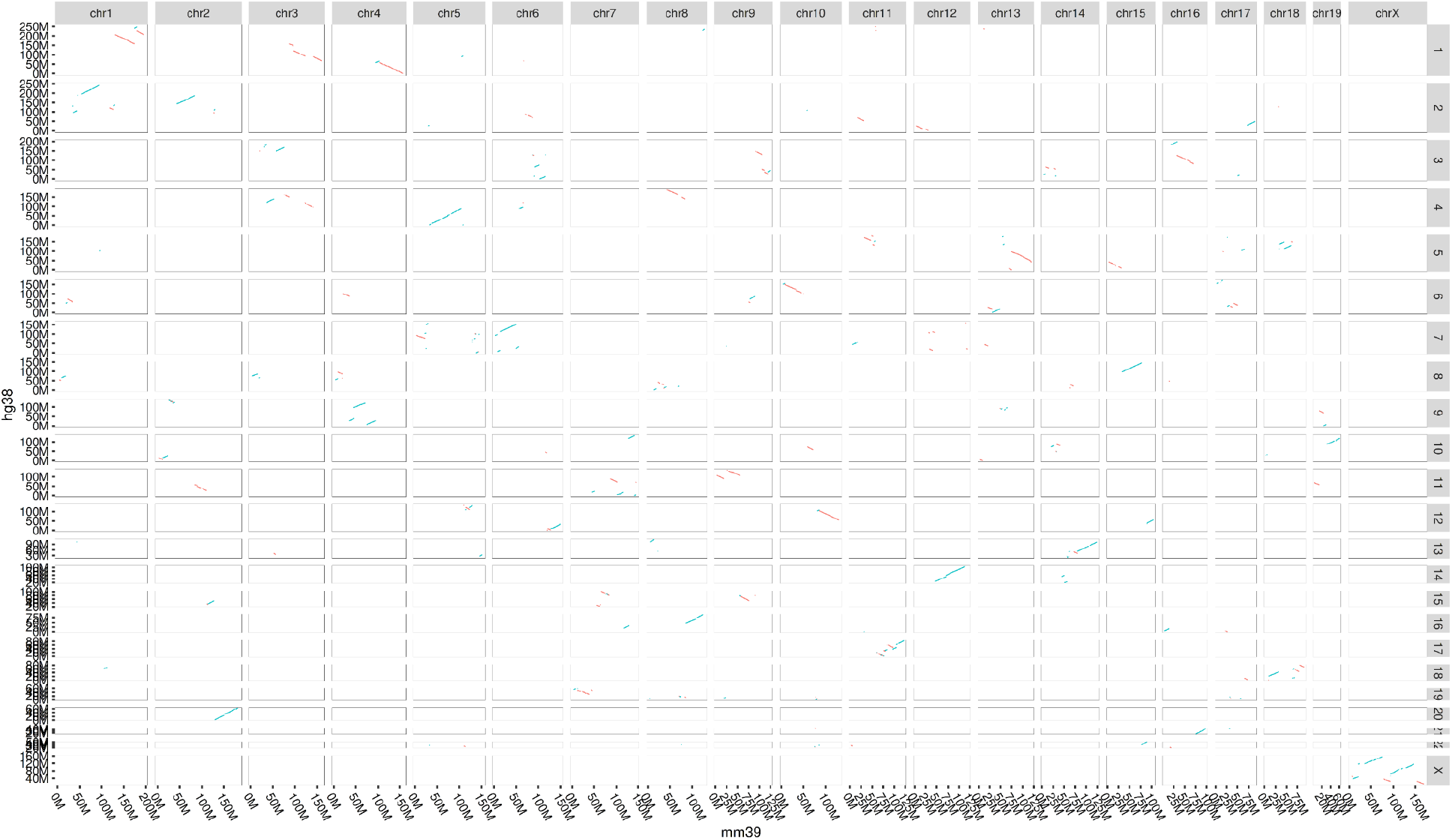
Collinear anchors between the human (hg38) genome assembly and mouse (mm39) genome assembly. Each point is plotted at its start position of each anchor on the reference (hg38) genome and the query genome (mm39). Blue points represent anchors on the same strand between the reference genome and the query genome. Red points represent anchors on different strands between the reference genome and the query genome.

On the autosomes and the X chromosome, 2.616Gbp (86.29%) of the human genome is aligned using AnchorWave. This is larger than that of commonly used chain and net (33.06%, 2.697Gbp) (Figure S12). The number of sites being aligned as position matches is comparable between these two methods (AnchorWave:892.1 Mbp, chain and net: 892.3 Mbp).

**Figure S12.**
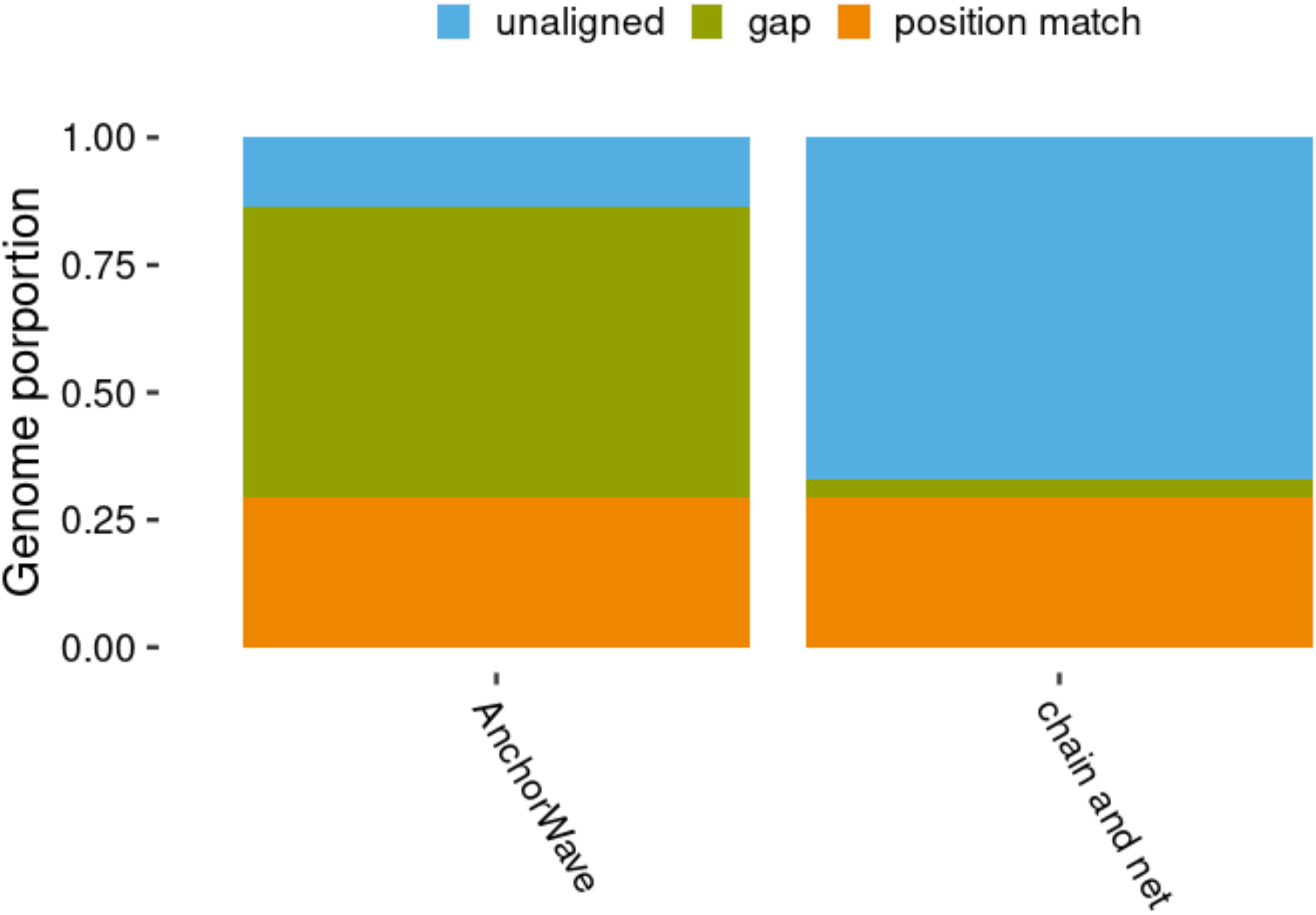
Comparison of the proportion of human (hg38) genome sites being aligned to mouse (mm39) using AnchorWave versus genome chain and net genome alignment. The chain and net genome alignment is downloaded from http://hgdownload.cse.ucsc.edu/goldenpath/hg38/vsMm39/hg38.mm39.synNet.maf.gz.

To identify overlap with regulatory sequence, we get ENCODE human candidate *cis*-regulatory elementsts via commands:

~~~
wget http://hgdownload.soe.ucsc.edu/gbdb/hg38/encode3/ccre/encodeCcreCombined.bb
bigBedToBed encodeCcreCombined.bb encodeCcreCombined.bed
sed -i -E ‘s/^chr//g’ encodeCcreCombined.bed
bedtools sort -i encodeCcreCombined.bed | bedtools merge | grep -v “^Y” > encodeCcreCombined_merged.bed
~~~

AnchorWave aligned 121.15 Mbp (48%) of *cis*-regulatory elements as position match, which is slightly lower than the downloaded chain and net alignment (131.68 Mbp 52%).

**Supplementary Note 9:**

We test AnchorWave for collinear block identification between the human genome (hg38) and the chimpanzee *(Pan troglodytes)* genome (panTro3).

We downloaded the chimpanzee genome using the following commands:

~~~
wget ftp://ftp.ensembl.org/pub/release-102/fasta/pan_troglodytes/dna/Pan_troglodytes.Pan_tro_3.0.dna.toplevel.fa.gz
~~~

We downloaded the human genome and genome annotation as described in Supplementary Note 8.

We extracted the human full-length CDS and mapped to the human genome and chimpanzee genome separately using the following commands:

~~~
anchorwave gff2seq -r hg38.fa -i Homo_sapiens.GRCh38.102.gff3 -o cds.fa
minimap2 -x splice -t 10 -k 12 -a -p 0.4 -N 20 hg38.fa cds.fa > ref.sam
minimap2 -x splice -t 10 -k 12 -a -p 0.4 -N 20 Pan_troglodytes.Pan_tro_3.0.dna.toplevel.fa cds.fa > cds.sam
~~~

The full-length CDS mapping is visualized as Figure S13, and the Y chromosome is shown in Figure S14. This pattern is consistent with previous reports of collinearity between human and chimpanzee(44).

**Figure S13.**
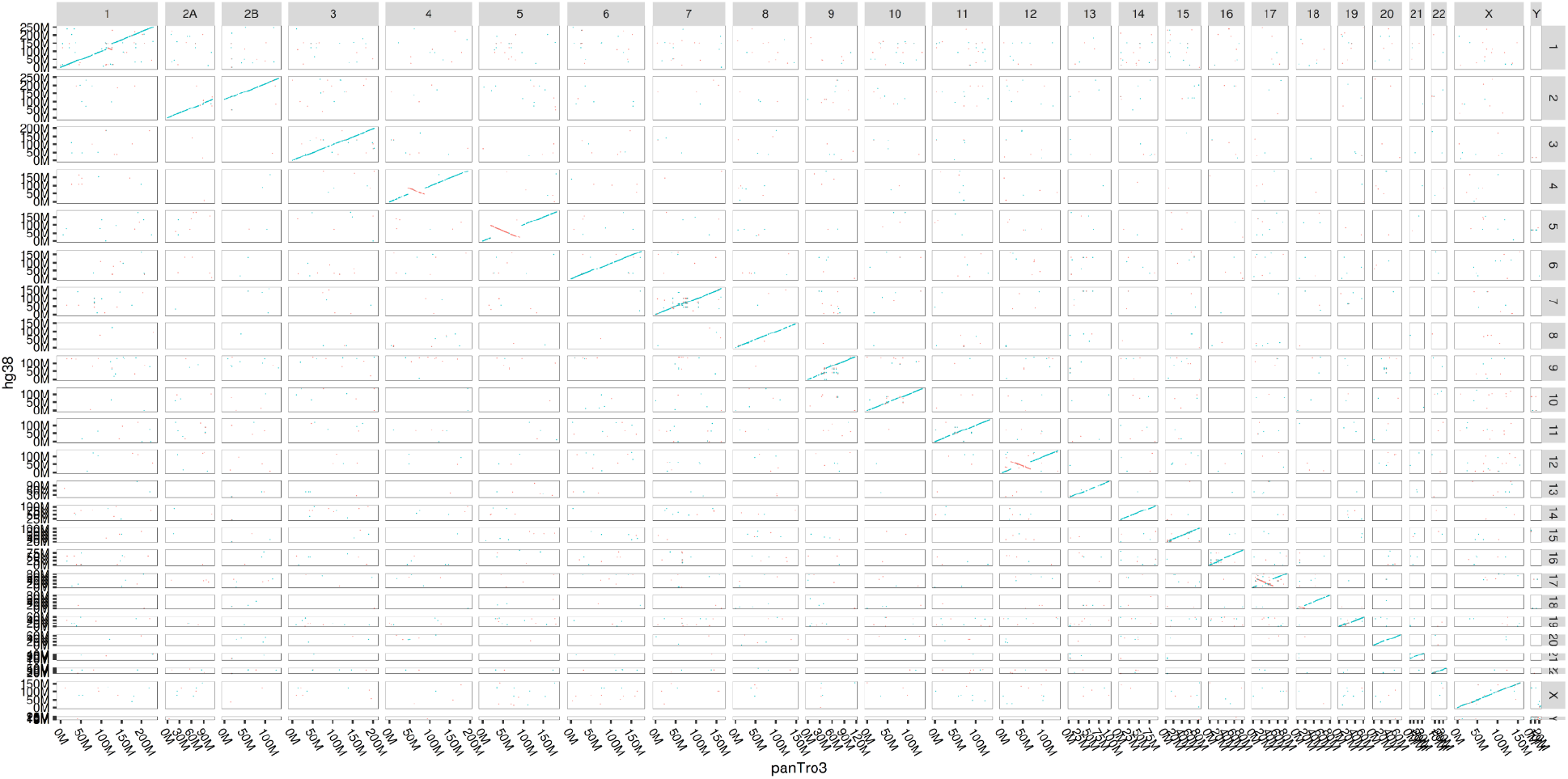
Anchor matches between the human (hg38) genome assembly and the chimpanzee (panTro3) genome assembly. Blue points represent anchors on the same strand between the reference genome and the query genome. Red points represent anchors on the different strands between the reference genome and the query genome.

**Figure S14.**
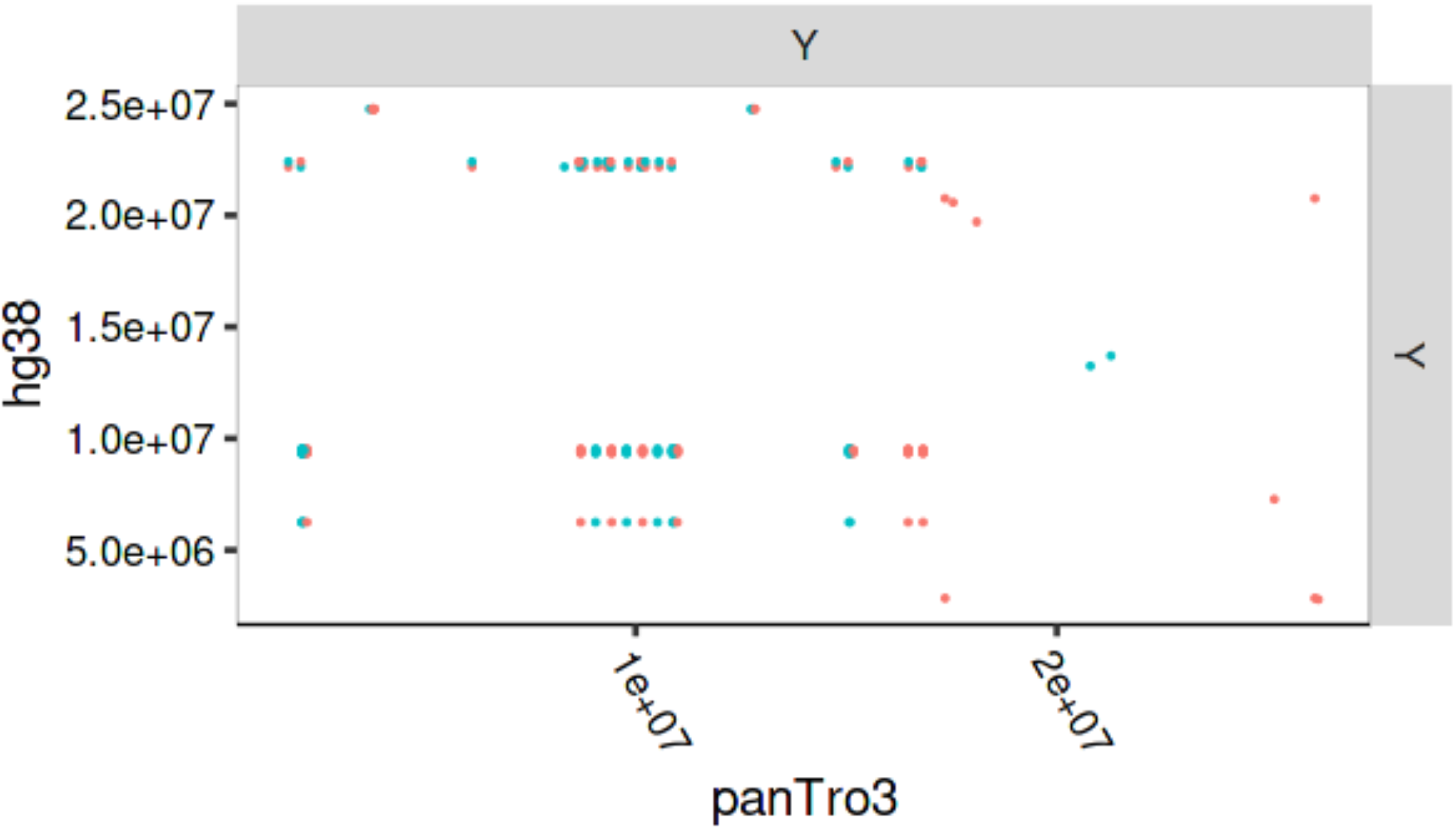
Anchor matches between the human (hg38) Y chromosome and the chimpanzee (panTro3) Y chromosome. Blue points represent anchors on the same strand between the reference genome and the query genome. Red points represent anchors on different strands between the reference genome and the query genome.

The genome sequence alignment was conducted using the following command:

~~~
anchorwave proali -i Homo_sapiens.GRCh38.102.gff3 -r hg38.fa -a cds.sam -as cds.fa -ar ref.sam -s Pan_troglodytes.Pan_tro_3.0.dna.toplevel.fa -n 2panTro3.anchors -R 1 -Q 1 -o 2panTro3.maf -f 2panTro3.f.maf -w 38000 -fa3 200000
~~~

The collinear anchors are plotted in Figure S15.

**Figure S15.**
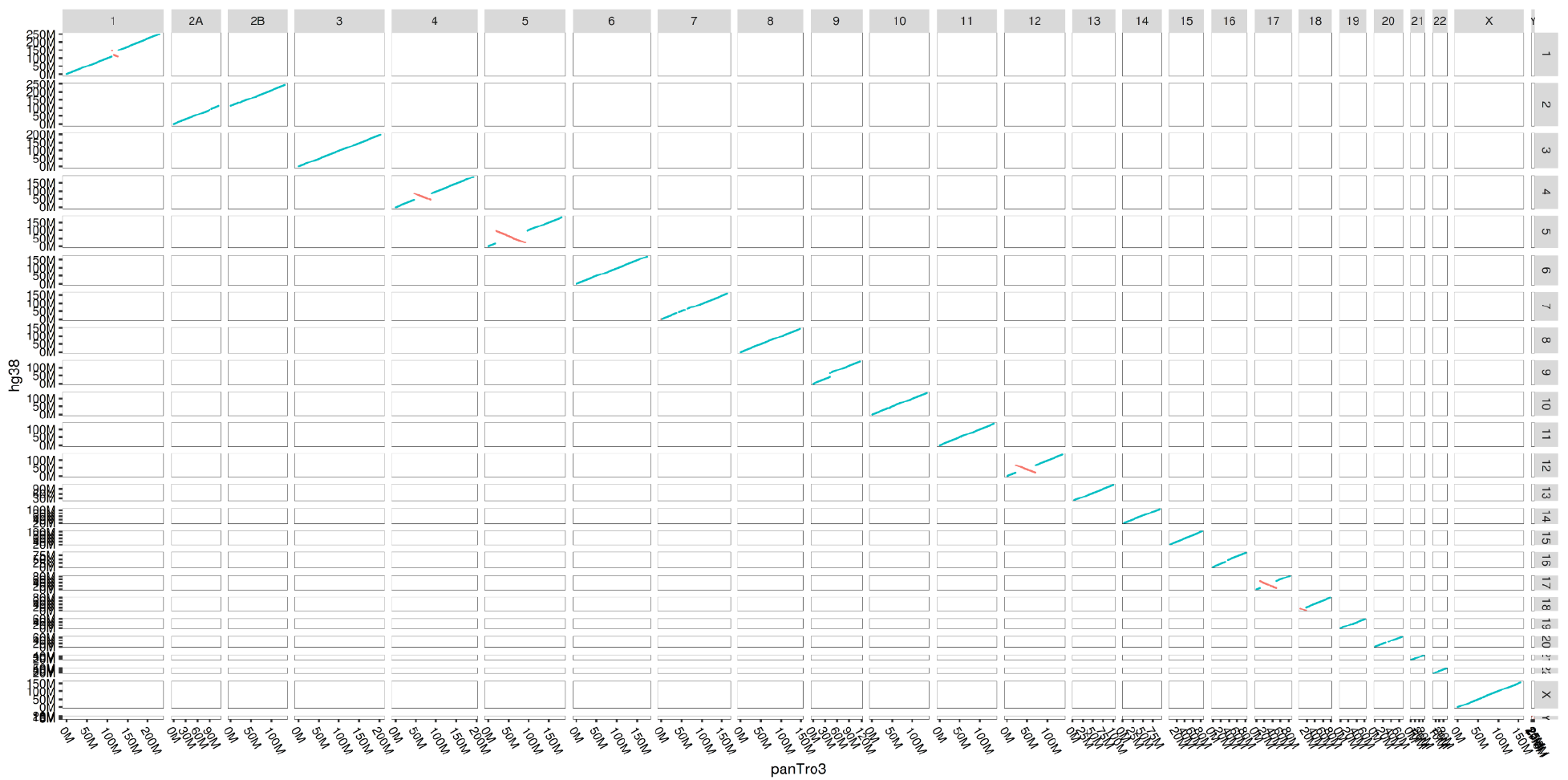
The plot of identified collinear anchors between human (hg38) genome assembly and chimpanzee (proTro3) genome assembly. Each dot was plotted by the start position on the reference (hg38) genome and the query genome (proTro3) of an anchor. Blue points represent anchors on the same strand between the reference genome and the query genome. Red points represent anchors on the different strands between the reference genome and the query genome.

We downloaded the widely used chain and net pipeline genome alignment of panTro3 against hg38 and transformed the alignment into bam format using the following commands:

~~~
wget http://hgdownload.cse.ucsc.edu/goldenpath/hg38/vsPanTro3/hg38.panTro3.synNet.maf.gz
gunzip hg38.panTro3.synNet.maf.gz
maf-convert sam hg38.panTro3.synNet.maf | sed ‘s/hg38.chr//g’ | sed ‘s/panTro3.chr//g’ > hg38.panTro3.synNet.sam
cat hg38.panTro3.synNet.sam | awk ‘$1 != “Y” && $3 != “Y” {print $0}’ > hg38.panTro3.synNet_noY.sam
cat hg38.panTro3.synNet_noY.sam | samtools view -O BAM --reference hg38.fa - | samtools sort - > hg38.panTro3.synNet_noY.bam
~~~

We used the following commands to calculate the number of aligned sites and position match sites for the human genome:

~~~
samtools depth 2panTro3_noY.bam | wc -l
samtools depth 2panTro3_noY.bam | awk ‘$3>0 {print $0}’ | wc -l
samtools depth hg38.panTro3.synNet_noY.bam | wc -l
samtools depth hg38.panTro3.synNet_noY.bam |awk ‘$3>0 {print $0}’ | wc -l
~~~

On the autosomes and the X chromosome, 2.902Gbp (95.73%) of the human genome is aligned using AnchorWave, this is larger than that of chain and net (88.97%, 2.697Gbp) (Figure S16). The number of sites aligned as position matches is comparable (AnchorWave:2.665Gbp, chain and net: 2.688Gb). The proportion of the genome identical from position match sites between the AnchorWave alignment and the downloaded chain and net alignment from UCSC website are very close to each other (98.5% from AnchorWave and 98.7% from chain and net alignment).

The position match ratio between the human and the chimpanzee genome is significantly higher than that between the maize line B73 and the maize line Mo17. Further, the DNA nucleotide identity ratio at position match sites between maize B73 and maize Mo17 (96.0% using AnchorWave) is lower than that between the human genome and the chimpanzee genome (98.7% using AnchorWave). This is consistent with the widely cited observation that maize lines are more different genetically than a human and a chimpanzee.

AnchorWave aligned 245.78 Mbp (97.13%) of candidate *cis*-regulatory elements as position match, which is slightly higher than the UCSC released chain and net alignment (244.11 Mbp, 96.59%).

**Figure S16.**
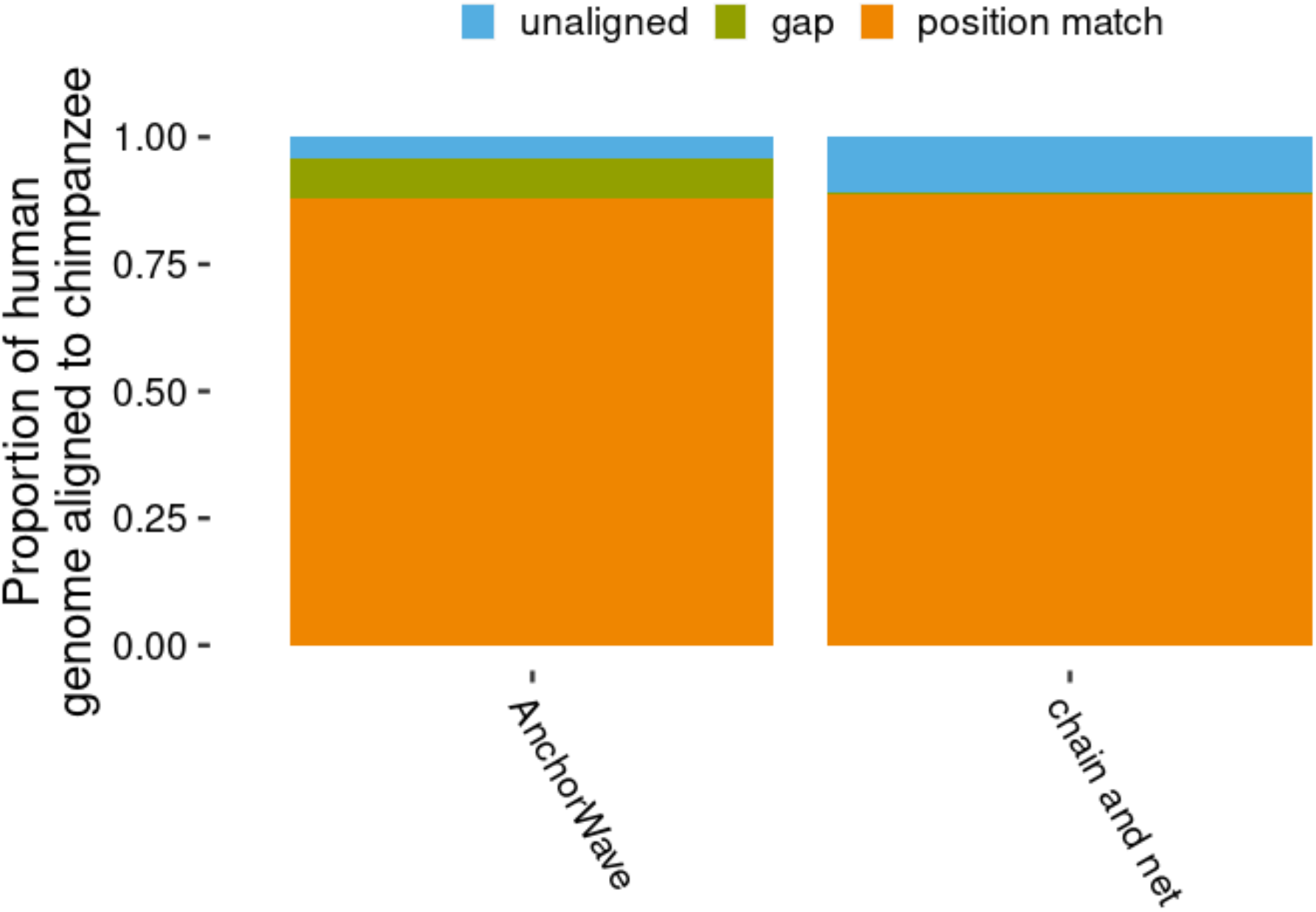
Comparison of the proportion of human (hg38) genome sites aligned to chimpanzee (panTro3) using AnchorWave versus genome chain and net genome alignment. The chain and net genome alignment is downloaded from http://hgdownload.cse.ucsc.edu/goldenpath/hg38/vsPanTro3/hg38.panTro3.synNet.maf.gz.

**Figure S17.**
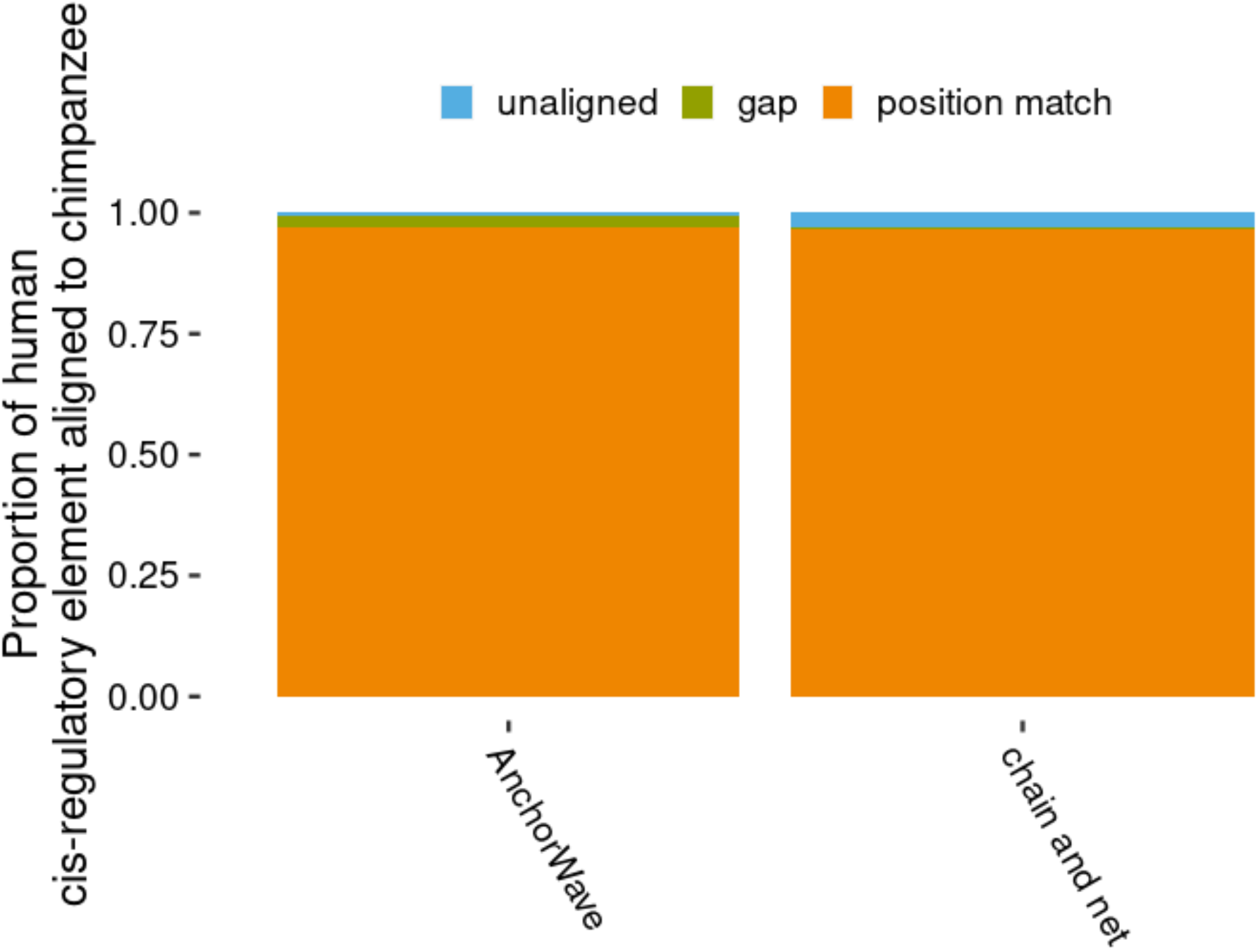
Comparison of the proportion of human (hg38) genome candidate *cis*-regulatory elements being aligned to chimpanzee (panTro3) using AnchorWave versus genome chain and net genome alignment.

## References

1. H. A. Lewin, et al., Earth BioGenome Project: Sequencing life for the future of life. Proc. Natl. Acad. Sci. U. S. A. 115, 4325–4333 (2018).

2. M. Exposito-Alonso, H.-G. Drost, H. A. Burbano, D. Weigel, The Earth BioGenome project: opportunities and challenges for plant genomics and conservation. Plant J. 102, 222–229 (2020).

3. B. Wei, et al., Genome-wide characterization of non-reference transposons in crops suggests non-random insertion. BMC Genomics 17, 536 (2016).

4. M. Freeling, M. J. Scanlon, J. E. Fowler, Fractionation and subfunctionalization following genome duplications: mechanisms that drive gene content and their consequences. Curr. Opin. Genet. Dev. 35, 110–118 (2015).

5. S. F. Altschul, W. Gish, W. Miller, E. W. Myers, D. J. Lipman, Basic local alignment search tool. J. Mol. Biol. 215, 403–410 (1990).

6. H. Li, N. Homer, A survey of sequence alignment algorithms for next-generation sequencing. Brief. Bioinform. 11, 473–483 (2010).

7. X. Tu, et al., Reconstructing the maize leaf regulatory network using ChIP-seq data of 104 transcription factors. Nat. Commun. 11, 5089 (2020).

8. R. C. O’Malley, et al., Cistrome and Epicistrome Features Shape the Regulatory DNA Landscape. Cell 165, 1280–1292 (2016).

9. B. Song, et al., Conserved noncoding sequences provide insights into regulatory sequence and loss of gene expression in maize. Genome Res. (2021) https://doi.org/10.1101/gr.266528.120.

10. J. L. Bennetzen, J. Ma, K. M. Devos, Mechanisms of recent genome size variation in flowering plants. Ann. Bot. 95, 127–132 (2005).

11. S. M. Kiełbasa, R. Wan, K. Sato, P. Horton, M. C. Frith, Adaptive seeds tame genomic sequence comparison. Genome Res. 21, 487–493 (2011).

12. Z. Li, et al., Multiple large-scale gene and genome duplications during the evolution of hexapods. Proc. Natl. Acad. Sci. U. S. A. 115, 4713–4718 (2018).

13. T. E. Wood, et al., The frequency of polyploid speciation in vascular plants. Proc. Natl. Acad. Sci. U. S. A. 106, 13875–13879 (2009).

14. H. Tang, et al., Screening synteny blocks in pairwise genome comparisons through integer programming. BMC Bioinformatics 12, 102 (2011).

15. D. Liu, M. Hunt, I. J. Tsai, Inferring synteny between genome assemblies: a systematic evaluation. BMC Bioinformatics 19, 26 (2018).

16. H. Li, Minimap2: pairwise alignment for nucleotide sequences. Bioinformatics 34, 3094–3100 (2018).

17. S. Marco-Sola, J. C. Moure, M. Moreto, A. Espinosa, Fast gap-affine pairwise alignment using the wavefront algorithm. Bioinformatics (2020) https:/doi.org/10.1093/bioinformatics/btaa777.

18. B. Song, Q. Sang, H. Pei, X. Gan, F. Wang, Complement genome annotation lift over using a weighted sequence alignment strategy. Front. Genet. 10, 1046 (2019).

19. X. Gan, et al., Multiple reference genomes and transcriptomes for Arabidopsis thaliana. Nature 477, 419–423 (2011).

20. G. Marçais, et al., MUMmer4: A fast and versatile genome alignment system. PLoS Comput. Biol. 14, e1005944 (2018).

21. H.-N. Lin, W.-L. Hsu, GSAlign: an efficient sequence alignment tool for intra-species genomes. BMC Genomics 21, 182 (2020).

22. Y. Jiao, et al., Improved maize reference genome with single-molecule technologies. Nature 546, 524–527 (2017).

23. S. Sun, et al., Extensive intraspecific gene order and gene structural variations between Mo17 and other maize genomes. Nat. Genet. 50, 1289–1295 (2018).

24. S. N. Anderson, et al., Transposable elements contribute to dynamic genome content in maize. Plant J. 100, 1052–1065 (2019).

25. R. J. Mroczek, J. R. Melo, A. C. Luce, E. N. Hiatt, R. K. Dawe, The maize Ab10 meiotic drive system maps to supernumerary sequences in a large complex haplotype. Genetics 174, 145–154 (2006).

26. Z. Chen, et al., De novo assembly of the goldfish (Carassius auratus) genome and the evolution of genes after whole-genome duplication. Sci Adv 5, eaav0547 (2019).

27. K. Howe, et al., The zebrafish reference genome sequence and its relationship to the human genome. Nature 496, 498–503 (2013).

28. Z. Swigonová, et al., Close split of sorghum and maize genome progenitors. Genome Res. 14, 1916–1923 (2004).

29. M. C. Stitzer, S. N. Anderson, N. M. Springer, J. Ross-Ibarra, The Genomic Ecosystem of Transposable Elements in Maize. bioRxiv, 559922 (2019).

30. M. B. Hufford, et al., De novo assembly, annotation, and comparative analysis of 26 diverse maize genomes. bioRxiv, 2021.01.14.426684 (2021).

31. E. T. Dermitzakis, A. G. Clark, Evolution of transcription factor binding sites in Mammalian gene regulatory regions: conservation and turnover. Mol. Biol. Evol. 19, 1114–1121 (2002).

32. M. R. McKain, et al., Ancestry of the two subgenomes of maize. bioRxiv, 352351 (2018).

33. J. C. Schnable, N. M. Springer, M. Freeling, Differentiation of the maize subgenomes by genome dominance and both ancient and ongoing gene loss. Proc. Natl. Acad. Sci. U. S. A. 108, 4069–4074 (2011).

34. D. Ellinghaus, S. Kurtz, U. Willhoeft, LTRharvest, an efficient and flexible software for de novo detection of LTR retrotransposons. BMC Bioinformatics 9, 18 (2008).

35. L. Kistler, et al., Multiproxy evidence highlights a complex evolutionary legacy of maize in South America. Science 362, 1309–1313 (2018).

36. H. Li, et al., The Sequence Alignment/Map format and SAMtools. Bioinformatics 25, 2078–2079 (2009).

37. D. Earl, et al., Alignathon: a competitive assessment of whole-genome alignment methods. Genome Res. 24, 2077–2089 (2014).

38. P. Lamesch, et al., The Arabidopsis Information Resource (TAIR): improved gene annotation and new tools. Nucleic Acids Res. 40, D1202–10 (2012).

39. P. Danecek, et al., The variant call format and VCFtools. Bioinformatics 27, 2156–2158 (2011).

40. J. Liu, et al., Gapless assembly of maize chromosomes using long-read technologies. Genome Biol. 21, 121 (2020).

41. N. Yang, et al., Genome assembly of a tropical maize inbred line provides insights into structural variation and crop improvement. Nat. Genet. 51, 1052–1059 (2019).

42. W. Shen, S. Le, Y. Li, F. Hu, SeqKit: A Cross-Platform and Ultrafast Toolkit for FASTA/Q File Manipulation. PLoS One 11, e0163962 (2016).

43. A. H. Paterson, et al., The Sorghum bicolor genome and the diversification of grasses. Nature 457, 551–556 (2009).

44. J. F. Hughes, et al., Chimpanzee and human Y chromosomes are remarkably divergent in structure and gene content. Nature 463, 536–539 (2010).

